# Optimized quantitative bacterial two-hybrid (qB2H) for protein-protein interaction assessment

**DOI:** 10.1101/2025.10.29.680610

**Authors:** Antoine Guyot, Emma Maillard, Kelly Ferreira-Pinto, Laure Plançon-Arnould, Aravindan A. Nadaradjane, Raphaël Guérois, Françoise Ochsenbein, Loïc Martin, Oscar H.P. Ramos

**Affiliations:** Sanofi Large Molecule Research, Vitry-sur-Seine, France; CEA, Département Médicaments et Technologies pour la Santé (DMTS), SIMoS; Université Paris-Saclay, 91191 Gif-sur-Yvette, France; Université Paris-Saclay, CEA, CNRS, Institute for Integrative Biology of the Cell (I2BC), 91198, Gif-sur-Yvette, France

## Abstract

Characterizing mutation effects on protein-protein interactions (PPIs) is crucial for elucidating protein structure and function. Massively parallel PPI variant analyses such as deep mutational scanning (DMS) enable interface identification and generate datasets for machine learning. In cellulo strategies such as two-hybrid systems provide straightforward access to such data, but reliability depends on quantitative properties. Here, we show that existing bacterial two-hybrid (B2H) systems have limitations constraining accurate dataset generation. We engineered and benchmarked optimized quantitative B2H (qB2H) alternatives, enabling strain-independent assays, improved metrics, and generation of high-quality datasets. We demonstrate qB2H utility through interface mapping and binder optimization. Perturbation analysis of single-site variants accurately recovered known ASF1 complex contact positions, matching crystallographic data. Integration of generative AI–based design yielded an ASF1-binding peptide with a 70-fold increase in affinity. qB2H offers to R&D scientists a robust, reusable platform for quantitative PPI analysis, enabling both rational protein engineering and data-driven discovery.

## INTRODUCTION

Protein-protein interactions (PPIs) are fundamental to biological processes, underpinning fields from biochemistry and physiology to drug discovery. Analyzing protein complexes through mutation effects yields invaluable insights by pinpointing interface positions, quantifying residue contributions to binding strength, and identifying hot spots. Incorporated into machine learning models, these mutational effect datasets guide molecular optimization^1–3^ across research, diagnostics, and therapeutic development with potential broad societal benefits. These applications rely on precise and reliable readouts. Beyond single point mutants, the possibility of simultaneously mutating two complex partners can provide important clues about compensatory effects and the basis of binding specificity. However, few robust, quantitative, and user-friendly technologies exist to screen for high-throughput simultaneous variations in two partners within the same assay^4^ (dual-partner DMS, dpDMS). While *in vitro* approaches allow investigating complex formation under arbitrary and well-defined conditions, they can be burdensome for achieving higher throughput. *In cellulo* approaches^5^ are often cost-effective, compatible with high-throughput screening and can be coupled with next-generation sequencing (NGS) for massive data analysis. Protein complementation assays (PCAs) and two-hybrid (2H) systems are among the approaches suitable for large-scale dpDMS screening^6,7^. While PCAs provide simpler and more direct readouts, two-hybrid systems offer greater flexibility in selecting the nature of the generated signal.

The concept of 2H was first introduced in yeast^8^ (Y2H) followed by bacteria^9^ (B2H) and mammalian cells^10^ (M2H). While Y2H and M2H support post-translational modifications, B2H is considered more resilient to false results^9^ for eukaryotic complexes and better fitted for fast screening of large libraries. Bacterial cells are also convenient for synthetic systems engineering^11–13^. Some bacterial two-hybrid (B2H) systems rely on the co-expression of two fusion proteins: the bait, which includes a DNA-binding domain (e.g., the λ cI repressor fused to one interaction partner), and the prey, which carries a transcriptional activation domain (e.g., the α [rpoA] or ω [rpoZ] subunit of bacterial RNA polymerase fused to the other partner). Interaction between these hybrid proteins recruits RNA polymerase to a synthetic promoter, thereby inducing the transcription of one or more reporter genes. Hochschild and colleagues pioneered such B2H setup in 1997^9^. The entire system included two plasmids (pBRα-rpoA and pAC-cI) and a strain holding a B2H-responsive promoter that controlled *lacZ* gene expression. Later, Ranganathan and colleagues^14^ proposed an alternative system implemented in three plasmids (pZA31-RNAα, pZS22 and pZE1RM) and used eGFP as reporter. Based on the characteristics of these B2H systems, we reasoned that Hochschild’s system would provide low signal output, while Ranganathan’s setup could impose a metabolic burden due to the number of plasmids (and antibiotics for plasmid selection).

Here, we report the engineering, characterization and application of a convenient and robust quantitative bacterial two-hybrid system (qB2H) designed to serve as a versatile tool for future protein engineering projects. First, we describe the engineering strategy by evaluating alternative versions of the pivotal B2H-responsive promoter within a harmonized reporter context to characterize their properties and find options for improving B2H robustness and quantitativeness (qB2H). Next, we validate the optimized systems in representative applications, including interface mapping and binder optimization. As a model system for qB2H engineering, benchmarking, and validation, we employed complexes between the histone chaperone ASF1 (Anti-silencing Function 1) and synthetic peptides whose dissociation constants (*K_d_*) had been previously determined by isothermal titration calorimetry (ITC), covering a range from undetectable binding (> 100 µM) to nanomolar affinities^15^. These peptides have been originally designed as first steps towards the development of anti-cancer compounds impairing the proliferation of cancer cell lines^15–17^.

## RESULTS

An optimized quantitative bacterial two-hybrid (qB2H) system can advance protein engineering and expand its integration with machine learning applications. To achieve this, it is essential to identify current limitations and enhance qB2H signal quantification in relation to the thermodynamic properties of specific protein interactions. In developing improved qB2H variants, our initial efforts focused on the key promoter responsive to complex formation between hybrid fusion proteins (rpoA–partner_A and cI–partner_B).

We then examined additional factors likely to influence system performance, such as the number of plasmids, the balance between cI and rpoA fusion expression, the presence of methylation sites that could increase transcriptional stochasticity, and the choice of reporter gene. Based on these factors, four versions of the qB2H system (versions 1–4) were constructed, with their key differences summarized in Fig. 1A.

**Figure 1:**
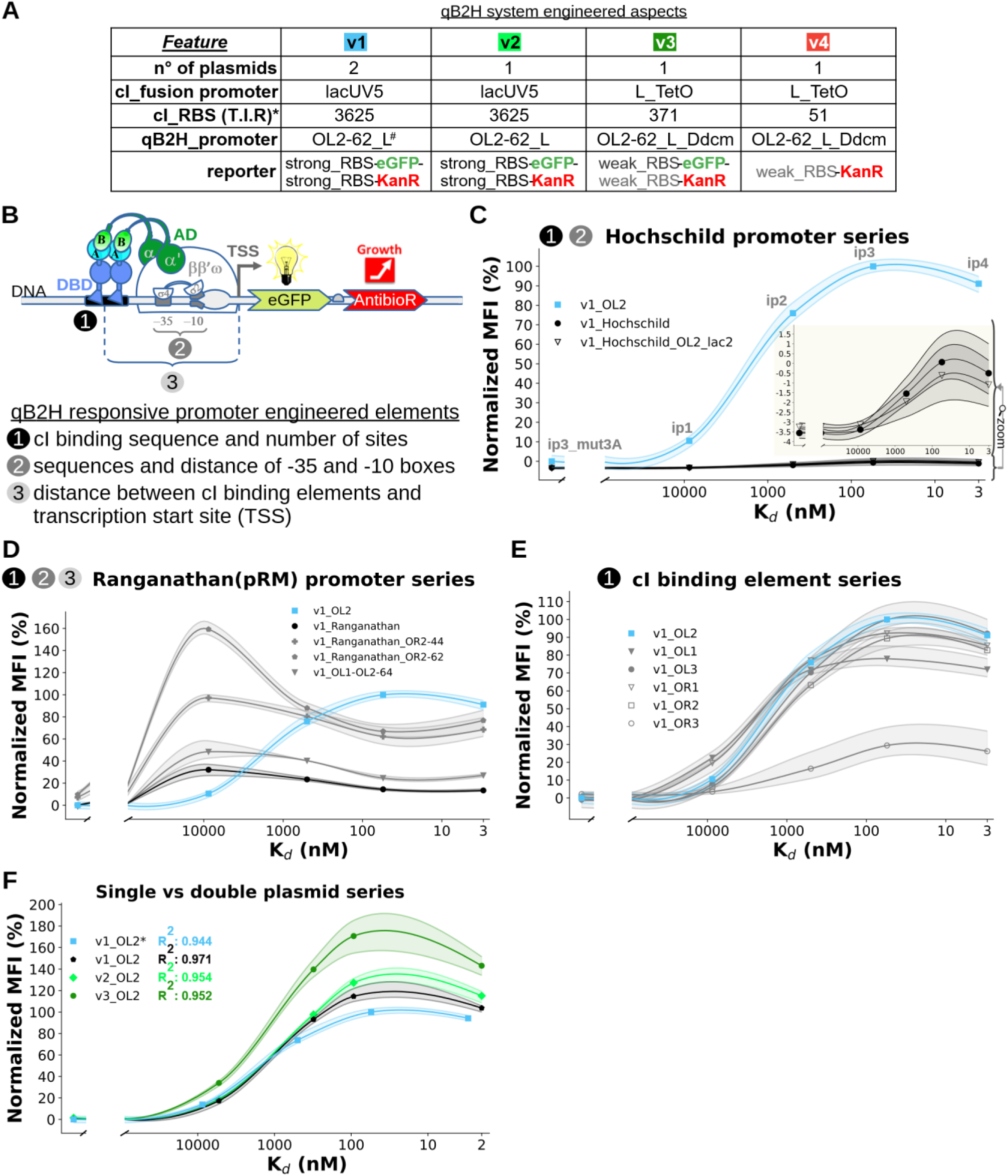
*B2H output signal as a function of K_d_ across various settings* (see Supplementary Table 2 for details on B2H- responsive promoters). A reference qB2H-responsive promoter, based on OL2 cI binding element and *pL* −35 and −10 boxes, is indicated by a blue line (**v1_OL2**, 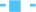) in panels C to F. (**A**) qB2H feature differences across versions (v1 to v4; details on Supplementary Fig. 1A – plasmid genealogy and 1B – plasmid maps). **\***T.I.R.: translation initiation rate-estimated using “Promoter Calculator” from “De Novo DNA” server (www.denovodna.com). ^#^Alternative promoters were evaluated in the context of v1 series plasmids, but OL2-62_L was assumed as reference. (**B**) Schematic qB2H reporter gene showing promoter structure and engineered elements (1, 2, 3). (**C**) Comparison of, double plasmid systems (v1) including **v1_OL2**, the promoter originally proposed by Hochschild and colleagues^9^ (Hoschild; **-**λ**-**) and an alternative version (Hoschild_OL2_lac2; **-▽-**). (**D**) Comparison of **v1_OL2** with promoters harboring 2 cI binding elements: v1_Ranganathan (*pRM*\*; originally employed by Ranganathan and colleagues^14^ with functional OR1 and OR2 elements; **-●-**); v1_Ranganathan_OR2-44 (*pRM* * variant where OR2 was displaced at −44 from TSS; −**+**-); v1_Ranganathan_OR2-62 (*pRM* * variant where OR2 was displaced at −62 from TSS; **-**⬟**-**) and; v1_OL1-OL2-64 (similar to v1_Ranganathan_OR2-62 but harboring OL1 and OL2 elements, the latter is displaced at −64 from TSS and −35 and − 10 boxes derives from lambda L promoter, *pL*; **-**▾**-**). (**E**) Comparison of B2H-responsive promoters harboring only one of the natural cI binding elements (OR1, OR2, OR3, OL1, OL2 or OL3) at position −62 from the transcription start (TSS) of *pL* using two-plasmid system (v1). (**F**) Comparison of signal output (Normalized MFI) from double (v1, ASF1A: 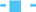, ASF1B: **-**⬟**-**) or single-plasmid series (v2, v3; respectively 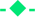 and 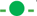) on pOL2-62_L output. v1_OL2* is the same reference version as other panels. qB2H versions without “*” represent system where ASF1B_N_ partner was used instead of ASF1A_N_ (used in reference, blue line). The confidence interval (ci = 95%) was represented by semi-transparent regions and results from three independent experiments (data available in Supplementary Table 3).

### qB2H engineering

To guide the engineering of the qB2H system, we used an eGFP reporter alongside a set of peptides with varying affinities for the N-terminal domain of ASF1A (UNIPROT: Q9Y294), ASF1A_N_, as shown by their interaction strengths^15^ (ip3_mut3A: > 100,000 nM; ip1: 8,700 nM; ip2: 500 nM; ip3: 55 nM; and ip4: 3 nM; Supplementary Table 1). In our first experiments, we employed a two-plasmid system (v1) to examine the effects of interactions on a set of 12 two-hybrid responsive promoters (Supplementary Table 2). The main goal was to explore two key features of the qB2H system: (i) the signal-to-noise ratio, and (ii) the correlation between the signal outputs and the binding affinities. Aiming to assess their impact on these features, we compared reported promoters to engineered alternatives by targeting specific functional regions. The promoters harbor one or more cI binding sites, with different binding affinities. Furthermore, the promoters included specific combinations of −35 and −10 boxes corresponding to different transcription strengths: *pLac* (weak), *pRM* and *pL* (strong). A bicistronic construction allowing signal output as fluorescence and kanamycin resistance, eGFP-KanR, was chosen as reporter. A schematic representation of these elements is provided in Fig. 1B.

The promoter described by Hochschild and colleagues, along with a closely related alternative, shows reasonable coherence between signal intensities (mean fluorescence intensity, MFI) and complex affinities (ip3_mut3A to ip4) but exhibits a low signal-to-noise ratio (2.36 to 3.23) and large confidence intervals (Fig. 1C). In contrast, promoters such as those proposed by Ranganathan and colleagues^14^ (*pRM*), which contain two cI binding sites, provide an acceptable signal-to-noise ratio (6.73 to 10.95) but demonstrate poorer consistency with complex affinities (R²) and even show an inverse correlation (Fig. 1D, Supplementary Table 3). Thus, both sets limit the ability to make precise quantitative estimates of interaction strength based on output signals. Overall, these results indicate that signals are higher and more coherent with interaction strength when only one cI binding site is positioned near the –62 location from the transcription start site (TSS) and coupled with a strong promoter.

Next, we analyzed the system’s sensibility cI operator sites, using promoters with affinities decreasing in the following order: OL1 > OR1 > OL3 > OL2 > OR2 > OR3^18,19^. Interestingly, we observed the highest signals for intermediate-affinity interactions between cI and its DNA binding sites (OL2 and OL3). In contrast, the lowest-affinity (OR3) and highest-affinity (OL1) interactions produced the lowest and second-lowest maximum signals, respectively (Fig. 1E). These results highlight the complex dynamics underlying B2H activity. Except for OR3 (*R*^2^ = 0.81, *n* = 4 measurable *K_d_* and fluorescent signal coordinates), all cI binding elements demonstrate a strong linear correlation between affinity and signal output for ASF1 — peptide interactions ranging from about tens of micromolar to nanomolar (*n* = 4, *R*^2^ between 0.935 and 0.999; Supplementary Table 3). Again, excluding OR3, undetectable interactions (> 100 µM; ASF1-ip3_mut3A) yield significantly lower outputs (n = 3 experiments, *p* < 0.05, *t*-test; Supplementary Table 3) than weak interactions (∼10 µM; ASF1 — ip1), suggesting that very low affinity interactions can be distinguished from background. However, even if often not statistically significant, the signal intensities for the strongest interaction (nanomolar affinity for ASF1A — ip4) appear lower than the slightly weaker interaction (tens of nanomolar affinity for ASF1A — ip3). An explanation is that the eGFP overexpression, which can lead to H_2_O_2_ accumulation^20^ and increases metabolic burden^21^, may result in counterselection against highly expressing clones.

A second set of experiments was conducted to improve the system’s quantitative properties. We focused on further improvements in signal-to-noise ratio and on reducing the system stochasticity. For that purpose, we merged all the required B2H elements into a single plasmid, improved cI and rpoA expression ratio, suppressed a methylation site (dcm) in the qB2H-responsive promoter, and simplified the B2H reporter gene. Furthermore, we also searched to reduce the required antibiotic concentration during selection experiments and explored the effect of a paralog of ASF1A, ASF1B (UNIPROT: Q9NVP2), on signal-to-noise ratio.

ASF1A_N_ and ASF1B_N_ exhibit similar response profiles, single-plasmid are comparable to two-plasmid systems and the latest qB2H implementation, v3, reaches the highest signal-to-noise ratio (Fig. 1F). Like for other tested versions, the signal values (MFI) can be almost entirely explained by *K_d_* variations (*n* = 4, *R*^2^ = 0.952). Furthermore, single-plasmid vectors are among the most reproducible systems based on fluorescence signals (the mean coefficient of variation, CV, among experiences for v2 and v3 series are respectively 3.67% and 7.04%). Comparing Hochschild’s and Ranganathan’s promoters to the v3 series, the signal-to-noise ratio (ASF1 — ip3 by ASF1 — ip3_mut3A) increases from 2.36–6.73 to 27.49, while the correlation (*n* = 4, *R²*) between fluorescent signal and *K_d_* improves from 0.783–0.786 to 0.952 (Supplementary Table 3).

Since most *Escherichia coli* strains express lac repressor (LacI), co-transformation with v1 (pAC-cI_E34P and pBRα) or transformation with v2 plasmid series allows for both, strain independent selection based on fluorescence (using fluorescence activated cell sorting, FACS) and/or kanamycin resistance. However, since the KanR gene expression was not optimized in these systems, the required kanamycin concentration can be remarkably high for strong interactions, especially for v2 plasmid series (e.g., ≥ 1 mg/mL for interactions such as ASF1B — ip3). The v3 plasmid series cumulates additional modifications that lead to higher signal-to-noise ratio, requires lower antibiotic concentrations and can reduce experimental stochasticity. These modifications include the invalidation of Dcm site close to the −35 box of the qB2H-responsive promoter and a stringent control of λ cI_E34P fusions from a strong and tightly regulated promoter (pLTetO promoter)^22^ coupled to a weak RBS. Indeed, earlier work demonstrated that this kind of setup can reduce stochasticity^23^. Single-plasmid from v4 series incorporate all the latter features, but they were also engineered to provide lower and monocistronic expression of KanR that reduce the system’s complexity and improves quantitative measures.

### Optimization for high-throughput experiments under antibiotic selection

Applying an appropriate selection pressure to a cell population, combined with NGS, allows estimation of genotype frequencies over time and calculation of their enrichment or depletion, which can be interpreted as fitness under the applied conditions. In this approach, antibiotics provide a convenient and effective means of imposing selection pressure.

Single-plasmid vectors linking PPI strength to kanamycin resistance—either as a single reporter (v4: KanR) or a dual reporter (v3: eGFP-KanR)—were compared under different conditions in triplicate. MIC_50_ estimations (see “Methods”) were employed to evaluate the effects of initial cell density (OD₆₀₀ = 0.05 or 0.01) and selection duration (4 h or 6 h) on kanamycin resistance for each ASF1–peptide interaction (Fig. 2A, left panel).

**Figure 2:**
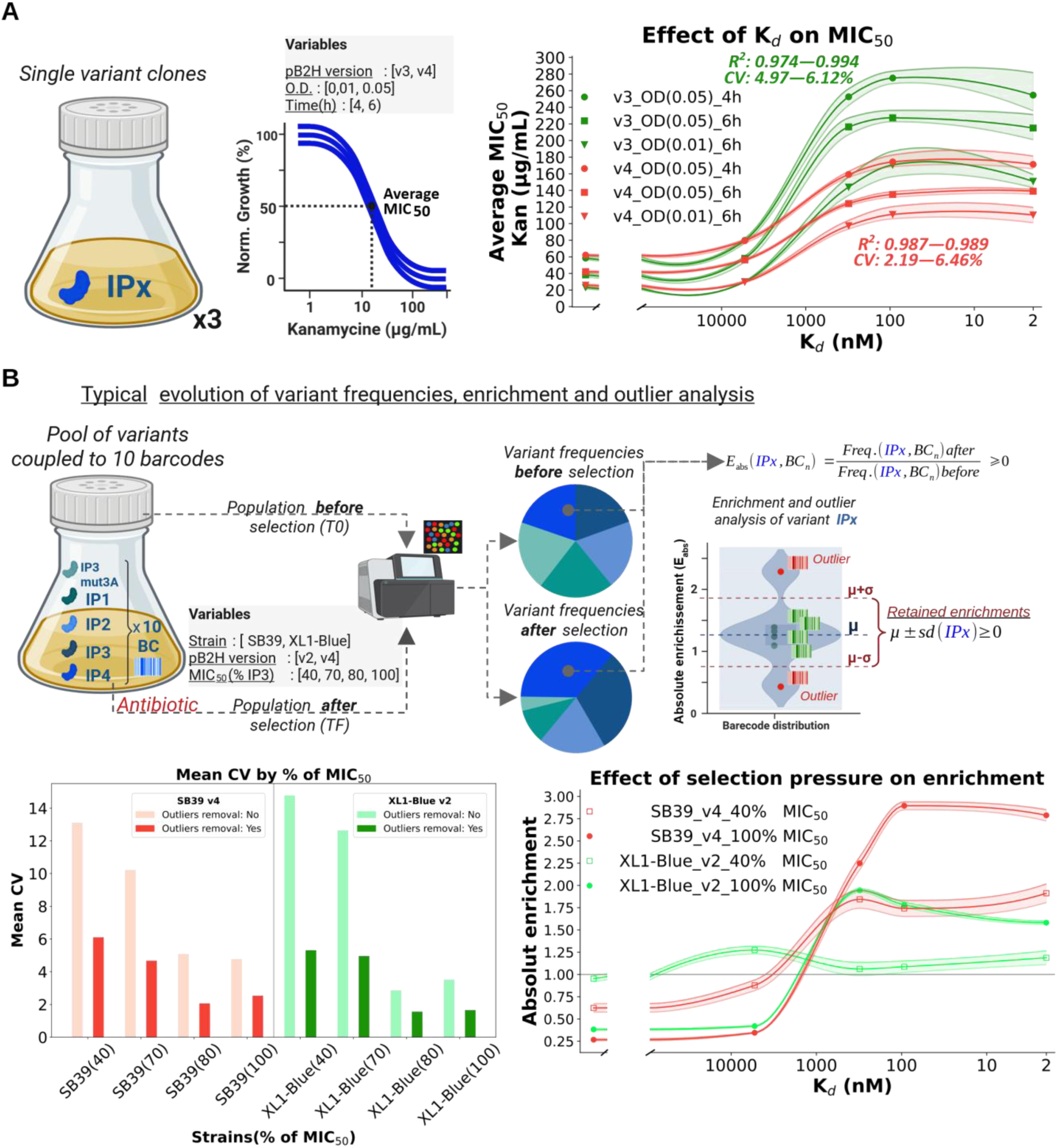
Effect of kanamycin on single or multiple variant clones’ growth. (**A**) <u>Left</u>: Schematic representation of the estimated average MIC_50_ for each affinity (ASF1 — ip_x) at varying conditions. <u>Right</u>: Antibiotic resistance (the average MIC_50_) versus *K_d_* plot. The experimental conditions are indicated by legend names composed of 3 fields separated by “_”, respectively: plasmid version, starting OD and time point (h) after kanamycin addition. The confidence interval (ci = 95%) was represented by semi-transparent areas. The results represent three independent experiments. Triplicate MIC_50_ curves for each *K_d_* are provided as supplementary information (Figures 2 to 7, Supplementary Table 4**). (B**) Comparison of non-optimized and optimized systems. <u>Top</u>: Schematic representation of the experiment designed to assess the behavior of optimized and unoptimized systems. Each peptide was coupled to 10 barcodes to estimate their independent frequencies and absolute enrichment (E_abs_), identify outliers and evaluate stochasticity (CV, coefficient of variation). The right side of the panel represents calculation of E_abs_ for each peptide (ip_x) and barcode (Bc_n_) from their frequencies before and after application of selection pressure (kanamycin supplementation). Enrichment values that are far from average (outliers, red) can be removed using z-score method to improve standard deviation (sd). <u>Bottom left</u>: Mean coefficient of variation (CV) by B2H system (SB39 v4, reddish; XL1-Blue v2, greenish) and selection pressure (40, 70, 80, 100% of MIC_50_ for ASF1B_N_ — ip3 interaction). Light colors indicate no outliers’ removal, while dark colors indicate the opposite. Conditions that are represented in the bottom right panel are surrounded by black lines. <u>Bottom right</u>: Enrichment profile over the studied *K_d_*range for SB39 and XL1-Blue strains under the lowest (40% MIC_50_) or highest (100% MIC_50_) selection pressure (without outliers’ removal). The experimental conditions are indicated by legend names composed of 3 fields separated by “_”, respectively: strain, plasmid version, kanamycin concentration (as percentage of ASF1— ip3 MIC_50_). Enrichment of 1 (dashed gray line) corresponds to invariable frequency in the population before and after kanamycin selection. Raw data available as supplementary information (Supplementary Table 5).

Fig. 2A (right panel) shows changes in average MIC_50_ under these conditions. Consistent with earlier studies^24,25^, we observed lower MIC_50_ values (higher antibiotic sensitivity) for lower starting cell density and longer selection times, showing that lower antibiotic concentrations can be used under these conditions. Compared to the single-gene reporter (v4), the bicistronic reporter (v3) exhibits higher MIC_50_ values, a pronounced signal drop for the strongest affinity interaction (ASF1B— ip4), and wider confidence intervals, reflecting higher stochasticity. Consequently, owing to its superior MIC_50_–*K_d_* consistency and lower data variability, the v4 system is considered the most suitable for quantitative analysis of interactions. It is also a more economical and environmentally friendly option, as it requires lower antibiotic concentrations.

The results indicate that lower starting OD and longer selection times provide better signal-to-noise ratios (MIC_50_ ASF1 — ip3 by MIC_50_ ASF1 — ip3_mut3A), better coherence between antibiotic resistance (MIC_50_) and *K_d_* values (*n* = 4 *K_d_* values ranging from 10 µM to nM, *R*^2^ > 0.98) and reproducible results (mean CV = 2.19-6.46%; Supplementary Table 4). Across all experimental conditions, no statistically significant differences were observed for MIC_50_ values corresponding to *K_d_* values of 93 nM and 2 nM, suggesting the presence of a plateau in the nanomolar to tens-of-nanomolar range. The lack of a signal drop at the highest affinity interaction (ASF1 — ip4; *K_d_* = 2 nM) is likely due to the absence of the eGFP reporter and the dynamic kinetics of complex formation among cI — DNA, ASF1 — ip4, and the RNA polymerase transcription machinery. Finally, SB39 v4, starting OD_600_ = 0.05 and selection over 6 hours was the only condition where significant difference was observed between the *K_d_*values of 310 nM and 2 nM (*n* = 3 experiments, *p* > 0.05, *t*-test).

Mapping these antibiotic sensitivity landscapes helped define the optimal antibiotic concentration range for subsequent stochasticity analyses and benchmarking of the optimized systems (Fig. 2B).

To evaluate the impact of our latest design optimizations on system performance and stochasticity, we compared v2 in the XL1-Blue strain (the B2H chassis used in the commercial BacterioMatch II kit, Agilent) with the v4 series in SB39 (our engineered strain). We focused on two key properties relevant to qB2H systems for PPI quantification: (i) the effect of selection pressure on stochasticity, using the coefficient of variation (CV) as a proxy, and (ii) the correlation between enrichment values and interaction strength. CVs were calculated as the percentage of enrichment standard deviation (from 10 unique DNA barcodes for each ASF1 peptide binder) compared to the mean enrichment. We also evaluated the effect of outliers’ removal on data discrepancy (Fig. 2B, top). Fig. 2B (bottom, left) shows the stochastic profiles (mean CV) by B2H system and selection pressure. As expected, the results show that outliers’ removal using a z-score threshold considerably reduces data dispersion, particularly under conditions of low selection pressure. Furthermore, this approach does not markedly change mean enrichment values (Supplementary Table 5). Fig. 2B, bottom, right, shows the enrichment profile over the tested range of affinities for selection pressures corresponding to 40 or 100% MIC_50_ of the ASF1B — ip3 interaction in each system. The SB39 v4 showed a more coherence between enrichment and affinity compared to XL1-Blue v2 under both selection pressures. The results also indicate that higher selection pressures enhance the predictive linkage between enrichment by *K_d_* (particularly for stronger interactions) and reduce stochasticity. While CV values were comparable among B2H systems, only SB39 v4 displayed the expected correlation between enrichment and affinities. Indeed, strong change in *K_d_* vs enrichment coherence is observed between the worst condition (XL1-Blue v2, 40% MIC_50_, without outliers’ removal condition; *R*^2^ = 0.082) and the best (SB39 v4, 70% MIC_50_, with outliers’ removal; *R*^2^=0.995). Concerning SB39 v4, all conditions provide good coherence (*n* = 4 *K_d_*s; *R*^2^ > 0.96, Supplementary Table 5). Therefore, experimental conditions such as B2H system and selection pressure should be carefully considered for best results.

Taken together, we found that the SB39 strain, v4 and high selection pressure (70-150% MIC_50_of a strong interaction for a given couple of partners) yield the best quantitative results. In experiments where different barcodes can be assigned to each variant, the removal of outliers can also be applied. Concerning the latter aspect, narrower z-scores around the mean (e.g., 0.5 or 0.25) can further reduce standard deviation. This property can be advantageous to increase the resolution during perturbation scores calculations (see the “Interface mapping” section).

SB39 strain represents the convergence of some key features that make it very convenient for qB2H experiments: i) fast growth (handy for performing batch experiments over a working day); ii) presence of chromosome gene copy of some of the most used transcription repressors (*phlF*^AM^, *cymR*^AM^, *luxR*, *vanR*^AM^, *lacI^A^*^M^, *tetR*) for genetic circuit design and regulation (suitable for future engineering of biological circuits); iii) deficiency in Lon and OmpT proteases that improves protein stability; iv) *endA* and *recA* mutations that improve plasmid transformation and stability, making it compliant with library construction and screening. Furthermore, this strain shows elevated electroporation efficiency (>1×10^9^ cfu/µg of pUC19) using in house preparations^26^.

### Interface mapping

Using the plasmids developed for qB2H, we next evaluated the method’s reliability for mapping the interaction surface between ASF1 and binding partners whose interaction have also been characterized by X-ray crystallography. Positional single-mutant libraries of ASF1 were obtained from Twist Bioscience and cloned into the pB2H v4 series (details in Methods and Supplementary Information: ‘Construction and screening of interface mapping libraries,’ Tables 1, 2, and 7). The number of CFUs obtained exceeded 100-fold coverage of the theoretical nucleotide diversity.

Variant frequencies in transformed cell populations were compared before and after antibiotic selection by NGS (Illumina NovaSeq X). Data analysis with the mave2imap pipeline enabled calculation of normalized enrichment values for each variant (Fig. 3A, Supplementary Table 8) and assignment of a perturbation score to each assessed position (Fig. 3B, Supplementary Table 9). Since enrichment directly reflects binding strength under selection pressure (Fig. 2, v4) and mutations at interface positions are more likely to disrupt complex formation, interface residues can be inferred by identifying the most perturbed positions. The positions with the highest perturbation scores mapped onto ASF1’s structure overlaps with the interaction interfaces observed in crystallographic complexes of two different binders (ip3 and HIRA - Histone Regulatory Homolog A; Fig. 3C and 3D). Variant fitness data, normalized to the wild-type enrichment, are provided in Supplementary Table 8-9 and Supplementary Fig. 9-10.

**Figure 3:**
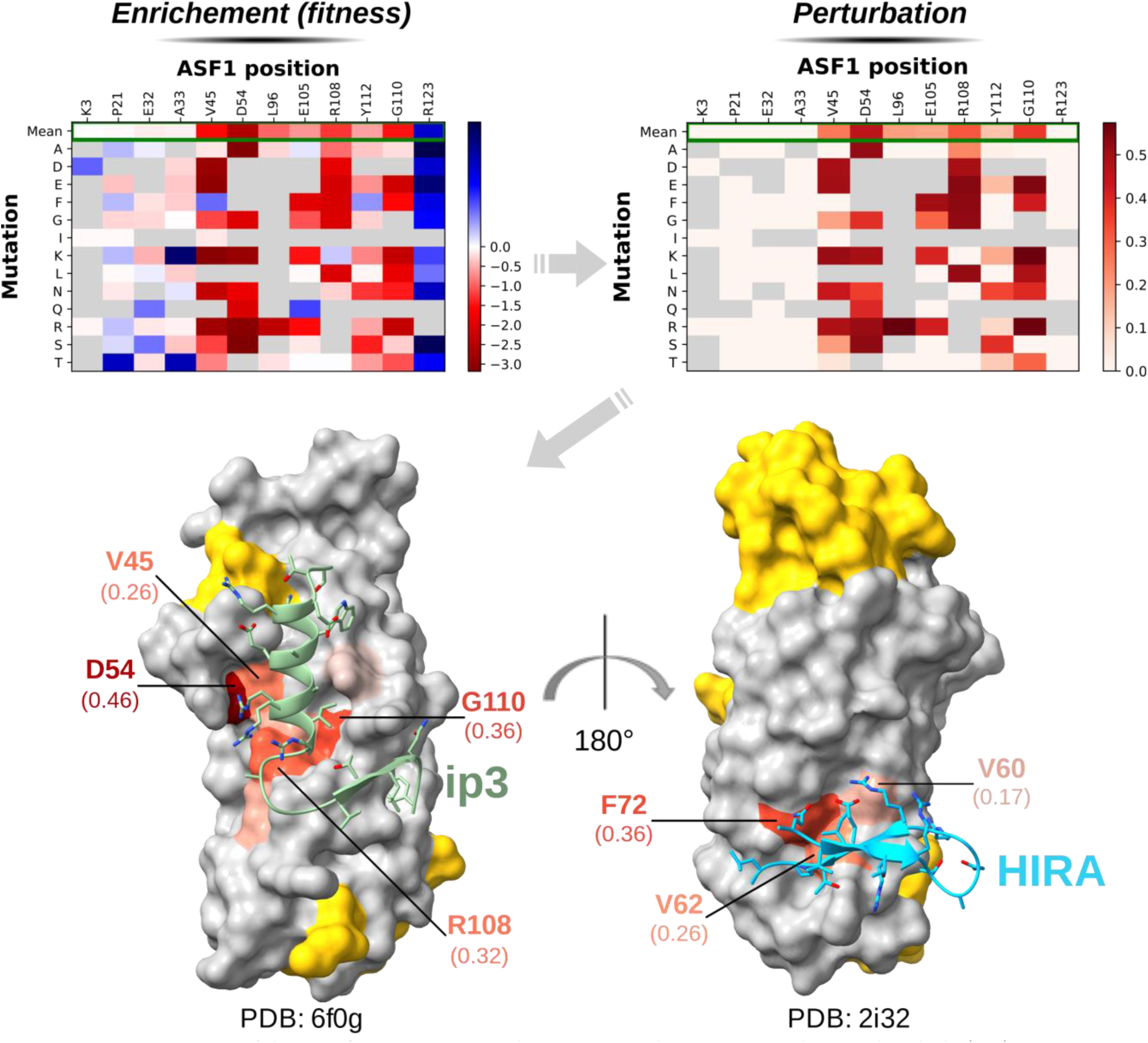
Demonstration of the interface mapping application. Perturbed positions above a threshold (0.1) were mapped onto the target’s (ASF1) surface and colored as a gradient from light orange to red based on their perturbation scores. Non-perturbed positions are shown in gray. Some positions of the target were not assessed (yellow) since the exploration of 2 regions for each partner already held sufficient positions that could be assumed as perturbation controls (not found at the interface) to be compared to interface positions. (**A**) Analysis of NGS data from the population before and after selection allows obtaining normalized enrichment for each variant (compared to WT). Representative positions are displayed as a heatmap, colored based on their log2 normalized enrichment. (**B**) Since mutation of interface residues are more likely to negatively affect interaction, enrichment scores are used to calculate perturbation scores (loss-of-interaction scores). Representative positions are shown (same as in panel A) as a heatmap, colored based on perturbation scores. (**C**) Interface mapping obtained for the ip3 binder (green). (**D**) Interface mapping obtained for the HIRA binder (light blue). In (C) and (D) the top most perturbed positions are labeled with their names and perturbation scores (in parentheses).

Overall, our results demonstrate the applicability of the qB2H v4 system for interface mapping, as well as for identifying interaction hotspots.

### Binders’ screening

AI-based approaches, such as those implemented in BindCraft^27^, represent major advances for *in silico* binder design and provide sets of solutions to be experimentally evaluated. We tested whether our qB2H v4 system can select an optimal solution among a set of generated models. Using a generative AI pipeline (by coupling of RFdiffusion^28^, ProteinMPNN^29^ and AlphaFold2^30–32^) we have generated and selected hundreds of N-terminal extensions of the ip4 peptide to be screened by qB2H (pB2H v4 series). These designs were coupled to 3 linkers of variable lengths (GSK, GSEK, GSEAK). This dataset also includes a restricted number of manually designed variants based on structural inspection of the modeled interface RFdiffusion (so-called ‘rational designs’) and some negative controls. These variable extensions were fused to a constant peptide, ip4_mutG (a weak ASF1B_N_ binder; *K_d_* = 10.6 µM), which contains a mutation to lower the affinity for ASF1, thereby enabling selection of extensions that enhance interaction strength within the dynamic range of the qB2H system (from tens of µM to nM). As a result, the library encompasses 1,134 peptides composed by a variable and a constant region with global lengths ranging from 34 to 41 amino acids (Fig. 4A). The library was subjected to 2 different selection conditions: i) short-term (4–5 hours) batch selection using increasing kanamycin concentrations (0, 174, 261, 348 µg/mL kanamycin), and ii) long-term (24, 32 hours) selection in a turbidostat with fixed kanamycin concentration (209 µg/mL kanamycin). Samples collected before (T0) and after selection (TF) were compared to evaluate the impact of different selection pressures on population outcomes.

**Figure 4:**
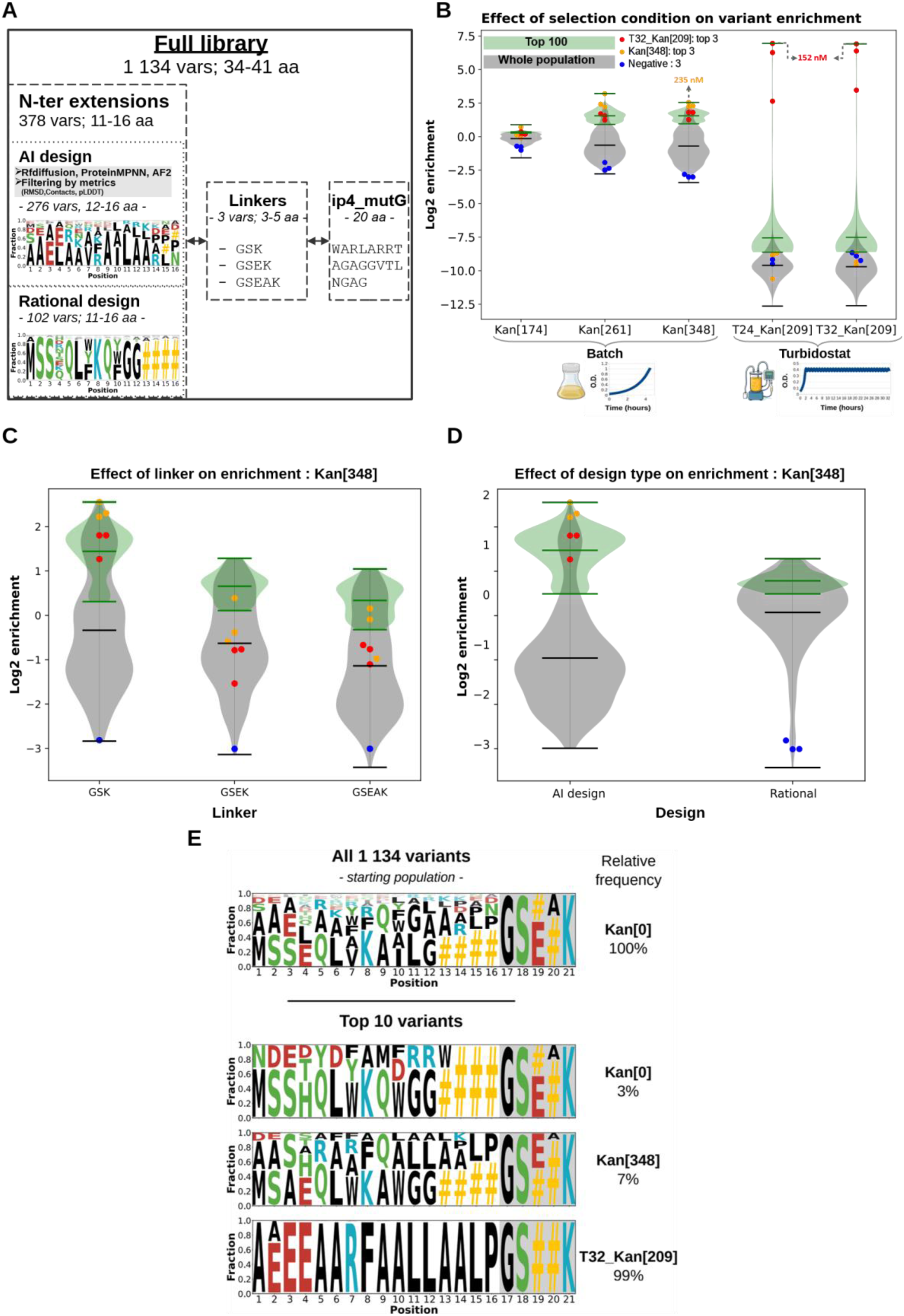
Binder selection experiment. (**A**) Library structure (extension-linker-ip4mutg), composition and diversity. (**B**) Impact of selection condition on variant enrichment. Violin plots show the enrichment of all variants (gray) or top 100 variants (green) under different kanamycin concentrations in short-term (batch; Kan[174], Kan[261] and Kan[348]) or the long-term experiments (turbidostat; T24_Kan[209] and T32_Kan[209]). (**C**) Effect of linker on short-term (batch, Kan[348]) selection experiments. (**D**) Effect of design approach (AI or Rational) on short-term (batch, Kan[348]) selection experiments. (**E**) Weblogo representing the fraction of amino acids residues by position for the entire population before selection (top of the panel) and for the top 10 most frequent variants under different conditions (Kan[0], Kan[348] and T32_Kan[209]). Relative frequencies of the selected number of variants are shown (right) as percentage for each condition.

As expected, stronger selection pressures lead to greater shifts in variant frequencies within the population (enrichment or depletion), reflecting their relative fitness. In batch experiments, variant frequencies under 174 µg/mL kanamycin correlated more strongly to those of the non-selected population (*n* = 734, *r* = 0.96, *p* = 0; Pearson correlation coefficient) than with those under stronger selection pressures, 261 µg/mL (*n* = 734, *r* = 0.76, *p* = 3.03 × 10⁻¹⁶³; Pearson correlation coefficient) or 348 µg/mL (*n* = 734, *r* = 0.76; *p* = 1 × 10⁻¹⁶²; Pearson correlation coefficient). Selection at 261 and 348 µg/mL produced nearly identical frequency profiles (*n* = 734, *r* = 0.99; *p* = 0; Pearson correlation coefficient), indicating that increasing the concentration from 150% to 200% of the ASF1B_N_ — ip3 MIC_50_ did not further affect variant frequencies. In long-term turbidostat experiments (24 or 32 h at constant O.D.), variant frequencies were more correlated to those observed at higher kanamycin concentrations in batch culture than to those under lower or no selection. Moreover, continuous cultures at 24h and 32h displayed almost identical profiles (*n* = 734, *r* = 0.99; *p* = 0), indicating that 24h of selection is sufficient to produce strong shifts in variant frequencies compared to no selection (*n* = 734, *r* = 0.24; *p* < 3.12 × 10⁻¹¹; Pearson correlation coefficient). High correlation coefficients (*r*) across similar conditions serve as pseudo-replicates, confirming the system’s reproducibility over large libraries. A detailed comparison among conditions is provided as supporting information (Supplementary Figures 11–22, Supplementary Table 10). Supplementary Figures 17 and 18 provide a general overview of the phylogenetic relationships and fitness landscape of the screened library.

These results suggest that 4–5 h of selection at 150% of the MIC₅₀ of a strong interaction provides a suitable starting point for short-term batch experiments, while 24h of selection at 120% MIC₅₀ can be employed for long-term experiments. Short-term experiments are preferable when the goal is to generate comprehensive variant datasets, while long-term experiments exert more drastic effects on population composition, which may be helpful for selection campaigns aiming to isolate the strongest binders. In other words, long-term selection drives extreme enrichment and depletion, resulting in convergence of the population toward high fitness phenotypes. Indeed, in long-term experiments, the first top 2 enriched variants accounted for about 89% of the population (about 66% and 23%; Supplementary Table 10), showing strong and reproducible convergence toward closely related solutions, differing by only one amino acid substitution, and sharing the same linker sequence (GSK). Under these conditions, the bulk of remaining variants show a pronounced drop in frequency, with many approaching zero or disappearing entirely. For milder selection in turbidostat, lower kanamycin concentration should be employed. Fig. 4B illustrates the effect of increasingly stringent selection conditions on binders’ enrichment. Overall, the results show that stronger selection conditions produce more pronounced enrichment or depletion effects, and turbidostat selection represent the harshest tested condition.

The impact of the linker sequence can also be assessed. Fig. 4C shows that the short linker (GSK) is more enriched compared to longer ones. Specifically, under the most stringent selection condition (T32_Kan[209], Supplementary Table 10), GSK, GSEK, and GSEAK account respectively for 66, 21 and 13% of the top 100 most frequent variants. Moreover, the GSK linker corresponds to 9 out of 10 most frequent variants.

Across the full set of variants, the so-called ‘rational designs’ showed higher average enrichment than AI designs. However, this trend was reversed considering only the top 100 variants (Fig. 4D). Overall, under the present experimental conditions, AI design produced a small fraction of high-quality solutions amid most low-quality variants, while rational design generated a distribution centered on intermediate-quality solutions.

Fig. 4E shows the effect of different selection conditions (no selection – Kan[0], Kan[348], or T32_Kan[209]) on residue frequencies by position, as well as the frequencies of the top 10 variants relative to the entire population.

### Binder validation (ITC and NMR)

The top two variants from the most stringent condition in batch and turbidostat selections were considered for production and evaluation by isothermal microcalorimetry (ITC). T32_Kan[209]_top2 was not kept in the list, as it differed by only one amino acid from T32_Kan[209]_top1. The three retained variants, T32_Kan[209]_top1 (AEEEAARFAALLAALP-GSK-ip4mutG), Kan[348]_top1 (SAAARWAAALAALP-GSK-ip4mutG) and Kan[348]_top2 (AAAERARFAALLAKLP-GSK-ip4mutG), all holding the GSK linker, were expressed, purified, and their affinities for ASF1A_N_ measured using ITC (Supplementary Fig. 23).

The best binder, T32_Kan[209]_top1, showed an affinity of 152 ± 25 nM corresponding to a 70-fold improvement compared to ip4mutG (*K_d_* = 10.6 ± 1.0 μM), followed by Kan[348]_top1 (235 ± 26 nM, 50-fold improvement), and Kan[348]_top2 (581 ± 98 nM, 20-fold improvement). Remarkably, the measured affinities correlated with their selection stringency and enrichment levels.

We next analyzed the binding mode of the best binder, T32_Kan[209]_top1, with ASF1 using NMR spectroscopy. Chemical-shift perturbation of uniformly labeled ASF1A_N_ upon addition of unlabeled T32_Kan[209]_top1 was measured and then mapped on the AlphaFold3 model of the corresponding complex (Fig. 5). The surfaces interacting with both, the initial ip4_mutG peptide, and the designed N-terminal extension are highlighted, showing that the chemical-shift changes are in good agreement with the predicted binding interface.

**Figure 5:**
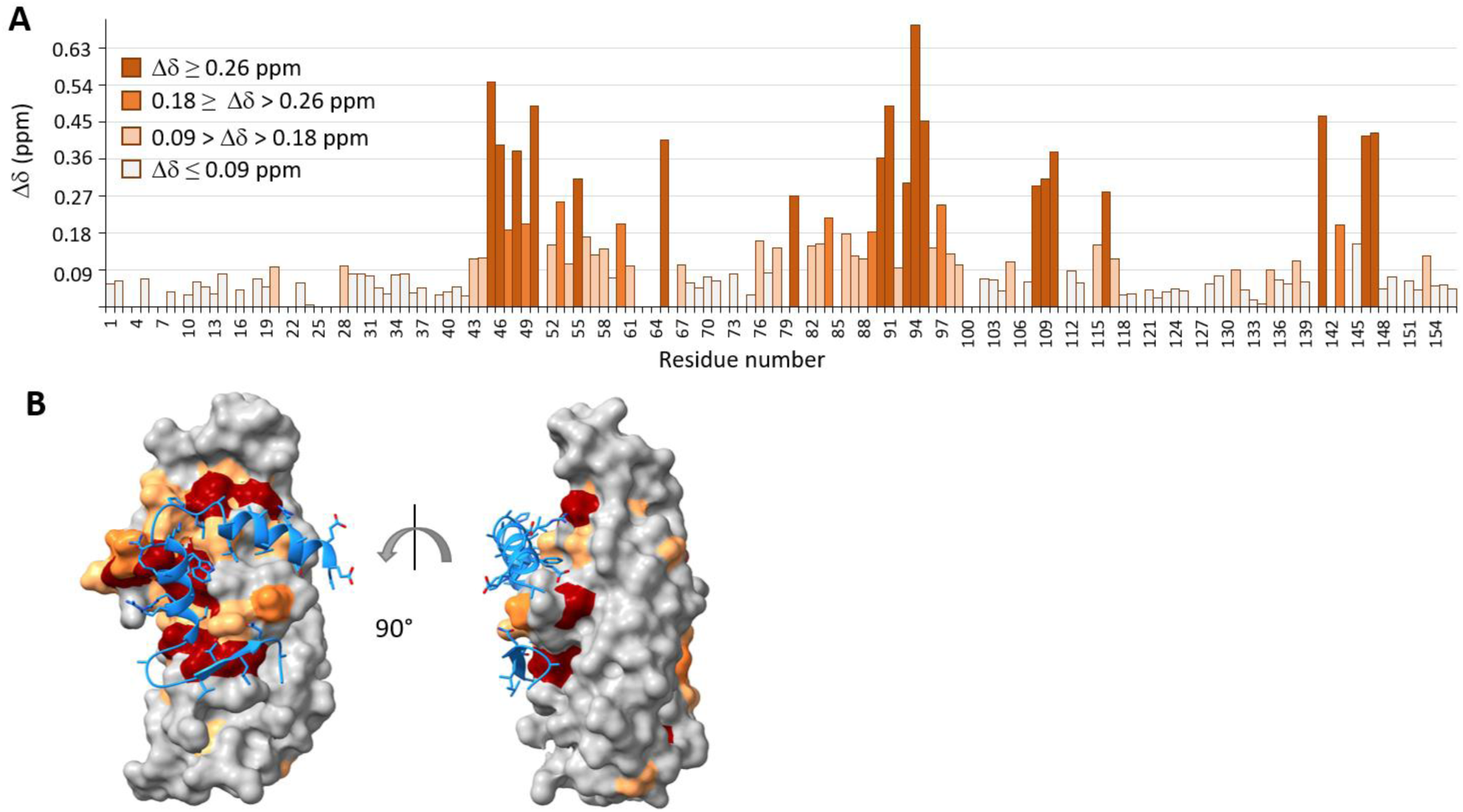
ASF1A_N_ — T32_Kan[209] binding mode. (**A**) NMR titration of ^15^N-ASF1A_N_: chemical shift perturbation (CSP) observed after the addition of an excess of T32_Kan[209]_top1 peptide (^15^N-ASF1A_N_: T32_Kan[209]_top1 ratio = 1:2). Chemical shift changes of residues are highlighted with the following color code: dark red (Δδ ≥ 0.26 ppm), dark orange (medium: 0.18 ≥ Δδ > 0.26 ppm), light orange (low: 0.09 > Δδ > 0.18 ppm), gray (background: Δδ ≤ 0.09 ppm). (**B**) AlphaFold 3 model of ASF1A_N_ bound to T32_Kan[209]_top1. The optimized peptide is shown as a cartoon and colored in blue. ASF1A_N_ surface is colored as in A.

## DISCUSSION

Building from previously described B2H systems based on λ *cI* and *E. coli rpoA* fusions engineered by Ann Hochschild (Harvard University), Rama Ranganathan (University of Chicago), and their colleagues, we engineered optimized variants with improved quantitative properties (qB2H). In early versions of Hochschild’s B2H, the PPI responsive promoter had an OR2 cI-binding element centered at position −62 from transcription start (*pLac* OR2-62) and it was inserted into the genome of KS1 strain (MC1000 F’ *lacI*^q^ with *pLac* OR2-62 *lacZ*)^9^ while later versions employed the OL2 element at the same position (*pLac* OL2-62) and was inserted into the F episome of FW102 strain^33^. Ranganathan and cols^14^ proposed an alternative implementation in three plasmids (pZA31-RNAα, pZS22 and pZE1RM-eGFP) that used MC4100 Z1 strain as chassis (MC4100 cells that express LacI^q^ and TetR repressors). pZA31-RNAα was designed to express *rpoA* fusions, pZS22 to express cIE34P fusions and pZE1RM-eGFP to provide an eGFP-based reporter under control of a λ *pRM* promoter carrying a C-to-T mutation at the 10th position of the OR3 element, which abolishes cI binding and provides stronger output^34^. Based on these system features, we anticipated incompatibilities with accurate quantification of protein–protein interactions (PPIs), which were confirmed by the results and are expected to negatively impact the generation of high-quality datasets. Machine learning (ML) models have been increasingly employed in diverse fields, including protein engineering, where data quality is well recognized as crucial for model training, interpretation, and application.

Although a similar system was recently reported by Kosuri and colleagues^35^, it relies on cAMP overproduction as a function of the PPI affinity. While this represents a significant advance toward a quantitative B2H, the CRP–cAMP complex directly regulates approximately 200 genes in *E. coli* by binding with high-affinity^36,37^, with over 100 of these genes involved in metabolic pathways for alternative carbon source^38^. Computational analyses further identify more than 10,000 lower-affinity sites, indicating broader effects mediated by weaker interactions^37^. Such non-targeted effects may impose unnecessary burdens and potentially result in stochastic consequences in specific cases. The growth medium, specially glucose concentration, modulates intracellular cAMP levels and thus influences B2H assay outcomes. The reporter used, sfGFP, may result in counterselection against high-affinity clones, as observed here. Unfortunately, we could not find clear data correlating *K_d_* values with signal intensity. Although the authors claimed plasmids were deposited at Addgene, we could not locate them, limiting adoption and reproducibility in the research community.

In contrast, the specific interaction between λ cI and its cognate DNA sequence is expected to enhance insulation of the synthetic system, thereby minimizing off-target effects and improving robustness across laboratories. In the case of the qB2H system described here, all necessary materials have been made publicly available through Addgene.

The qB2H systems presented here demonstrate high-quality quantitative metrics, indicating reliable assay performance. The latest dual reporter version, v3, provides a high signal-to-noise ratio (27.49 for the fluorescent reporter), *K_d_*-Signal consistency (*R*^2^ = 0.952) and low inter experiment variation on (CV = 7.04% for fluorescent reporter; CV = 4.97-6.12% for kanamycin resistance reporter) in SB39 strain. pB2H v4, was derived from v3 by replacing the dual reporter with a single antibiotic resistance marker and shows slightly improved metrics such as high *K_d_*-Signal consistency (coefficient of determination, *R*^2^ *=* 0.99), reproducibility (Pearson correlation of pseudo replicates, *r* = 0.99; *p* = 0, *n* = 734) and low inter-experiment variation (CV = 2.19–6.46%). Quantitative fitness values can be derived from NGS data. Barcode implementation provides statistical robustness and enables identification of outlier clones.

When sequencing resources are limited or rapid screening is needed, colonies can be analyzed on LB–agar plates with inducers and antibiotics. In qB2H v3, interaction strength can be estimated from colony fluorescence (or cytometry – stronger interactions yield higher brightness; see Supplementary Fig. 25) or colony size in the presence of kanamycin. In qB2H v4, only colony size can be evaluated. Antibiotic pressure can promote off-target adaptations (e.g., efflux pumps, altered metabolism, ribosomal mutations). To exclude off-target resistance events, selected target variants can be subcloned into the qB2H v3 vector and transformed into fresh cells for rapid clonal validation. Regarding general system features, the results indicate that higher selection pressures enhance the predictive relationship between enrichment and *K_d_* and reduce stochasticity. Nevertheless, very stringent selection pressures, such those that can be achieved in turbidostat experiments, can drastically impact population outcome, leading towards the large dominance of high-affinity binders and extinction of low-affinity ones. That strategy can be useful for the isolation of high-affinity binders, but it would be injudicious when trying to generate comprehensive datasets.

Building on the current qB2H system, we anticipate it will serve as a versatile platform for a wide range of projects. Supplementary Table 12 and Fig. 26 demonstrate the general applicability of qB2H versions by evaluating additional model interactions, protein folding and quaternary structure, cell chassis, and induction conditions. Encoding both interaction partners on a single plasmid (pB2H v2–v4) makes the system well suited for dual-partner deep mutational scanning (dpDMS) analyses —a method with few reliable, user-friendly solutions—thus broadening access to this approach within scientific and engineering communities (see “Materials, data and code availability”). Applications include, but are not limited to, PPI discovery and validation, interface mapping, identification of binding hot spots, interface engineering of orthologous complexes using dpDMS, high-throughput binder screening, data generation for AI-driven studies, and 3D modeling of protein complexes constrained by experimental data. For complex modeling, AI-based structure predictions often provide high-quality subunit models, while interface mapping via single-partner DMS (spDMS) helps verify uncertain complex models. Combined with dpDMS, this system offers reliable constraints to guide modeling of challenging complexes, such as antigen–antibody interactions^39^, integrating experimentally informed generative AI approaches^40^.

## METHODS

### Strains, construction of plasmids, and precultures

XL1-Blue chemically competent cells were used for routine cloning. Marionette Clo, XL1-Blue and SB39 strains were used as chassis for the experiments (detailed strain related information available in Supplementary Table 11; SB39 strain is available at Addgene, ID: 235121).

Detailed information about the oligonucleotides and the construction of vectors used in this work is available as supporting information (Plasmids construction, Supplementary Table 6: oligonucleotides, Supplementary Table 2: plasmids and, Supplementary Fig. 1).

Briefly, plasmids related to the two-plasmids system were derived from pBRα and pACλcI–β-flap, engineered by Ann Hochschild and colleagues (Harvard University)^41^. The latter, pACλcI–β-flap, was modified to include à B2H bicistronic reporter comprising eGFP and a kanamycin resistance coding region and to create a cI_E34P (GenBank: AAA96581.1; residues 1-237) fusion with the N-terminal domain of human ASF1A, ASF1A_N_ (UniProt: Q9Y294; residues 1-156) or the N-terminal domain of human ASF1B, ASF1B_N_ (UniProt: Q9NVP2 residues 1-156) giving rise to pAC-cI_E34P plasmid series. pBRα plasmid series correspond to the expression of 5 peptides (corresponding to different affinities against ASF1, Supplementary Table 1) fused to the alpha subunit of *Escherichia coli* RNA polymerase (RpoA lacking 81 C-terminal residues: residues 1-248; GenBank: AAC76320.1). The two-plasmid system from v1 series was composed of one plasmid of pBRα and one plasmid of pAC-cI_E34P series.

Single-plasmids from v2 series were created by assembling all the required B2H elements (from pAC-cI_E34P and pBRα series) into a single-plasmid. Single-plasmids from v3 series derived from v2 by invalidating a Dcm site in B2H-responsive promoter (*pOL2-62_L*) to reduce stochasticity, and by adjusting the cI fusion expression using a strong inducible promoter combined with a weak RBS to maximize signal output and increased dynamic range. Finally, plasmids from v4 series were derived from v3 series by converting the bicistronic reporter into a single-protein reporter (KanR), optimizing its RBS strength through a weak RBS library and sequencing small colonies in inducing LB-Agar supplemented with kanamycin (100 µg/mL).

Unless otherwise stated, precultures were inoculated late afternoon and grown overnight (37 °C, 190 rpm) from glycerol stocks in 10ml LB supplemented with antibiotics as required. The following final concentrations were used: ampicillin (100 µg/mL for single-plasmid selection and 75 µg/mL for two-plasmid set-ups) and chloramphenicol (34 µg/mL for single-plasmid and 25 µg/mL for two-plasmids set-ups), aTc (200 ng/µL for single-plasmid set-ups from the v3 and v4 series) and IPTG (20 nM for two-plasmid from v1 and single-plasmid from v2 series; 200 µM for single-plasmid from v3 and v4 series).

### qB2H experiments using a fluorescent reporter

Precultures of untransformed (for intrinsic fluorescence evaluation) and transformed SB39 cells were diluted (3 µL into 300 µL of fresh media) in 2 ml microtubes and grown in ThermoMixer C (Eppendorf; 37 °C, 900 rpm) for 2h. Cultures with an OD_600_ of approximately 0.5 were then mixed with 100 µL of the same media containing 4X inducer concentration. Final inducer concentrations, chosen based on maximal signal output, were 20 µM IPTG for v1 or v2 and 200 µM IPTG plus 200 µM aTc for v3 series. The induced cultures were grown overnight at 18 °C, diluted (1 µL) in freshly filtered (0.22 µm) PBS buffer (1 mL per OD unit), and analyzed using GUAVA EasyCyte HT (488 nm blue laser). The gating for event analysis was set to minimize background noise (fewer than 100 gated events per 10 µL of sterile PBS). Ten thousand events were collected per culture, and the Mean Fluorescence Intensity (MFI) recorded. To allow comparison across independent experiments, SB39 cells co-transformed with VN519 (pBRα-rpoA-ip3_mut3A) and VN550 (pAC-cI_E34P-ASF1A_Term_f1-pOL2-62_L-eGFP-KanR), or VN517 (pBRα-rpoA-ip3) and VN550 were included as internal controls. Their MFI were used to normalize those of other conditions (*x*) using the following equation:

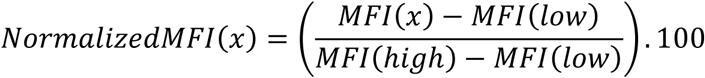

MFI(x) represents the mean fluorescence intensity of a given condition or culture. MFI(low) corresponds to the fluorescence produced by SB39 cells co-transformed with VN519 and VN550, indicating no detectable interaction (K_d_ > 100,000 nM). In contrast, MFI(high) corresponds to the fluorescence generated by SB39 cells co-transformed with VN517 and VN550, indicating a strong interaction (K_d_ = 55 nM). Each experiment was conducted in three independent assays performed on different days. Figures were generated using Python notebooks in conjunction with Matplotlib."

### MIC_50_ estimations

Precultures of transformed SB39 cells were diluted in 50 mL tubes (50 µL in 5 mL of fresh media supplemented with 17.5 µg/mL chloramphenicol) and grown in an orbital shaker (37 °C, 190 rpm). After 2 h, cultures (OD_600_ ∼ 0.5) were adjusted to OD_600_ = 0.05 in the same media containing inducers: 20 µM IPTG (v2 series) or 200 µM IPTG plus 200 ng/mL aTc (v3 or v4 series). Cultures were incubated at 37 °C for an additional 2 h. Subsequently, 300 µL of culture at OD600 = 0.1 or 0.02 was prepared in induced media and mixed with 300 µL of the same media supplemented with varying 2X concentrations of kanamycin. Cultures were incubated in a ThermoMixer C (Eppendorf; 37 °C, 950 rpm), and OD_600_ was measured after 4 and 6 h. The MIC_50_ for each condition (including strain, plasmid, starting OD_600_, and time in the presence of kanamycin) was calculated by fitting the values of kanamycin concentration and the normalized OD (OD_600_ normalized to the condition without antibiotics, assumed as 100% growth).

Python code for these calculations is available as a notebook. Each experiment was conducted in triplicate, with each replicate performed on a different day.

### Comparison of XL1-Blue v2 and SB39 v4

A detailed description of the construction and screening of interface mapping libraries is provided in the Supporting Information (“Construction and testing of libraries for stochasticity assessment,” Supplementary Tables 2 and 6). Briefly, DNA fragments encoding each ASF1 binder peptide, each coupled to 10 distinct barcodes, were generated and cloned into v2 or v4 series plasmids. Sublibraries, each corresponding to a specific peptide, were used to transform XL1-Blue or SB39 cells, respectively. Equiproportional cell libraries comprising all barcoded peptides were then prepared. Enrichment and stochasticity for XL1-Blue v2 and SB39 v4, as well as for each peptide, were evaluated by next-generation sequencing (NGS) under selection pressures corresponding to 40%, 70%, 80%, or 100% MIC_50_ of the ASF1–ip3 interaction within each system (strain plus plasmid series).

The coefficient of variation (CV), expressed as the percentage of the standard deviation relative to the mean enrichment, was used as a proxy to assess stochasticity.

### Interface mapping

A detailed description of the construction and screening of interface mapping libraries is provided in the Supporting Information (“Construction and screening of interface mapping libraries,” Supplementary Tables 2 and 6; Supplementary Fig. 8). Briefly, independent position libraries of ASF1B_N_ were obtained from Twist Bioscience, and equimolar pools of positions were cloned into the v4 series plasmid containing rpoA-ip3 or rpoA-HIRA fusions. HIRA corresponds to residues 442–472 of Histone Regulatory Homolog A (Uniprot: P54198), harboring an arbitrary point mutation, C465S, to prevent intermolecular disulfide bond formation. Both ASF1B_N_ binders were chosen due to the availability of crystallographic structures in complex with ASF1 (PDB IDs 6F0G and 2I32, respectively). SB39 cells were transformed with ASF1 region-specific libraries corresponding to three distinct position pools (N-terminus, middle, or C-terminus) and overnight precultures were induced. Cell populations were analyzed before (T_0_) and after antibiotic selection (T_F_) to assess the impact of each position on binder interaction by next-generation sequencing (NGS) coupled with data analysis.

The results were processed using the mave2imap pipeline and Python notebooks to generate fitness and perturbation score datasets and figures (see Data Availability).

### Design and selection of binders

An AI pipeline integrating RFdiffusion, ProteinMPNN, and AlphaFold2 (detailed in the Supplementary Methods) was implemented to design N-terminal extensions of ip4_mutG, a weak ASF1 binder (K_d_ ∼10 µM). Briefly, N-terminal extensions of 11 to 16 amino acids were fused to linkers (GSEAK, GSEK, or GSK) and appended to the ip4_mutG coding sequence. Rational designs and controls (flexible N-terminal extensions) were incorporated into the library, ordered as oligonucleotide pools (IDT, oPoolsTM), and cloned into the v4 series plasmid for transformation into SB39 cells. Library selection was performed in two experimental setups: i) short batch selection and ii) prolonged continuous selection in a turbidostat (Chi.Bio^42^).

In the batch selection, induced cultures diluted to OD_600_ = 0.05 were exposed to kanamycin concentrations ranging from 0 to 348 µg/mL. After 4.5–5 hours, only cultures exhibiting a decrease in OD_600_ of ≥0.5 relative to antibiotic-free controls were subjected to NGS analysis. In the turbidostat setup, induced cultures at OD_600_ = 0.05 were exposed to 209 µg/mL kanamycin (120% MIC_50_ of ASF1B_N_–ip3 in the same vector). Cultures were maintained at OD_600_ 0.4 for 21 or 29 hours, followed by an additional 3-hour growth period after sample collection, totaling 24 and 32 hours.

Plasmids extracted from samples taken before (T0) and after selection (TF) were sequenced by NGS (Illumina), and reads corresponding to expected sequences were quantified to assess relative frequencies and enrichment.

### Expression and purification of ASF1’s binders

DNA sequence encoding T32_Kan[209]_top1, Kan[348]_top1 and Kan[348]_top2 peptides were cloned into pSMT3 plasmid as a (His)_6_-SUMO-peptide fusion proteins and expressed in *Escherichia coli* BL21 (DE3) Star. Protein expression was induced by 0.5 mM isopropyl-β-D-thiogalactopyranoside (IPTG) at 37 °C for 3 hours. Cells were collected by centrifugation and were then resuspended in the lysis buffer (50 mM Tris-HCl pH 8, 1% Triton, 500 mM NaCl, 5% glycerol) supplemented with proteinase inhibitors (100 µg/mL Pefabloc, 1 µg/mL Pepstatine, 0.5 µg/mL Leupeptine, 1 µg/mL Chymostatine, 1 mM PMSF, 8 µg/mL Aprotinine), 250 µM DTT and 1X Roche EDTA-free Complete protease inhibitor cocktail. Bacterial lysis was performed using a French press on frozen pellets. Lysates were immediately treated with Benzonase (12 U/mL) and 1 mM MgCl_2_, incubated at 4 °C for 15 minutes, then centrifuged at 20,000rpm for 20 minutes. Affinity purification was performed using a nickel-charged nitrilotriacetic acid (NiNTA) agarose column (Macherey Nagel). After elution (50 mM Tris pH 8, 150 mM NaCl, 300 mM imidazole), purified SUMO protease was added at a 1:40 mass ratio (protease:peptide) and incubated at 30 °C for 1 hour to cleave the fusion. Peptides were further purified on a Ressource RPC 3 mL column (Cytiva) and eluted via acetonitrile gradient, then lyophilized overnight using a SpeedVac system and resuspended in water. Final purification was achieved by size-exclusion chromatography using a Superdex 30 10/300 GL column (GE Healthcare) in 50 mM Tris-HCl pH 7.5. Peptide mass was confirmed by LC-MS. Fractions containing intact peptide were pooled and stored at −20 °C.

### Expression and purification of ASF1

Recombinant human ASF1A_N_ (residues 1-156) was produced in *Escherichia coli* as a (His)_6_– glutathione S-transferase (GST)–TEV cleavage site–ASF1 fusion protein using the pETM30 plasmid. Soluble (His)_6_-tagged GST fusion protein was purified on reduced glutathione agarose beads (Sigma-Aldrich). After overnight cleavage at room temperature with recombinant (His)_6_–Tobacco Etch Virus protease (TEV), the (His)6–GST tag and protease were captured on a nickel-charged nitrilotriacetic acid (NiNTA) agarose column (Macherey Nagel). The flow-through fraction containing hASF1A_N_ protein was further purified by anion exchange chromatography using a Resource Q 6 mL column (GE Healthcare). The protein was concentrated in an Amicon device (Millipore) and the buffer was exchanged to 50 mM Tris-HCl pH 7.5. Unlabeled hASF1A_N_ for ITC experiments was purified from cells grown in LB medium, whereas uniformly 15N- and/or 15N-13C-labeled ^15^N-^13^C-labeled ASF1A_N_ was purified from bacteria grown in M9 minimal medium supplemented with (^15^NH_4_)Cl (0.5 g/L; Eurisotop) as the sole nitrogen source and ^13^C-glucose (2g/L) as carbon source.

### Isothermal microcalorimetry (ITC)

Isothermal titration calorimetry (ITC) experiments were performed on a VP-ITC titration calorimeter (Microcal/Malvern) at 20 °C using 50 mM Tris-HCl buffer (pH 7.5). Concentrations of ASF1 and binders ranged from 5 to 90 μM. Optimal ASF1:binder ratios were chosen to achieve clear transition curves and minimize fitting errors. Both proteins were prepared in the same buffer and degassed for 5 minutes by sonication or vacuum. After equilibrating the sample cell at 293.15 K, 10 µL aliquots of binder solution (total of 30 injections) were injected into the ASF1 solution at 180-second intervals with the syringe rotating at 310 rpm until saturation was reached. Raw data were processed using Origin 7.0 software (OriginLab, Malvern) applying the One-Set of Sites binding model. Experiments were performed in duplicate, and representative curves for each binder are shown in Supplementary Figure 23.

### Chemical shift mapping of ASF1A_N_ upon peptide binding

The binding mode of binders was assessed using nuclear magnetic resonance (NMR) spectroscopy by monitoring amide chemical shifts of ASF1A_N_ residues. NMR experiments were performed at 293 K on Bruker NEO 700 MHz spectrometer equipped with a cryoprobe (Bruker). Purified uniformly labeled ^15^N hASF1A_N_(1-156) was diluted in the NMR buffer comprising 50 mM Tris-HCl (pH 7.5), 0.1 mM EDTA, 0.1 mM dextran sulfate sodium (DSS), 0.1 mM NaN3, protease inhibitor cocktail (per manufacturer’s instructions, Roche), and 10% D_2_O. Proton chemical shifts (in parts per million) were referenced relative to internal DSS, and ^15^N reference was set indirectly relative to DSS using frequency ratios^16^. NMR data were processed using Topspin (Bruker) and analyzed using Sparky (T. D. Goddard and D. G. Kneller, University of California San Francisco). Amide assignments were adopted from previous studies^43^. An excess of T32_Kan[209]_top1 binder was added at a ratio 1:2 and a two-dimensional ^1^H-^15^N SOFAST HMQC (heteronuclear multiple-quantum coherence) spectrum was recorded In the formula, and the NMR spectra are represented in Supplementary Fig. 24. Chemical shift perturbations were quantified for all resonances using ^13^C-^15^N labeled hASF1A_N_ in presence of T32_Kan[209]_top1. Chemical shift variation was calculated with the following formula:

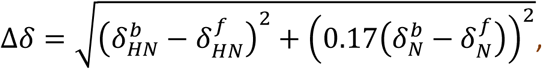

In the formula, δ represents the measured chemical shift value; b and f denote the bound or free forms, respectively, while HN or N to the amide proton or nitrogen atoms, respectively. The factor 0.17 is a scaling constant used to normalize proton and nitrogen chemical shift changes (expressed in parts per million). The calculated chemical shift variations were plotted as a function of residue number for all 156 residues of hASF1A_N_ (1-156) (Fig. 5).

### Statistical analyses

#### Engineering experiments

Flow cytometry and MIC50 results were obtained from three independent biological replicates (*n* = 3). The coefficient of determination (*R*^2^) was used to evaluate the correlation between *K_d_* and mean fluorescence intensity (MFI) or MIC50. Comparisons of MFIs or MIC_50_ values at different *K_d_* were performed using a one-tailed Student *t*-test assuming unequal variances (based on F-test results), with significance defined as *p*<0.05. The coefficient of variation (CV) was employed as a proxy for stochasticity, defined as the ratio of the standard deviation to the mean of measurements, expressed as a percentage.

#### Application experiments

In the context of binders’ optimization experiment, Pearson correlation coefficients (*PCC*, *r*) were used to assess correlations of variant frequencies or enrichment under different selection pressures. Only variants present in all conditions were used (*n* = 734 of 1134 variants) were included in these analyses.

### Data Availability

The atomic coordinates of the ASF1 complexes with ip3 and HIRA are available in the Protein Data Bank (PDB) under accession codes <u>6f0g</u> and <u>2i32</u>, respectively. Materials, source data, and software used in this study are provided. Key plasmids utilized herein have been deposited at Addgene under IDs 235097 to 235120. The engineered strain SB39 is also available through Addgene (ID: <u>235121</u>). Next-generation sequencing (NGS) data, additional results, Python code, and notebooks for data analysis and figure generation are accessible at Zenodo: https://zenodo.org/records/15690360. Interface mapping results were analyzed using the mave2imap pipeline, which is available on GitHub (https://github.com/synth-bio-evo/mave2imap) and Pypi (https://pypi.org/project/mave2imap/). Additional information on designs and data analyses is provided in the Supporting Information.

## AUTHOR INFORMATION

### Corresponding Author

Oscar H.P. Ramos − CEA, Département Médicaments et Technologies pour la Santé (DMTS), SIMoS, Université Paris-Saclay, 91191 Gif-sur-Yvette, France;

### Authors

Antoine Guyot − Sanofi Large Molecule Research, Vitry-sur-Seine, France; CEA, Département Médicaments et Technologies pour la Santé (DMTS), SIMoS, Université Paris-Saclay, 91191 Gif-sur-Yvette, France;

Emma Maillard − CEA, CNRS, Institute for Integrative Biology of the Cell (I2BC), Université Paris-Saclay, 91191 Gif-sur-Yvette, France;

Kelly Ferreira-Pinto − CEA, Département Médicaments et Technologies pour la Santé (DMTS), SIMoS, Université Paris-Saclay, 91191 Gif-sur-Yvette, France;

Laure Plançon-Arnould − CNRS, Institute for Integrative Biology of the Cell (I2BC), Université Paris-Saclay, 91191 Gif-sur-Yvette, France;

Aravindan A. Nadaradjane − CNRS, Institute for Integrative Biology of the Cell (I2BC), Université Paris-Saclay, 91191 Gif-sur-Yvette, France;

Françoise Ochseinbein − CEA, CNRS, Institute for Integrative Biology of the Cell (I2BC), Université Paris-Saclay, 91191 Gif-sur-Yvette, France;

Raphaël Guérois − CEA, CNRS, Institute for Integrative Biology of the Cell (I2BC), Université Paris-Saclay, 91191 Gif-sur-Yvette, France;

Loic Martin − CEA, Département Médicaments et Technologies pour la Santé (DMTS), SIMoS, Université Paris-Saclay, 91191 Gif-sur-Yvette, France;

### Authors contribution

- A.G. participated in molecular biology steps, performed experiments (fluorescent qB2H, MIC_50_, stochasticity minimization and selection of binders), protein production, wrote codes for data analysis and participated in the writing of this manuscript.
- E.M. participated in protein production, ITC, and NMR experiments. She also participated in the writing of this manuscript.
- K.F-P. participated in molecular biology experiments (fluorescent qB2H, MIC_50_, Interface mapping and NGS samples preparation).
- L.P-A. contributed to the development of protein and peptide production protocols.
- A.A.N. participated in mave2imap development.
- R.G. provided financial support; wrote codes for data analysis; suggested the interface mapping approach and participated in the writing of this manuscript.
- F.O. is the leader concerning the development of ASF1 binders applied to cancer, with abundant backtrack publication in the subject. She tested, evaluated complex formation by ITC and NMR, and participated in the writing of this manuscript.
- L.M. is the director of A.G. thesis. He provided financial support; participated in the design of qB2H system variants; participated in molecular biology steps, protein production, experiments (epitope mapping) and NGS samples preparation; participated in the writing of this manuscript.
- O.H.P.R. directed the development of the qB2H system. He provided financial support; designed qB2H system variants; proposed and participated in the implementation of the generative AI pipeline for peptide binders’ design; wrote codes for data analysis; coordinated manuscript writing and wrote most of it. He also prepared and submitted the code and data for distribution (GitHub, Pypi and Zenodo).

## Notes

- This work was financially supported by Sanofi (Collaboration agreement Sanofi/CEA). A.G. was supported by a CIFRE fellowship (N2022/1461) funded in part by the National Association for Research and Technology (ANRT) on behalf of the French Ministry of Education and Research, and in part by Sanofi.
- A.G. is a Sanofi employee and may hold shares and/or stock options in the company.
- The remaining authors declare no competing interest.

## SUPPORTING INFORMATION

- **Supporting_information.**pdf: main supporting information containing Supplementary Materials, Methods, file descriptions, and Figures.
- **Supplementary Table 1**: ASF1 — peptides interaction: interaction strength (calculated by ITC) between ASF1A_N_ or ASF1B_N_ against different peptides.
- **Supplementary Table 2**: Plasmids: contains data such as name, features, replication origin, corresponding antibiotics and inducers used concentrations and library related information.
- **Supplementary Table 3**: qB2H engineering cytometry data: raw and normalized MFI (mean fluorescence intensity) data obtained for different qB2H plasmid systems. Results of statistical test are also included.
- **Supplementary Table 4**: qB2H engineering MIC_50_ data: raw and normalized MIC_50_ (minimum inhibitory concentration) data obtained for different qB2H plasmid systems. It also includes results of statistical tests.
- **Supplementary Table 5**: Stochasticity data: results related to the comparison of some variables (selection pressure variation; strain or outliers’ removal) on key features of qB2H system (correlation between affinity and enrichment; homogeneity of data values).
- **Supplementary Table 6**: Oligonucleotides: contains data such as name, sequence, template, destination vector of PCR products, description and special features concerning oligonucleotides used for this publication.
- **Supplementary Table 7**: Twist library for interface mapping screening: information about the positional library of point mutants order from Twist Bioscience.
- **Supplementary Table 8**: Fitness data from interface mapping experiment, expressed as normalized enrichment values. The values were generated by using mave2imap pipeline.
- **Supplementary Table 9**: Perturbation data from interface mapping experiment, expressed as perturbation scores. The values were generated from fitness data analysis.
- **Supplementary Table 10**: Binders’ selection data: variant description and information such as their count, frequencies and, when relevant, their enrichments. The results of statistical analysis are also included.
- **Supplementary Table 11**: Strain description **—** strain related genotypes and features.
- **Supplementary Table 12**: Signal output of additional model complexes and the influence of induction conditions on IgG CH3 domain signal output.

## Supporting information

Promoters and plasmids

qB2H engineering flow cytometry data

qB2H engineering minimal inhibitory concentration (MIC50) data

Stochasticity data

Oligonucleotides

Twist library for interface mapping screening

Fitness data from interface mapping experiment

Perturbation data from interface mapping experiment

Strain description

Results from additional tests

Phylogenetic tree and frequency heatmap of variants

Phylogenetic tree and enrichment heatmap of variants

## ACKNOWLEDGMENTS

We thank the following scientists and collaborators for their contributions to this work:

- Professor Ann Hochschild (Harvard Medical School) for providing their plasmids and strain (FW102, beta galatosidase complementation-based reporter) that were used in preliminary tests. The plasmids pBRα pACλcI–β-flap were used as basis to construct several plasmids described in this work.
- Professor Rama Ranganathan (University of Chicago) for providing their plasmids and strain (MC4100-Z1, expression of TetR and LacI repressors for better control of inducible promoters) that were used in preliminary tests.
- Emmanuelle Vigne and Klervi Desrumeaux (SANOFI) for their regular scientific discussions.
- Robert Thai and Matthieu Fonvielle for their contribution to the measurement of molecular mass of purified peptides and proteins by LCMS.
- This work was supported by the French National Research Agency (ANR) under grant number ANR-15-CE11-0008-01.

## ABBREVIATIONS

2H: two-hybrid system
AA: amino acid
AI: artificial intelligence
aTc: anhydrotetracycline
B2H: bacterial two-hybrid
cfu: colony forming units
ci: confidence interval
cI: a lambda (λ) bacteriophage DNA binding protein
CV: coefficient of variation, ratio between the standard deviation and the mean (here represented as percentage)
Dcm site: DNA sequence (CCA/TGG) recognized by Dcm methylase which methylates the second C base of the motif in C5 position
dpDMS: dual-partner deep mutational scanning
FACS: fluorescence activated cell sorting
Gb: gigabases (1×10^9^ bases)
GG: Golden Gate, a scarless and one pot assembly method that uses type IIS restriction enzymes and a DNA ligase for DNA assembly
HIRA: Histone Regulatory Homolog A (a mutant C465S was used in this work)
ITC: isothermal titration calorimetry
*K_d_*: equilibrium dissociation constant, a ratio of koff/kon (inversely correlated to the affinity)
MFI: mean fluorescence intensity (AU)
MIC_50_: minimum inhibitory concentration (here antibiotics concentration that inhibits 50% bacterial growth compared to no antibiotics)
ML: machine learning
NGS: next generation sequencing
NT: nucleotide
ORF: open reading frame
PCA: protein complementation assay
PPI: protein-protein interaction
qB2H: quantitative bacterial two-hybrid
*rpoA*: DNA-directed RNA polymerase subunit alpha
T.I.R.: translation initiation rate
UMI: unique molecule identifier
WPS: whole plasmid sequencing.

## REFERENCES

1. Luo, Y. et al. ECNet is an evolutionary context-integrated deep learning framework for protein engineering. Nat Commun 12, 5743 (2021).

2. Li, M. et al. SESNet: sequence-structure feature-integrated deep learning method for data-efficient protein engineering. Journal of Cheminformatics 15, 12 (2023).

3. Notin, P., Rollins, N., Gal, Y., Sander, C. & Marks, D. Machine learning for functional protein design. Nat Biotechnol 42, 216–228 (2024).

4. Younger, D., Berger, S., Baker, D. & Klavins, E. High-throughput characterization of protein– protein interactions by reprogramming yeast mating. Proceedings of the National Academy of Sciences 114, 12166–12171 (2017).

5. Stynen, B., Tournu, H., Tavernier, J. & Van Dijck, P. Diversity in Genetic *In Vivo* Methods for Protein-Protein Interaction Studies: from the Yeast Two-Hybrid System to the Mammalian Split-Luciferase System. Microbiol Mol Biol Rev 76, 331–382 (2012).

6. Michnick, S. W., Ear, P. H., Manderson, E. N., Remy, I. & Stefan, E. Universal strategies in research and drug discovery based on protein-fragment complementation assays. Nat Rev Drug Discov 6, 569–582 (2007).

7. Richardson, R. M. & Pascal, S. M. Bacterial two-hybrid systems evolved: innovations for protein-protein interaction research. J Bacteriol 207, e00129–25.

8. Fields, S. & Song, O. A novel genetic system to detect protein–protein interactions. Nature 340, 245–246 (1989).

9. Dove, S. L., Joung, J. K. & Hochschild, A. Activation of prokaryotic transcription through arbitrary protein-protein contacts. Nature 386, 627–630 (1997).

10. Luo, Y., Batalao, A., Zhou, H. & Zhu, L. Mammalian Two-Hybrid System: A Complementary Approach to the Yeast Two-Hybrid System. BioTechniques 22, 350–352 (1997).

11. Adams, B. L. The Next Generation of Synthetic Biology Chassis: Moving Synthetic Biology from the Laboratory to the Field. ACS Synth. Biol. 5, 1328–1330 (2016).

12. Chang, M., Ahn, S. J., Han, T. & Yang, D. Gene expression modulation tools for bacterial synthetic biology. Biotechnology for Sustainable Materials 1, 6 (2024).

13. Bittihn, P., Din, M. O., Tsimring, L. S. & Hasty, J. Rational engineering of synthetic microbial systems: from single cells to consortia. Curr Opin Microbiol 45, 92–99 (2018).

14. McLaughlin Jr, R. N., Poelwijk, F. J., Raman, A., Gosal, W. S. & Ranganathan, R. The spatial architecture of protein function and adaptation. Nature 491, 138–142 (2012).

15. Bakail, M. et al. Design on a Rational Basis of High-Affinity Peptides Inhibiting the Histone Chaperone ASF1. Cell Chem Biol 26, 1573–1585.e10 (2019).

16. Mbianda, J. et al. Optimal anchoring of a foldamer inhibitor of ASF1 histone chaperone through backbone plasticity. Sci Adv 7, eabd9153 (2021).

17. Perrin, M. E. et al. Unexpected binding modes of inhibitors to the histone chaperone ASF1 revealed by a foldamer scanning approach. Chem Commun (Camb) 59, 8696–8699 (2023).

18. Lei, X., Tian, W., Zhu, H., Chen, T. & Ao, P. Biological Sources of Intrinsic and Extrinsic Noise in cI Expression of Lysogenic Phage Lambda. Sci Rep 5, 13597 (2015).

19. Sedhom, J. & Solomon, L. A. Lambda CI Binding to Related Phage Operator Sequences Validates Alignment Algorithm and Highlights the Importance of Overlooked Bonds. Genes 14, 2221 (2023).

20. Sample, V., Newman, R. H. & Zhang, J. The structure and function of fluorescent proteins. Chem. Soc. Rev. 38, 2852–2864 (2009).

21. Matsuyama, C. et al. Metabolome analysis of metabolic burden in Escherichia coli caused by overexpression of green fluorescent protein and delta-rhodopsin. J Biosci Bioeng 137, 187–194 (2024).

22. Meyer, A. J., Segall-Shapiro, T. H., Glassey, E., Zhang, J. & Voigt, C. A. Escherichia coli “Marionette” strains with 12 highly optimized small-molecule sensors. Nat Chem Biol 15, 196–204 (2019).

23. Bandiera, L., Furini, S. & Giordano, E. Phenotypic Variability in Synthetic Biology Applications: Dealing with Noise in Microbial Gene Expression. Front. Microbiol. 7, (2016).

24. Udekwu, K. I., Parrish, N., Ankomah, P., Baquero, F. & Levin, B. R. Functional relationship between bacterial cell density and the efficacy of antibiotics. J Antimicrob Chemother 63, 745–757 (2009).

25. LaPlante, K. L. & Rybak, M. J. Impact of High-Inoculum Staphylococcus aureus on the Activities of Nafcillin, Vancomycin, Linezolid, and Daptomycin, Alone and in Combination with Gentamicin, in an In Vitro Pharmacodynamic Model. Antimicrobial Agents and Chemotherapy 48, 4665 (2004).

26. Tu, Q. et al. Room temperature electrocompetent bacterial cells improve DNA transformation and recombineering efficiency. Sci Rep 6, 24648 (2016).

27. Pacesa, M., et al. BindCraft: one-shot design of functional protein binders. bioRxiv 2024.09.30.615802 (2025) doi:10.1101/2024.09.30.615802.

28. Watson, J. L. et al. De novo design of protein structure and function with RFdiffusion. Nature 620, 1089–1100 (2023).

29. Dauparas, J. et al. Robust deep learning-based protein sequence design using ProteinMPNN. Science 378, 49–56 (2022).

30. Jumper, J. et al. Highly accurate protein structure prediction with AlphaFold. Nature 596, 583– 589 (2021).

31. Evans, R., et al. Protein complex prediction with AlphaFold-Multimer. bioRxiv 2021.10.04.463034 (2022) doi:10.1101/2021.10.04.463034.

32. Mirdita, M. et al. ColabFold: making protein folding accessible to all. Nat Methods 19, 679–682 (2022).

33. Dove, S. L. & Hochschild, A. Bacterial Two-Hybrid Analysis of Interactions between Region 4 of the ς ^70^ Subunit of RNA Polymerase and the Transcriptional Regulators Rsd from *Escherichia coli* and AlgQ from *Pseudomonas aeruginosa*. J Bacteriol 183, 6413–6421 (2001).

34. Meyer, B. J., Maurer, R. & Ptashne, M. Gene regulation at the right operator (*O*R) of bacteriophage λ. Journal of Molecular Biology 139, 163–194 (1980).

35. Boldridge, W. C. et al. A multiplexed bacterial two-hybrid for rapid characterization of protein– protein interactions and iterative protein design. Nat Commun 14, 4636 (2023).

36. Grainger, D. C., Hurd, D., Harrison, M., Holdstock, J. & Busby, S. J. W. Studies of the distribution of Escherichia coli cAMP-receptor protein and RNA polymerase along the E. coli chromosome. Proceedings of the National Academy of Sciences 102, 17693–17698 (2005).

37. Robison, K., McGuire, A. M. & Church, G. M. A comprehensive library of DNA-binding site matrices for 55 proteins applied to the complete Escherichia coli K-12 genome. J Mol Biol 284, 241–254 (1998).

38. Balsalobre, C., Johansson, J. & Uhlin, B. E. Cyclic AMP-Dependent Osmoregulation of crp Gene Expression in Escherichia coli. Journal of Bacteriology 188, 5935–5944 (2006).

39. Hitawala, F. N. & Gray, J. J. What does AlphaFold3 learn about antibody and nanobody docking, and what remains unsolved? mAbs 17, 2545601 (2025).

40. Team, C. D. et al. Chai-1: Decoding the molecular interactions of life. 2024.10.10.615955 Preprint at 10.1101/2024.10.10.615955 (2024).

41. Kuznedelov, K. et al. A Role for Interaction of the RNA Polymerase Flap Domain with the σ Subunit in Promoter Recognition. Science 295, 855–857 (2002).

42. Steel, H., Habgood, R., Kelly, C. L. & Papachristodoulou, A. In situ characterisation and manipulation of biological systems with Chi.Bio. PLoS Biol 18, e3000794 (2020).

43. Mousson, F. et al. 1H, 13C and 15N resonance assignments of the conserved core of hAsf1 A. J Biomol NMR 29, 413–414 (2004).

