## Supplementary figures and images for "Optimized quantitative bacterial two-hybrid (qB2H) for protein-protein interaction assessment"

### Phylogenetic tree and enrichment heatmap of variants

**Fitness of N-terminal extensions**

*– heatmap color gradient based on enrichment –*

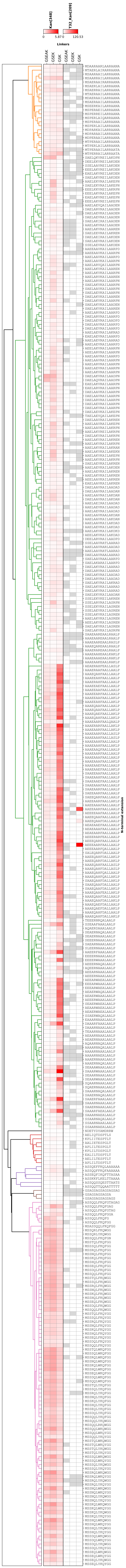

### Phylogenetic tree and frequency heatmap of variants

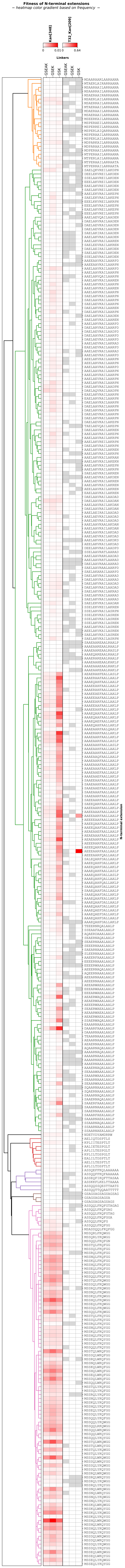
