## Supporting information for "Optimized quantitative bacterial two-hybrid (qB2H) for protein-protein interaction assessment"

### qB2H, an optimized and versatile bacterial two-hybrid for quantitative assessment of protein-protein interactions

**TABLE OF CONTENTS**

1. Methods ... 4

1.1. Plasmid construction ... 4

1.2. Construction and testing of libraries for stochasticity assessment ... 5

1.3. Construction and screening of interface mapping libraries ... 6

1.4. Design, library construction, and selection of binders ... 7

1.4.1. Binder design ... 7

1.4.2. Library construction ... 8

1.4.3. Short selection runs (batches) ... 8

1.4.4. Long selection runs (continuous cultures under stable cell density—turbidostat) ... 8

1.4.5. NGS sample preparation and analysis ... 9

1.4.6. Comparison of samples submitted to different selection pressures ... 9

1.5. Calculation of the required amount of gygabases and NGS data treatment ... 9

2. Tables (description of Excel files) ... 9

2.1. Table S1: ASF1–peptide interactions ... 9

Table S2: Promoters and plasmids ... 10

Table S3: qB2H engineering flow cytometry data ... 10

Table S4: qB2H engineering minimal inhibitory concentration ($MIC_{50}$) data ... 10

Table S5: Stochasticity data ... 10

Table S6: Oligonucleotides ... 10

Table S7: Twist library for interface mapping screening ... 10

Table S8: Fitness data from interface mapping experiment ... 10

Table S9: Perturbation data from interface mapping experiment ... 10

Table S10: Binders' selection data ... 10

Table S11: Strain description ... 10

Table S12: Results from additional tests ... 10

3. Figures ... 11

Figure S1: Flowchart of plasmid version construction ... 13

Figure S2: Effect of kanamycin concentration on proliferation of SB39 pB2H v3 (starting OD = 0.05, time = 4 h) ... 14

Figure S3: Effect of kanamycin concentration on proliferation of SB39 pB2H v3 (starting OD = 0.05, time = 6 h) ... 15

Figure S4: Effect of kanamycin concentration on proliferation of SB39 pB2H v3 (starting OD = 0.01, time = 6 h) ... 16

Figure S5: Effect of kanamycin concentration on proliferation of SB39 pB2H v4 (starting OD = 0.05, time = 4 h) ... 17

Figure S6: Effect of kanamycin concentration on proliferation of SB39 pB2H v4 (starting OD = 0.05, time = 6 h) ... 18

Figure S7: Effect of kanamycin concentration on proliferation of SB39 pB2H v4 (starting OD = 0.01, time = 6 h) ... 19

### 1. Methods

#### 1.1. Plasmid construction

Detailed information regarding plasmids and oligonucleotides referenced in this section can be found in Supplementary Tables 2 and 6, respectively, along with Supplementary Figure 1. Q5® high-fidelity polymerase (NEB) was employed for all PCR amplification intended for cloning.

Two-plasmid system (v1 series) was based on pBRα and pACλcI–β-flap[1], kindly provided by Professor Ann Hochschild, Harvard University. pBRα was utilized to create rpoA fusions by inserting DNA fragments encoding various alternative peptides (between *Not*I and *Bam*HI sites) that correspond to different affinities (Supplementary Table 1): ip3_mut3A, ip1, ip2, ip3, and ip4[2]. pACλcI–β-flap was modified to enable the expression of cI fusions and to facilitate two-hybrid interactions through the following steps: (i) introduction of λ cI E34P mutation to [illegible], achieved by plasmid mutagenesis using primers OR131 and OR132; (ii) replacement of β-flap (between *Not*I and *Bam*HI restriction sites) with DNA encoding human ASF1A$_{N}$ (GenBank: CAG33628.1), which was amplified from the plasmid"pETM30_(His)6-GST-Tev site-ASF1A$_{N}$"[illegible] using primers OR564 and OR565; (iii) insertion of synthetic DNA construct (ordered from GeneArt) between the cI fusion gene and p15A replication origin, containing the following elements: "bi-directional terminator (BBA_B1007)_*Pac*I-OL2-62_L-*Bsi*WI_RBS_eGFP-RBS-KanR_f1_bi-directional terminator (Bba_B0014)". Subsequently, various B2H-responsive promoters were inserted between *Pac*I and *Bsi*WI sites of the resulting vector (VN550) using hybridized oligonucleotides. These modifications led to the creation of the pAC-cI_E34P plasmid series, which features eGFP (GenBank: ANC98519.1) and aminoglycoside O-phosphotransferase (GenBank: HCE8982156, KanR) as reporters for the two-hybrid system. The plasmid variant VN627 was generated by substituting the human ASF1A$_{N}$ (located between *Not*I and *Bam*HI sites) in VN550 with human ASF1B[illegible] sequence (GenBank: NP_060624.1) amplified from "pETM30_(His)6-GST-Tev site-ASF1B$_{N}$" using primers OR865 and OR866.

Before single-plasmid construction, a VN627 derivative was constructed to: (i) replace *Not*I and *Bam*HI sites with *Acc*65I and *Sac*I sites, respectively, using primers OR1040 and OR1041; (ii) replace the strong ribosome binding site (RBS) between eGFP and KanR[illegible] and insert a *Spe*I site downstream of the KanR-encoding region. This was accomplished by Gibson Assembly, using Nebuilder® HiFi to combine two PCR fragments (generated by primers OR466+OR467 and OR1593+OR1624).

Single-plasmid VN991 was constructed by Gibson Assembly of three fragments: (1) a PCR amplicon from a VN627 derivative using primers OR337 and OR340 (comprising almost the entire plasmid); (2) a PCR amplicon of VN515 using primers OR339 and OR869 (comprising pLpp+lac_UV5-rpoA-ip1 gene) and; (3) hybridized oligonucleotides OR1024 and OR1025 (corresponding to the L3S2P21 terminator). Plasmid VN991 served as basis for constructing other plasmids from v2 series by replacing ip1 peptide coding sequence between *Not*I and *Bam*HI sites with hybridized oligonucleotides encoding alternative peptides.

A new generation of single-plasmid system with two reporters (v3 series; eGFP + KanR) was constructed in multiple steps from VN990. First, VN990 was opened upstream the λ cI ORF by PCR using oligonucleotides OR213 and OR1449 (F1). A fragment intended for *Bsa*I-based Golden Gate cloning was produced by PCR filling using oligonucleotides OR1903 and OR1904 (F2). Both fragments (F1+F2) were assembled using Nebuilder HiFi and the resulting plasmid, VN1170, was used for Golden Gate cloning of dsDNA created by PCR filling of oligonucleotides OR1910 and OR1911. This dsDNA included a weak RBS library designed using the "de novo DNA" website (TAGWCCARCTCGCHAGSTCATATA; comprising 24 variants with predicted translational initiation rate (T.I.R.) values between 1.57 and 3,732.37; ) In order to select RBS providing the most intense B2H signal (by balancing cI fusion amounts compared to rpoA fusion). The resulting plasmid, VN1171, was screened for maximum fluorescent on LB-Agar supplement with 200 ng/mL aTc, 200 µM IPTG. The retained RBS correspond to a predicted T.I.R. of 371. Finally, we invalidated a Dcm site upstream the -35 site of pOL2-62_L promoter by replacing it (between *Pac*I/*Bsi*WI sites) with hybridized oligonucleotides OR1917 and OR1918. The resulting plasmid, VN1315, served as the foundation for other plasmids in the v3 series by replacing the DNA coding for ip3 (between *Not*I and *Bam*HI) with fragments encoding alternative peptides.https://www.denovodna.com) to select RBS providing the most intense B2H signal by optimizing cI fusion amounts relative to rpoA fusion levels. The resulting plasmid, VN1171, was screened for maximum fluorescence on LB-agar supplement with 200 ng/mL aTc and 200 µM IPTG. The selected RBS corresponded to a predicted T.I.R. of 371. Finally, we inactivated a Dcm site upstream of the -35 site of pOL2-62_L promoter by replacing it (between *Pac*I and *Bsi*WI sites) with hybridized oligonucleotides OR1917 and OR1918. The resulting plasmid, VN1315, served as the foundation for other plasmids in the v3 series by replacing the DNA coding for ip3 (between *Not*I and *Bam*HI sites) with fragments encoding alternative peptides.

Another series of single-plasmids variants harboring a single reporter (v4 series; KanR) was also created in a final effort to reduce stochasticity and the required kanamycin concentration during selection campaigns. First, the RBS and dual reporter region of VN1315 were replaced with a library of weak RBS variants designed using the "de novo DNA" website (ATTTGASAGSGGSASGTACD; comprising 48 variants with predicted T.I.R. between 0.7 and 4,914.55) directly fused to the KanR reporter. Specifically, VN1315 was digested with restriction enzymes *Bsi*WI and *Spe*I and recombined by Gibson assembly (Nebuilder HiFi) with the amplicon generated from the same plasmid using primers OR1861 and OR1865. The resulting plasmid library (VN1265) was screened for small colonies on LB-agar supplemented with 100 µg/mL kanamycin, 200µM IPTG, and 200 ng/mL aTc. The selected RBS had a predicted T.I.R. of 51. Other plasmids of this series were constructed by replacing the region encoding the ip3 peptide (between *Not*I and *Bam*HI sites) with DNA fragments encoding alternative peptides.

All plasmids were verified by Sanger sequencing or whole-plasmid sequencing (WPS, based on nanopore technology, Eurofins genomics).

Main plasmids and their sequence maps were deposited at Addgene (IDs from 235097 to 235120).

#### 1.2. Construction and testing of libraries for stochasticity assessment

Small libraries were constructed to evaluate relevant B2H variables and features, such as strain dependency, correlation between complex affinity and enrichment under selection pressure, and stochasticity.

First, acceptor vectors were constructed by digesting v2 or v4 series plasmids with *Not*I and *Bam*HI and inserting annealed oligonucleotides (OR2383 and OR2384) using Gibson assembly (NEBuilder HiFi, E2621) resulting in VN1896 and VN1897, respectively. Each ASF1 binder peptide was coupled to ten barcodes by PCR filling, and the generated dsDNA was cloned into the acceptor vectors by *Bsa*I-based Golden Gate cloning. XL1-Blue (v2 series) or SB39 (v4 series) were independently transformed by plasmid libraries from each binder peptide and grown overnight in LB medium containing 34 µg/mL chloramphenicol. On the following day, precultures were diluted 1:100 in the same medium and pooled at log phase ($OD_{600}$ = 0.4–0.6) to ensure equiproportional representation of binders coupled to barcodes. The pooled culture was then grown to early stationary phase ($OD_{600}$ = 2.5–3.5), and glycerol stocks were prepared.

Overnight cultures were grown from glycerol stocks in LB medium containing 34 µg/mL chloramphenicol (~16 h, 37 °C, 200 rpm). The next day, cultures were diluted to $OD_{600}$ = 0.05 in 150 mL LB medium supplemented with chloramphenicol (17 µg/mL) and inducers (XL1-Blue v2: 20 µM IPTG; SB39 v4: 200 µM IPTG, 200 ng/mL aTc). The cultures were grown for 2 h ($OD_{600}$ ~ 0.5), and 100 mL was immediately pelleted (6 800 × g, 3 min) for plasmid miniprep ($T_0$ sample). The remaining culture was used to prepare a diluted culture ($OD_{600}$ = 0.05, 50m L) in the same inducing medium supplemented with kanamycin at concentration corresponding to 40, 70, 80, or 100% of the estimated $MIC_{50}$ for the interaction ASF1—ip3

(XL1-Blue v2: 11.76 µg/mL kanamycin; SB39 v4: 174 µg/mL kanamycin). The cultures were incubated (37 °C, 200 rpm) until $OD_{600}$ ~ 1.0 or for at maximum of 7 hours. At that time, the cells were pelleted, plasmids extracted and used as template for direct generation of NGS samples by PCR ($T_{F-40}$, $T_{F-70}$, $T_{F-80}$, $T_{F-100}$ samples) using primer OR2889 in conjunction with one of the reverse primers (OR2890-OR2894). $T_0$ and $T_F$ samples were sequenced using Illumina NGS (NovaSeq X).

The generated data were analyzed using a dedicated Python script, and graphics were generated using Matplotlib in a JupyterLab notebook. The following z-score thresholds were used for outlier removal: 1 (default), 0.5, and 0.25.

Plasmid and oligonucleotide information are available at Supplementary Tables 2 and 6, respectively.

#### 1.3. Construction and screening of interface mapping libraries

To assess the applicability of the engineered qB2H for interface mapping, we selected two binders with solved structures in complex with ASF1: ip3 (PDB: 6F0G) and HIRA (PDB: 2I32).

Plasmid libraries were constructed by Golden Gate cloning of pooled ASF1 surface positions – as linear dsDNA ordered from Twist Bioscience (Table S7) – into acceptor plasmids (v4 series; Table S2) VN1272 (rpoA-ip3 fusion) or VN1279 (rpoA-HIRA fusion). Briefly, single-position libraries were pooled based on three ASF1B regions: N-terminus (K3–V62), Middle (A35–L96), and C-Terminus (V90–D154). Equimolar pools were cloned into the acceptor vectors by *Bsa*I-based Golden Gate assembly, followed by XL1-Blue transformation. This yielded the stock library vectors VN1296 (ip3 vs. $ASF1B_N$ – C-terminus), VN1297 (HIRA vs. $ASF1B_N$ – Middle), VN1382 (ip3 vs $ASF1B_N$ – N-terminus), and VN1383 (HIRA vs. $ASF1B_N$ – N-terminus). Stock libraries were amplified, and their plasmids were extracted using the GeneJet Miniprep kit (Thermo Fisher Scientific). Each working library was obtained by performing 4–6 Marionette Cloning (Clo) transformations using 100 ng of stock library plasmid. Library construction statistics are available at Supplementary Table 2.

Precultures of the working library were grown overnight in LB medium supplemented with 34 µg/mL chloramphenicol (~16 h, 37 °C, 200 rpm). The next day, cultures were diluted to $OD_{600}$ = 0.05 in 150 mL LB medium containing chloramphenicol (17 µg/mL) and inducers (200 µM IPTG and 200 ng/mL aTc). After 2 h ($OD_{600}$ ~ 0.5), 100 mL was pelleted (6,800 x g, 3 min.) for plasmid extraction ($T_0$ sample). The remaining culture was diluted to $OD_{600nm}$ = 0.05 in 50 mL of the same inducing medium supplemented with 20 µg/mL kanamycin and incubated for 4–7 h (final $OD_{600}$ ~ 1.0). Cells were then pelleted and plasmids extracted ($T_F$ sample).

$T_0$ and $T_F$ samples from each library, were PCR-amplified using primers listed in Table S6 and Figure S8 (C–F). The purified amplicons were coupled to Illumina adapters via *Bsa*I-based Golden Gate assembly (NEB catalog no. E1601S). Fully assembled products were then amplified (PCR1) with primers OR1774 and OR1775 (Table S6) to increase DNA yield. Resulting amplicons were diluted to isolate approximately 50,000 individual dsDNA molecules, which were subsequently re-amplified (PCR2) with the same primers to generate multiple copies of each selected molecule, each harboring a unique pair of unique molecular identifiers (UMIs) that traced back to the original molecule. PCR1 and PCR2 were performed in 50 µL reactions containing 1× Q5 buffer, 0.3 mM dNTPs, 500 nM primers, and 1 U Q5 polymerase, with identical thermocycling conditions except for the number of cycles (N = 32 and 50, respectively): 98 °C for 1 min; N cycles of 98 °C for 10 s, 65 °C for 20 s, 72 °C for 40 s; and a final extension at 72 °C for 3 min. PCR2 products were directly subjected to Illumina sequencing (NovaSeq 6000). Sequencing data (≥1.5 Gb) were processed using custom Python scripts and JupyterLab notebooks to generate a ranking of most perturbed positions (positions showing the most deleterious impact on the interaction, considering the combined effect of all mutations). Perturbation scores for each position were calculated as the mean of individual mutation scores (IMS), where IMS was defined as 1 minus the normalized enrichment of the given mutant, adjusted by subtracting the experimental standard deviation. Structural mapping of perturbation scores was performed on the ASF1 protein.

$$Pscore = \frac{1}{n}\sum_{k=1}^{n} ims(k)$$

*Pscore* = position-associated score (average of individual mutation scores for the position)
*n* = number of mutations at a given position
*k* = index of the current mutation for the position

Given that most mutations at the complex interface positions are anticipated to negatively impact affinity and prevent potential compensatory effects in the perturbation score for each position, we we focused on loss-of-interaction (LoI) mutations. To this end, enrichment values exceeding 1−experimental standard deviation were defined as zero.

$$ims(k) = \big(1 - sd - MEnrich(k)\big) \geq 0$$

*ims* = individual mutation score
*sd* = standard deviation of the experiment
*MErinch(k)* = normalized mean enrichment of mutation *k*

#### 1.4. Design, library construction, and selection of binders

##### *1.4.1. Binder design*

N-terminal residues of ip4 were deleted or mutated using Pymol based on the ASF1— ip4 crystal structure (PDB: 6F0H) to comply with different linker sequences (GSEAK, GSEK, GSK; where K corresponds to the 5th ip4 position in the PDB file). Mutated residues were minimized using Chimera version 1.17.3 (100 steps of steepest descent; ten steps of conjugate gradients; Amber ff14SB) and the resulting models were used as input for the design of N-terminal extensions. RFdiffusion[5] was used to generate 50 backbone conformation models for each linker and N-terminal extension length (12, 14, and 16 residues), resulting in a total of 3 linkers × 3 extension lengths × 50 = 450 structures. These structures were clustered using a 2.5 Å RMSD distance threshold calculated using the McLachlan algorithm[6] as implemented in Profit v3.3 (http://www.bioinf.org.uk/software/profit/). Clusters containing more than six models were selected, yielding 15 representative structural models out of 450 (four with 12-residue extensions, six with 14-residue extensions, and five with 16-residue extensions). Native side chains were reintroduced into the non-designed regions of each of the 15 RFdiffusion-derived scaffolds, and 1,000 sequence designs for the N-terminal region were generated using ProteinMPNN[7]. Each designed sequence was then used as input for an AlphaFold2 multimer simulation[8], built from an alignment of Asf1 homologs concatenated with the peptide as a single sequence. The ColabFold v1.5.2[9] implementation (commit 22b5bcf) was used to run the AlphaFold2 structure predictions, generating a single model for each of the 15,000 ProteinMPNN-designed sequences.

The resulting models (1,000 for each of the 15 representative scaffolds) were evaluated according to three criteria: (i) the the root mean square deviation (RMSD) between the predicted model and the original scaffold structure, (ii) the predicted local distance difference test (pLDDT) of the peptide in the designed region, and (iii) the number of contacts between the peptide and ASF1. Of the 15 scaffolds, only five produced solutions with RMSD values below 5 Å relative to the original scaffold. In total, 686 sequences were selected that showed a significant number of atomic contacts, a pLDDT above 50, and RMSDs below 5 Å when superimposing Asf1 and calculating the RMSD between the peptide extensions and the reference scaffold using Profit v3.3. These sequences were introduced into the library for further experimental screening. Over 100 rational designs and three negative controls (flexible linkers) were added to the designs generated by the artificial intelligence (AI) pipeline described above.

##### *1.4.2. Library construction*

The final library set, designed to be Golden Gate compliant, was ordered from IDT as oligonucleotide pools (oPools™, OR2928). dsDNA fragments were generated by PCR filling using a reverse primer (OR2933, Table S6).

In parallel, *Bsa*I-based Golden Gate acceptor cassettes were produced by PCR filling using one oligonucleotide from OR2880–OR2882 with OR2883 (Table S6). The resulting dsDNA products, corresponding to different linkers (GSEAK, GSEK, GSK, respectively) fused to a low-affinity ip4 variant (ip4mutG; qB2H signal between ip3_mut3A and ip1, $K_d \sim 10$ µM), were cloned using Gibson assembly (NEBuilder HiFi) into VN1263 plasmid (v4 series) previously digested by *Not*I and *Bam*HI, resulting in the Golden Gate vectors: VN1925, VN1937, and VN1938, respectively. Therefore, the ip3-coding sequence in VN1263 was replaced in the resulting plasmids with one of the linker-ip4mutG fusions.

The dsDNA library product was inserted into each of the above vectors by Golden Gate cloning, the reaction product was purified by standard ethanol precipitation and resolubilized in molecular biology grade water. Next, the products were used to electroporate SB39 cells prepared at room temperature[10]. After recovery, a small fraction of the culture was used to assure that the expected molecular diversity was covered at least 100X by colony counting on LB-agar plates supplemented with 34 µg/mL chloramphenicol. The remaining culture (sublibraries with different linkers) was incubated overnight (200 rpm, 37 °C, 34 µg/mL chloramphenicol in LB), diluted in the same medium, and bacterial cell populations were mixed equiproportionally at log growth phase ($OD_{600} \sim 0.5$), grown until $OD_{600}$ reached approximately 3.0 and used for glycerol stock preparation. Full library stocks were stored at -80 °C until use.

##### *1.4.3. Short selection runs (batches)*

Glycerol stocks of mixed libraries were inoculated at approximately 18:00 in fresh media and grown overnight at 30 °C, to slow growth kinetics and reduce saturation-related stress (by decreasing the time spent in stationary phase). The grown cultures were diluted to $OD_{600} = 0.05$ in 150 mL of LB supplemented with 17 µg/mL chloramphenicol, 200 µM IPTG, 200 ng/mL aTc, and incubated (200 rpm, 37 °C) for 2 h ($OD_{600}$ ~ 0.5). Then, 100 mL of the culture was used for plasmid miniprep ($T_0$ sample), and the remaining culture was used to inoculate another 100 mL of the same inducing media with varying concentrations of kanamycin: 0, 52, 87, 122, 174, 261 or 348 µg/mL (corresponding to 0, 30, 50, 70, 100, 150, or 200% of ASF1B–ip3 $MIC_{50}$ in the context of the same plasmid series, v4). Assuming that sufficiently strong selection pressure can impact population growth over a short term (few hours), only cultures with OD at least 0.5 absorbance units lower than the OD of the culture without kanamycin after 4.5–5 h were retained for NGS analysis ($T_{F\text{-}[\,X\,\mu g/mL\,kan]}$ samples).

##### *1.4.4. Long selection runs (continuous cultures under stable cell density—turbidostat)*

Culture conditions before kanamycin-based selection were identical to those in "*short selection runs in batches"* including $T_0$ sample preparation. However, the turbidostat function of Chi.Bio system[11,12] was used to implement a continuous culture over extended selection periods. Selection was performed in inducing medium containing kanamycin (209 µg/mL; corresponding to 120% of the ASF1B–ip3 $MIC_{50}$) under the following conditions: starting $OD_{600}$ = 0.05; temperature = 37 °C; stirring rate = 0.8; target OD control = 0.4. Samples (10 mL) were recovered at 21 and 29 h after selection initiation and cultured for an additional 3 hours (final OD ~ 2.5) totalizing, respectively, 24 ($T_{F\_24h}$) or 32 hours ($T_{F\_32h}$) selection.

##### *1.4.5. NGS sample preparation and analysis*

NGS samples were prepared by PCR using primer OR2895 with one of primers from OR2890– OR2894: 98 °C for 1 min; 26 cycles of (98 °C for 15 s.; 65 °C for 15 s.; 72 °C for 1 min) and 72 °C for 3 min. FASTQ-formatted assembled paired-end reads corresponding to the expected sequences were counted and their frequencies calculated. Enrichment of each variant was obtained by the $T_F/T_0$ frequency ratio.

##### *1.4.6. Comparison of samples submitted to different selection pressures*

Pearson correlation coefficient was calculated between pairs of samples using the module `stats` from SciPy (`pearsonr` function) considering only variants observed in all conditions (*n* = 734). The distance between two samples was calculated by subtracting the Pearson correlation coefficient from 1 (i.e., a Pearson correlation coefficient of 1 corresponds to a distance of 0). All pairwise distances were compiled as a distance matrix that was used to infer a phylogenetic tree by neighbor-joining using T-REX server (http://www.trex.uqam.ca). Finally, the unrooted ree was drawn using itol server (https://itol.embl.de).

#### 1.5. Calculation of the required amount of gygabases and NGS data treatment

The requested amount gygabases (Gb) for Illumina NGS sequencing was calculated using the following formula:

$$A = \frac{Ndivxdxrlxrpc}{10^9}$$

$A$ = required amount in gygabases (Gb)
$Ndiv$ = estimated nucleotide diversity
$d$ = sequencing depth (d = 100 in the present work)
$rl$ = read length
rpc = number of reads per cluster (2 for paired-end reads)

NGS data was systematic paired using PEAR[13].

### 2. Tables (description of Excel files)

#### 2.1. Table S1: ASF1–peptide interactions

Interaction strength ($K_d$, calculated by ITC[2]) between $ASF1A_N$ or $ASF1B_N$ and different peptides.

#### Table S2: Promoters and plasmids

Contains data such as name, features, replication origin, corresponding antibiotics and inducers with used concentrations, and library related information.

#### Table S3: qB2H engineering flow cytometry data

Raw and normalized mean fluorescence intensity (MFI) data obtained for different qB2H plasmid systems with statistical results.

#### Table S4: qB2H engineering minimal inhibitory concentration ($MIC_{50}$) data

Raw and normalized $MIC_{50}$ (minimum inhibitory concentration) data obtained for different qB2H plasmid systems with statistical results.

#### Table S5: Stochasticity data

Results related to the comparison of variables (strain, system version, selection pressure, or outlier removal) on key features of the qB2H system (correlation between affinity and enrichment; homogeneity of data values).

#### Table S6: Oligonucleotides

Contains data such as name, sequence, template, destination vector for PCR products, description, and special features concerning oligonucleotides used in this publication.

#### Table S7: Twist library for interface mapping screening

Information about the positional library of point mutants ordered from Twist Bioscience.

#### Table S8: Fitness data from interface mapping experiment

Fitness values expressed as variant enrichment normalized to wild-type.

#### Table S9: Perturbation data from interface mapping experiment

Perturbation values for variants.

#### Table S10: Binders' selection data

Variant description and information such as count, frequency, and when relevant, enrichment values. Pearson correlation analyses between samples are also included.

#### Table S11: Strain description

Strain-related genotypes and features

#### Table S12: Results from additional tests

Analysis of additional model complexes and the influence of induction conditions on immunoglobulin G (IgG) CH3 domain signal output.

### 3. Figures

A

**CONSTRUCTION OF PLASMID VERSION (Plasmid genealogy)**

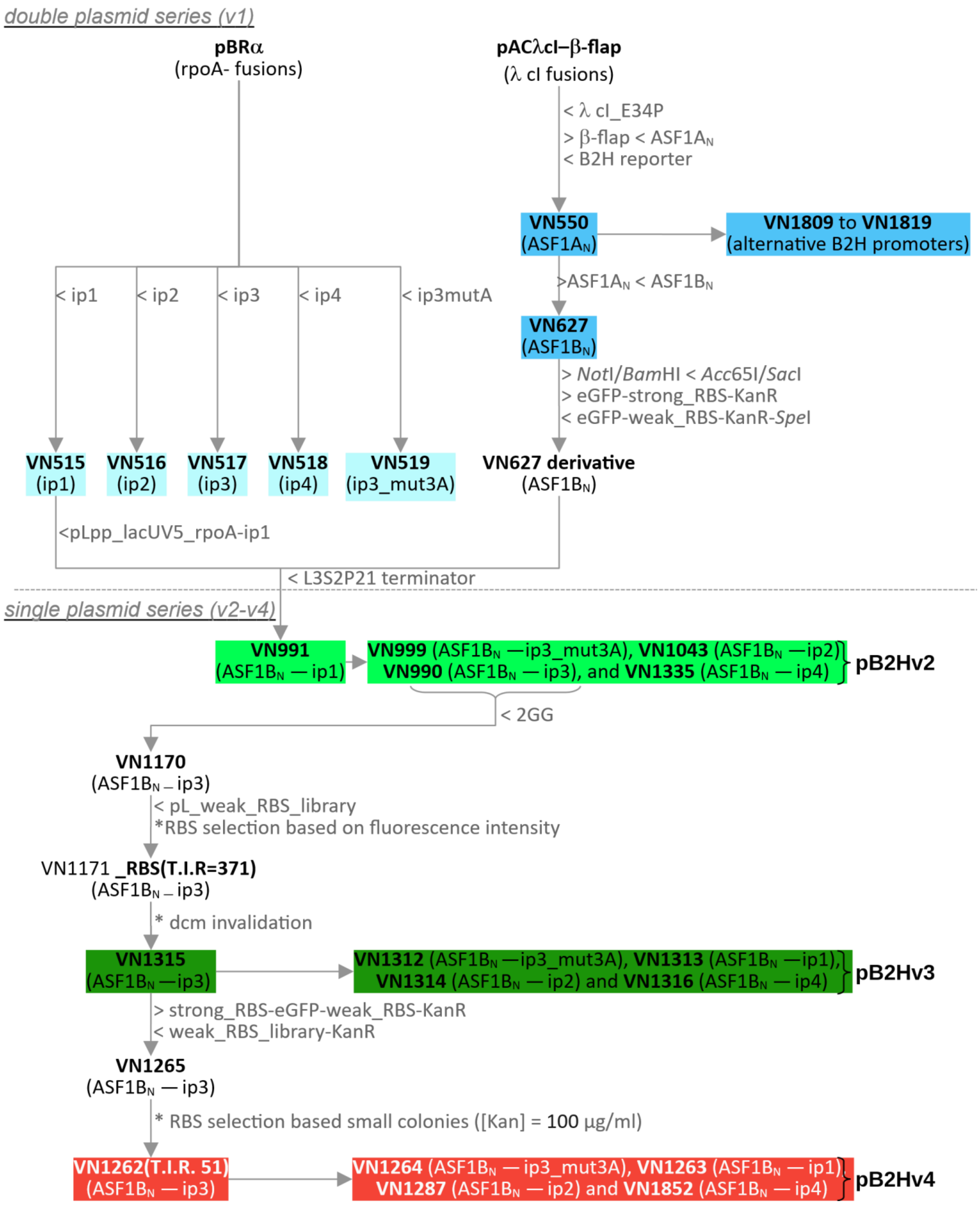

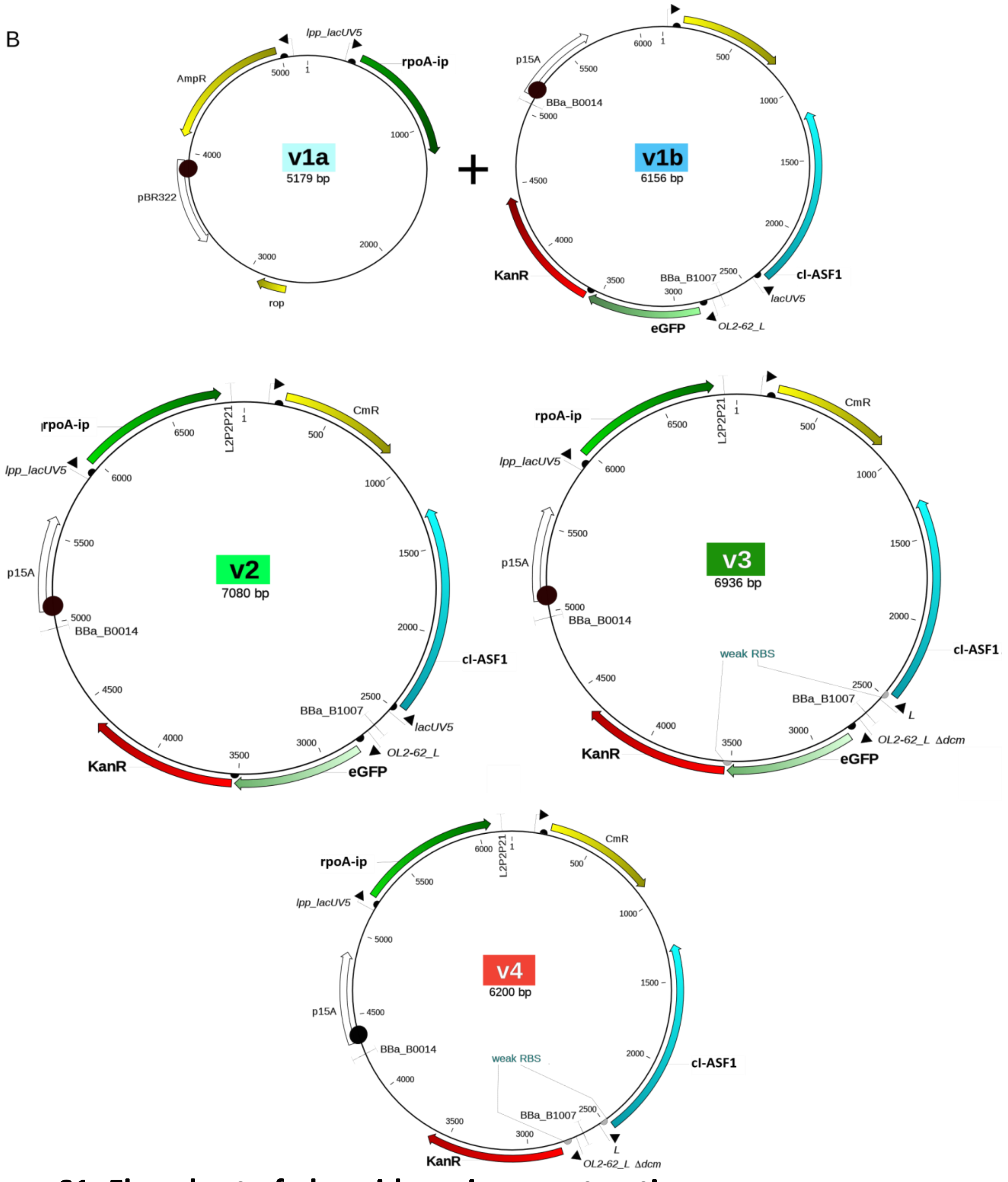

### Figure S1: Flowchart of plasmid version construction

**(A)** Flowchart representing relationships among the plasmids constructed in this work. Interaction partners are indicated in parentheses. "**>**" indicates that the element was removed from the ancestral plasmid, and "**<**" indicates that it was inserted. Plasmids from v1 (pBR$\alpha$: pB2H v1a; pAC-cI_E34P: pB2H v1b), v2 (pB2H v2), v3 (pB2H v3), and v4 (pB2H v4) series are shown in different background colors. B2H reference reporter: "BBa_B1007_*Pac*I-OL2-62_L-*Bsi*WI_RBS_eGFP-RBS-KanR_f1_BBa_B0014"; 2GG: DNA cassette intended for Golden Gate cloning. **(B)** Main features of each quantitative bacterial two-hybrid plasmid series are indicated in the plasmid maps.

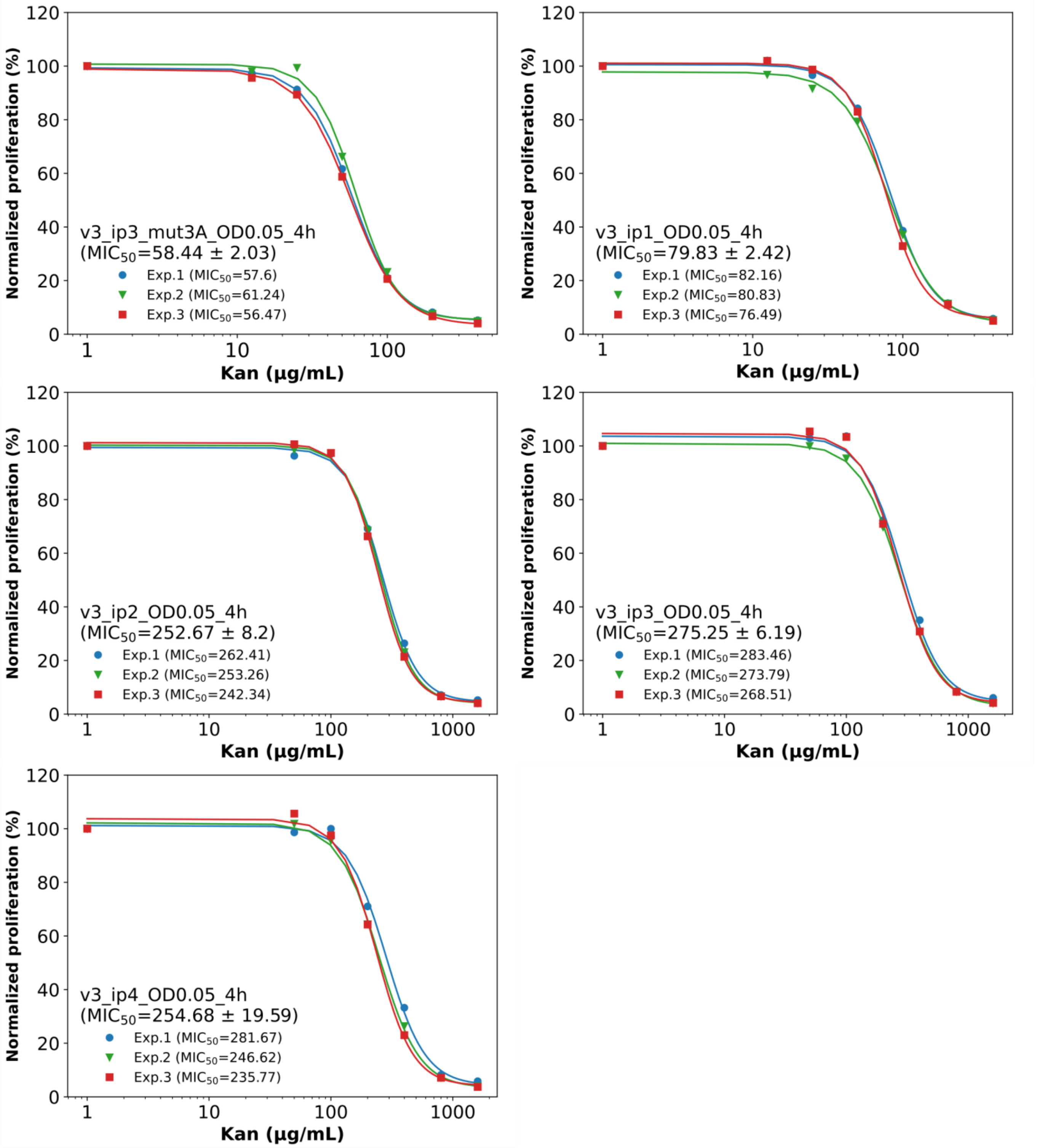

#### Figure S2: Effect of kanamycin concentration on proliferation of SB39 pB2H v3 (starting OD = 0.05, time = 4 h)

The following v3 series plasmids were analyzed: VN1312 (cI-ASF1B — rpoA-ip3_mut3A, $K_d$ > 100 µM), VN1313 (cI-ASF1B — rpoA-ip1, $K_d$ = 5 300 nM), VN1314 (cI-ASF1B — rpoA-ip2, $K_d$ = 310 nM), VN1315 (cI-ASF1B — rpoA-ip3, $K_d$ = 93 nM), and VN1316 (cI-ASF1B — rpoA-ip4, $K_d$ = 2 nM). Weak interactions (VN1312 and VN1313) were evaluated at 0, 12.5, 25, 50, 100, 200 and 400 µg/mL kanamycin while strong interactions (VN1314–VN1316) were evaluated at 0, 50, 100, 200, 400, 800, and 1,600 µg/mL kanamycin.

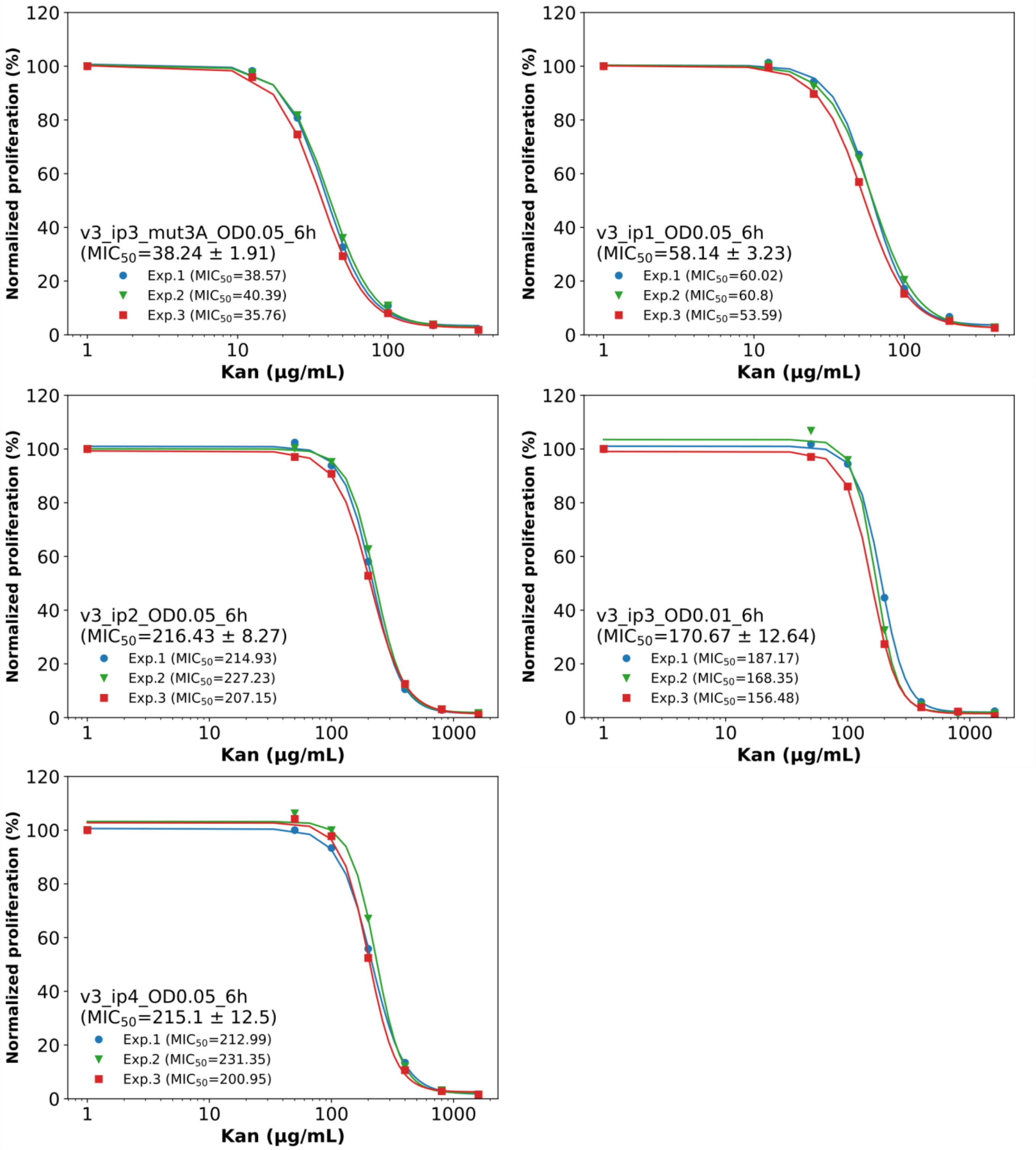

#### Figure S3: Effect of kanamycin concentration on proliferation of SB39 pB2H v3 (starting OD = 0.05, time = 6 h)

The following v3 series plasmids were analyzed: VN1312 (cI-ASF1B — rpoA-ip3_mut3A, $K_d$ > 100 µM), VN1313 (cI-ASF1B — rpoA-ip1, $K_d$ = 5 300 nM), VN1314 (cI-ASF1B — rpoA-ip2, $K_d$ = 310 nM), VN1315 (cI-ASF1B — rpoA-ip3, $K_d$ = 93 nM), and VN1316 (cI-ASF1B — rpoA-ip4, $K_d$ = 2 nM). Weak interactions (VN1312 and VN1313) were evaluated at 0, 12.5, 25, 50, 100, 200, and 400 µg/mL kanamycin while strong interactions (VN1314–VN1316) were evaluated at 0, 50, 100, 200, 400, 800, and 1,600 µg/mL kanamycin.

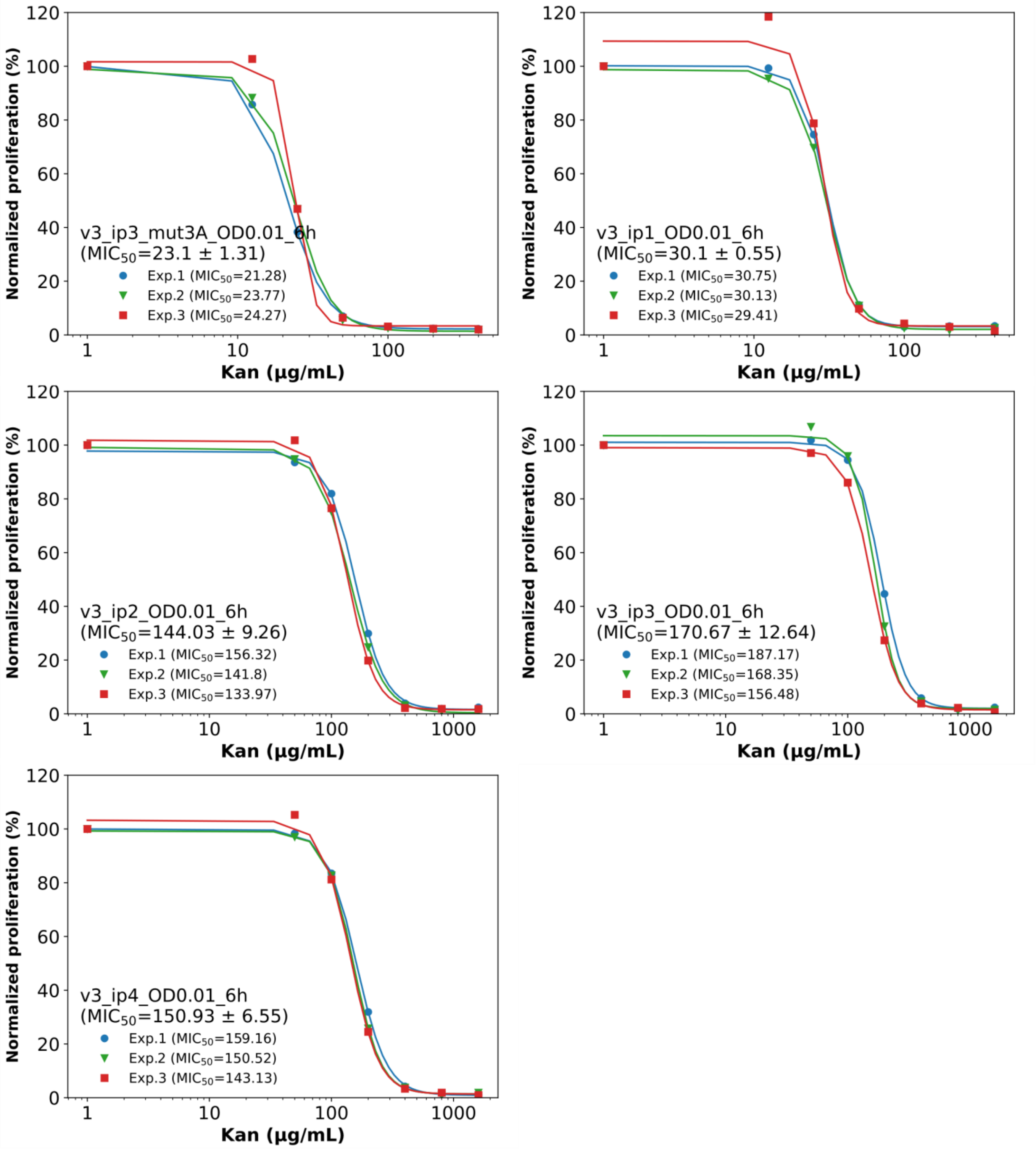

### Figure S4: Effect of kanamycin concentration on proliferation of SB39 pB2H v3 (starting OD = 0.01, time = 6 h)

The following v3 series plasmids were analyzed: VN1312 (cI-ASF1B — rpoA-ip3_mut3A, $K_d$ > 100 µM), VN1313 (cI-ASF1B — rpoA-ip1, $K_d$ = 5 300 nM), VN1314 (cI-ASF1B — rpoA-ip2, $K_d$ = 310 nM), VN1315 (cI-ASF1B — rpoA-ip3, $K_d$ = 93 nM), and VN1316 (cI-ASF1B — rpoA-ip4, $K_d$ = 2 nM). Weak interactions (VN1312 and VN1313) were evaluated at 0, 12.5, 25, 50, 100, 200, and 400 µg/mL kanamycin while strong interactions (VN1314–VN1316) were evaluated at 0, 50, 100, 200, 400, 800, and 1,600 µg/mL kanamycin.

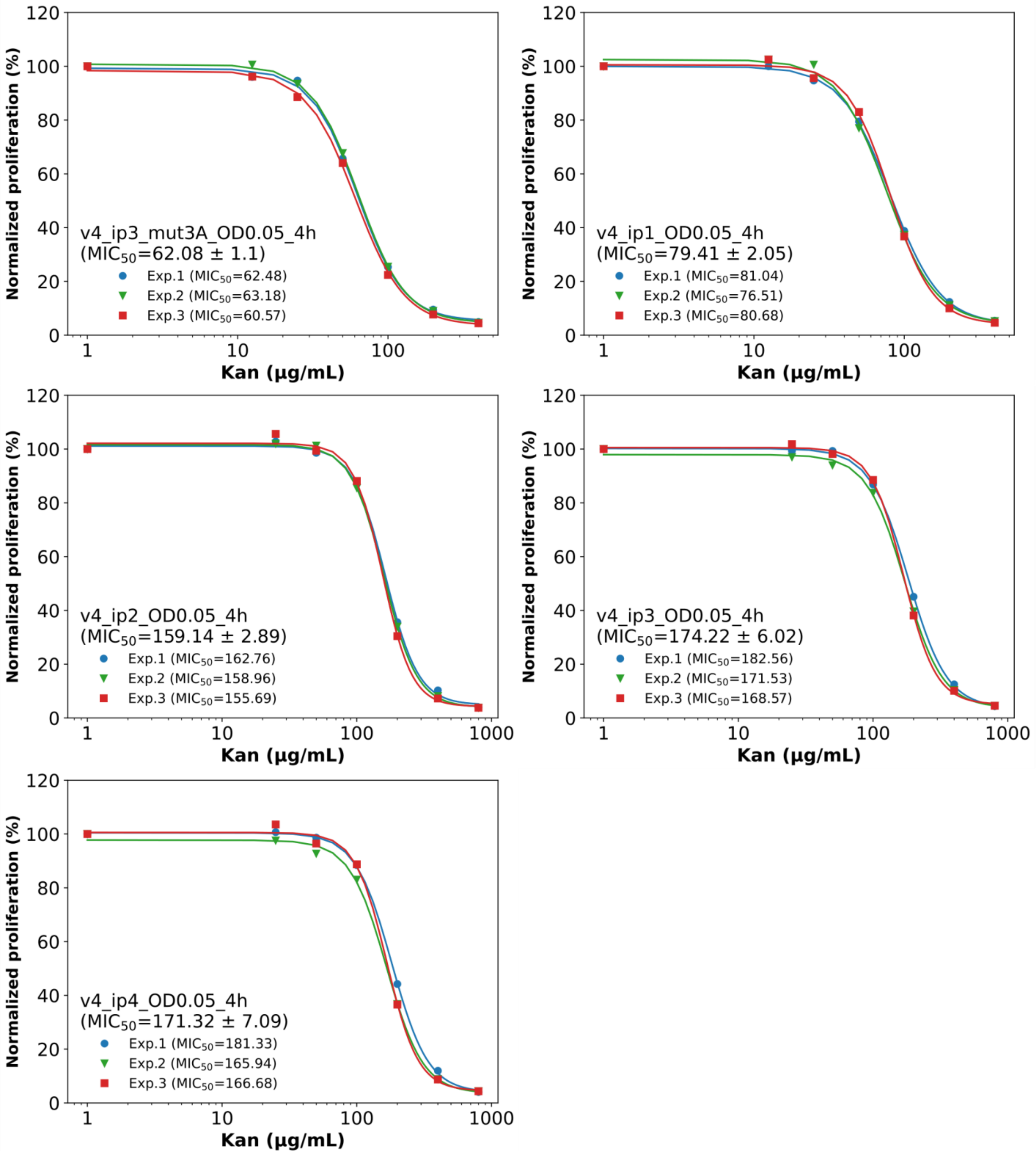

#### Figure S5: Effect of kanamycin concentration on proliferation of SB39 pB2H v4 (starting OD = 0.05, time = 4 h)

The following v4 series plasmids were analyzed: VN1264 (cI-ASF1B — rpoA-ip3_mut3A, $K_d$ > 100 µM), VN1263 (cI-ASF1B — rpoA-ip1, $K_d$ = 5 300 nM), VN1287 (cI-ASF1B — rpoA-ip2, $K_d$ = 310 nM), VN1262 (cI-ASF1B — rpoA-ip3, $K_d$ = 93 nM), and VN1852 (cI-ASF1B — rpoA-ip4, $K_d$ = 2 nM). Weak interactions (VN1264 and VN1263) were evaluated at 0, 12.5, 25, 50, 100, 200, and 400 µg/mL kanamycin, while strong interactions (VN1287, VN1262, and VN1852) were evaluated at 0, 25, 50, 100, 200, 400, and 800 µg/mL kanamycin.

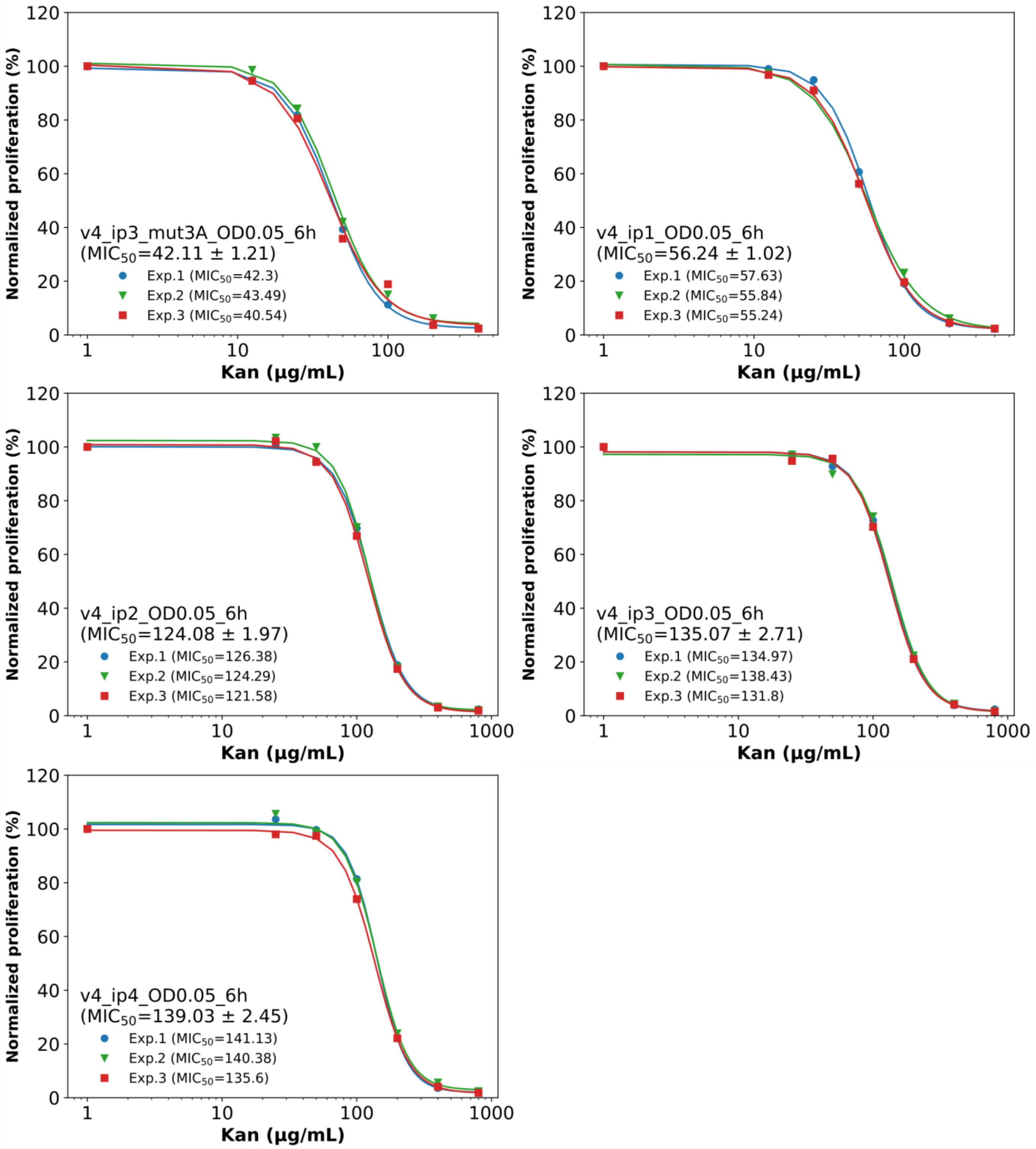

#### Figure S6: Effect of kanamycin concentration on proliferation of SB39 pB2H v4 (starting OD = 0.05, time = 6 h)

The following v4 series plasmids were analyzed: VN1264 (cI-ASF1B — rpoA-ip3_mut3A, $K_d$ > 100 μM), VN1263 (cI-ASF1B — rpoA-ip1, $K_d$ = 5 300 nM), VN1287 (cI-ASF1B — rpoA-ip2, $K_d$ = 310 nM), VN1262 (cI-ASF1B — rpoA-ip3, $K_d$ = 93 nM), and VN1852 (cI-ASF1B — rpoA-ip4, $K_d$ = 2 nM). Weak interactions (VN1264 and VN1263) were evaluated at 0, 12.5, 25, 50, 100, 200, and 400 μg/mL kanamycin, while strong interactions (VN1287, VN1262, and VN1852) were evaluated at 0, 25, 50, 100, 200, 400, and 800 μg/mL kanamycin.

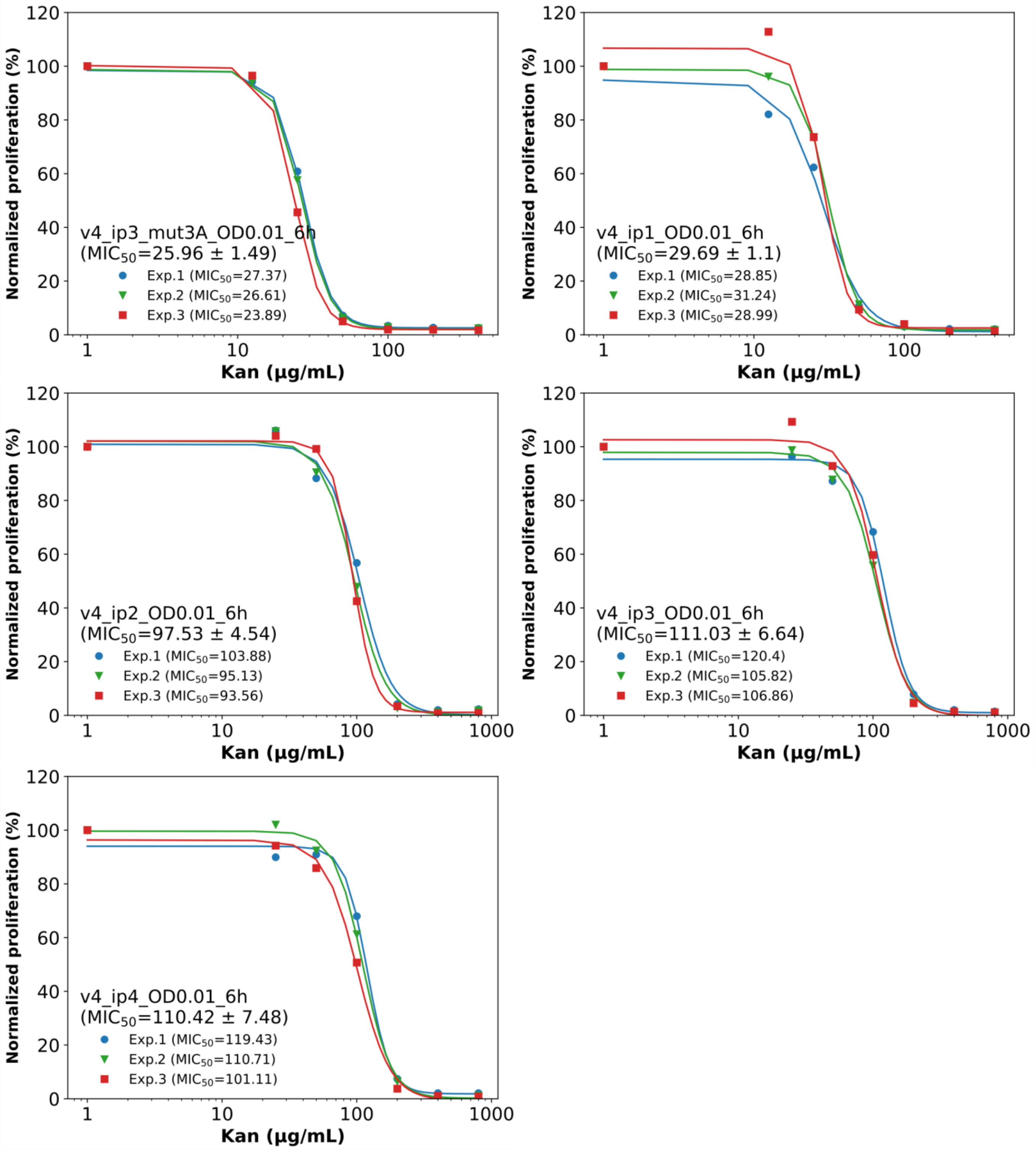

#### Figure S7: Effect of kanamycin concentration on proliferation of SB39 pB2H v4 (starting OD = 0.01, time = 6 h)

The following v4 series plasmids were analyzed: VN1264 (cI-ASF1B — rpoA-ip3_mut3A, $K_d$ > 100 µM), VN1263 (cI-ASF1B — rpoA-ip1, $K_d$ = 5 300 nM), VN1287 (cI-ASF1B — rpoA-ip2, $K_d$ = 310 nM), VN1262 (cI-ASF1B — rpoA-ip3, $K_d$ = 93 nM), and VN1852 (cI-ASF1B — rpoA-ip4, $K_d$ = 2 nM). Weak interactions (VN1264 and VN1263) were evaluated at 0, 12.5, 25, 50, 100, 200, and 400 µg/mL kanamycin, while strong interactions (VN1287, VN1262, and VN1852) were evaluated at 0, 25, 50, 100, 200, 400, and 800 µg/mL kanamycin.

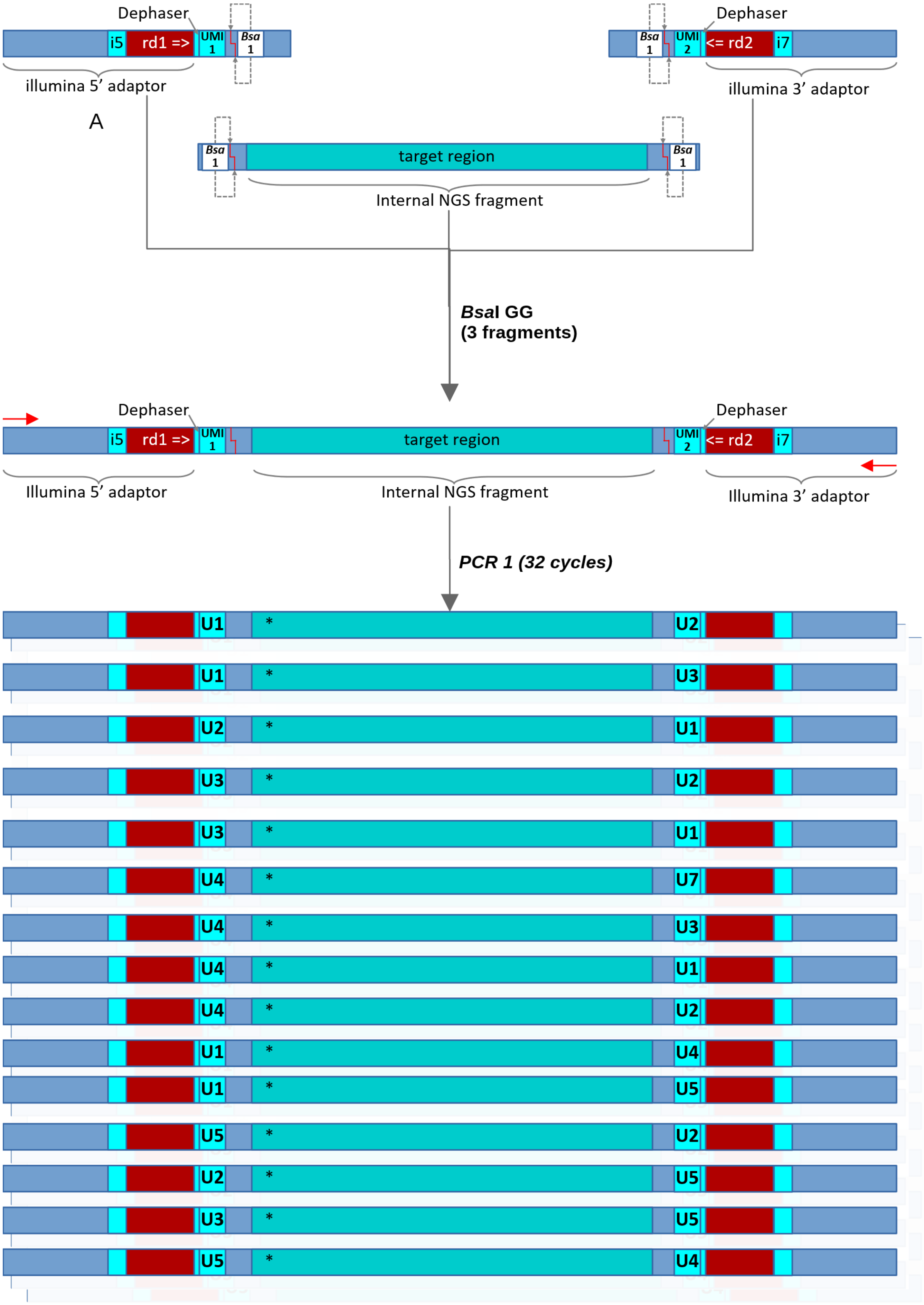
Dephaser
i5
rd1 =>
UMI 1
Bsa 1
illumina 5’ adaptor
Dephaser
Bsa 1
UMI 2
<= rd2
i7
illumina 3’ adaptor
A
Bsa 1
target region
Bsa 1
Internal NGS fragment
BsaI GG
(3 fragments)
Dephaser
i5
rd1 =>
UMI 1
target region
UMI 2
<= rd2
i7
Dephaser
Illumina 5’ adaptor
Internal NGS fragment
Illumina 3’ adaptor
PCR 1 (32 cycles)
U1 * U2
U1 * U3
U2 * U1
U3 * U2
U3 * U1
U4 * U7
U4 * U3
U4 * U1
U4 * U2
U1 * U4
U1 * U5
U5 * U2
U2 * U5
U3 * U5
U5 * U4

B

***PCR 1 product***
***- Dilution to 50 000 molecules -***

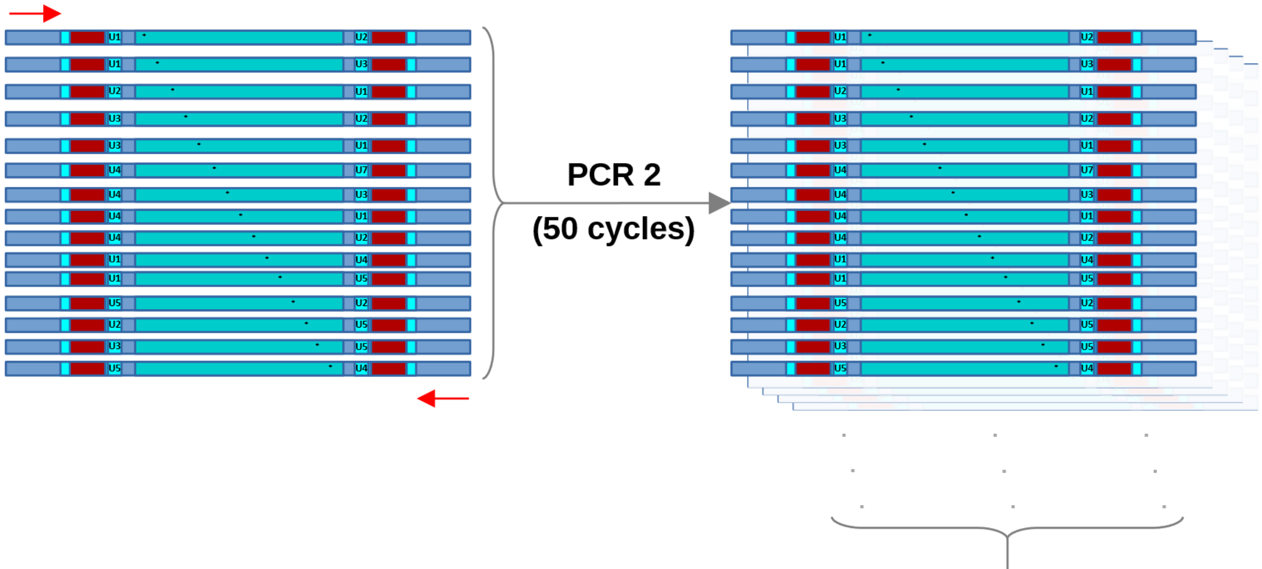

**NGS (Illumina)**

- $Data\ amount\,(Gb) \geq Ndiv \ \ x \ \ d \ \ x \ \ rl\ x \ \ rpc$

Data amount ≥ 50 000 x 100 x 150 x 2

Data amount ≥ 1.5 Gb

*Ndiv = theoretical molecular diversity*
*d = sequencing depth (desired number of Illumina clusters per theoretical molecule)*
*rl = Illumina read length (bases)*
*rpc = number of reads per Illumina cluster (single-read = 1; pair ed-end = 2)*

**Data**

- data treatment (python)
- data analysis (jupyter-lab / python)

**Results**

- **Most perturbed positions** (ranking; threshold = 0.1):

Position : Perturbation
54 : 0.46
110 : 0.36
108 : 0.32
96 : 0.26

*Mapping*

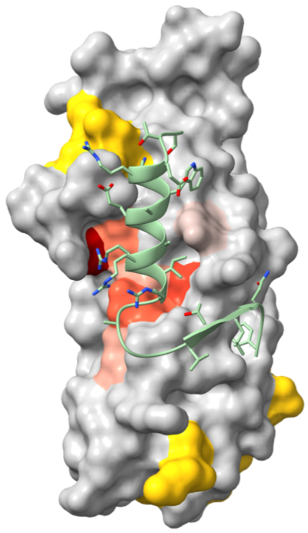

C

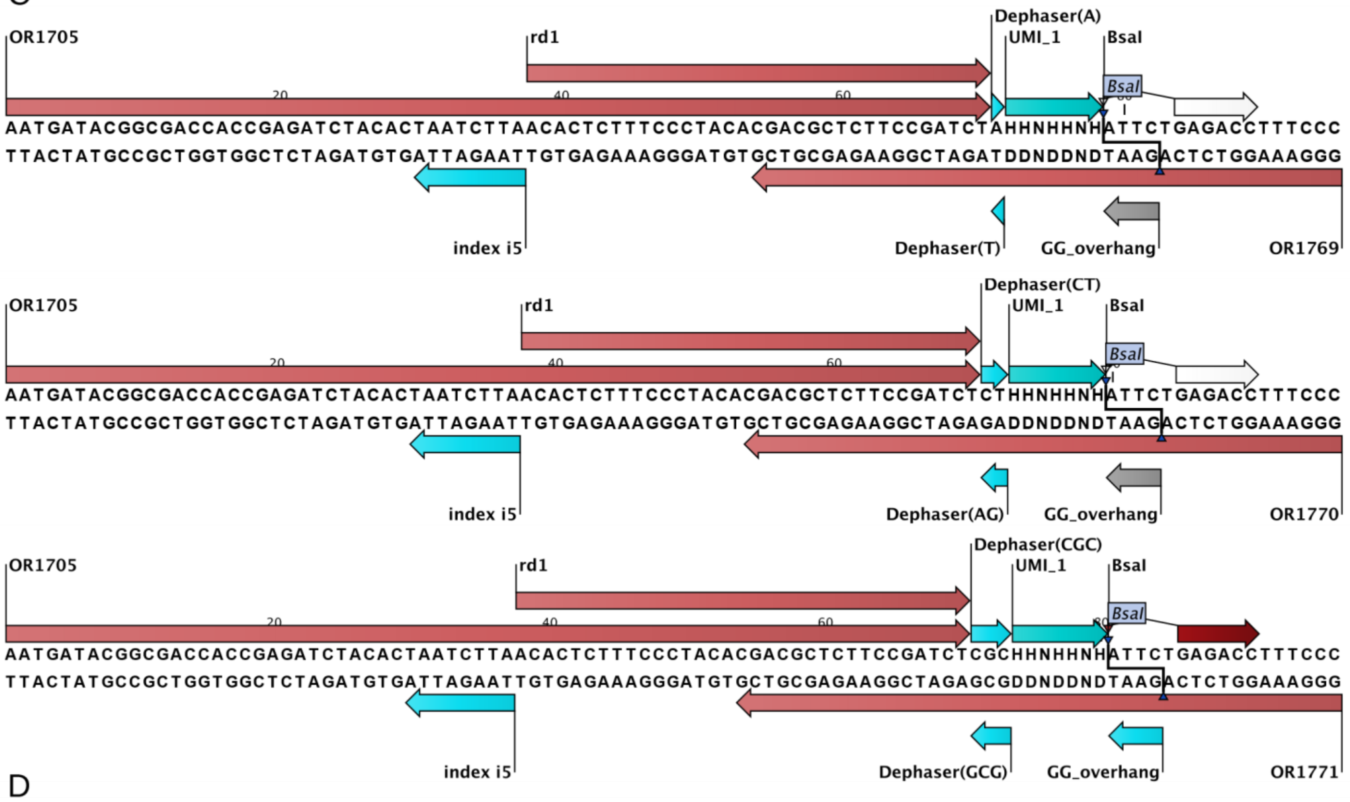

OR1705
rd1
Dephaser(A)
UMI_1
BsaI
AATGATACGGCGACCACCGAGATCTACACTAATCTTAACACTCTTTCCCTACACGACGCTCTTCCGATCTAHHNHHNHATTCTGAGACCTTTCCC
TTACTATGCCGCTGGTGGCTCTAGATGTGATTAGAATTGTGAGAAAGGGATGTGCTGCGAGAAGGCTAGATDDNDDNDTAAGACTCTGGAAAGGG
index i5
Dephaser(T)
GG_overhang
OR1769
OR1705
rd1
Dephaser(CT)
UMI_1
BsaI
AATGATACGGCGACCACCGAGATCTACACTAATCTTAACACTCTTTCCCTACACGACGCTCTTCCGATCTCTHHNHHNHATTCTGAGACCTTTCCC
TTACTATGCCGCTGGTGGCTCTAGATGTGATTAGAATTGTGAGAAAGGGATGTGCTGCGAGAAGGCTAGAGADDNDDNDTAAGACTCTGGAAAGGG
index i5
Dephaser(AG)
GG_overhang
OR1770
OR1705
rd1
Dephaser(CGC)
UMI_1
BsaI
AATGATACGGCGACCACCGAGATCTACACTAATCTTAACACTCTTTCCCTACACGACGCTCTTCCGATCTCGCHHNHHNHATTCTGAGACCTTTCCC
TTACTATGCCGCTGGTGGCTCTAGATGTGATTAGAATTGTGAGAAAGGGATGTGCTGCGAGAAGGCTAGAGCGDDNDDNDTAAGACTCTGGAAAGGG
index i5
Dephaser(GCG)
GG_overhang
OR1771

D

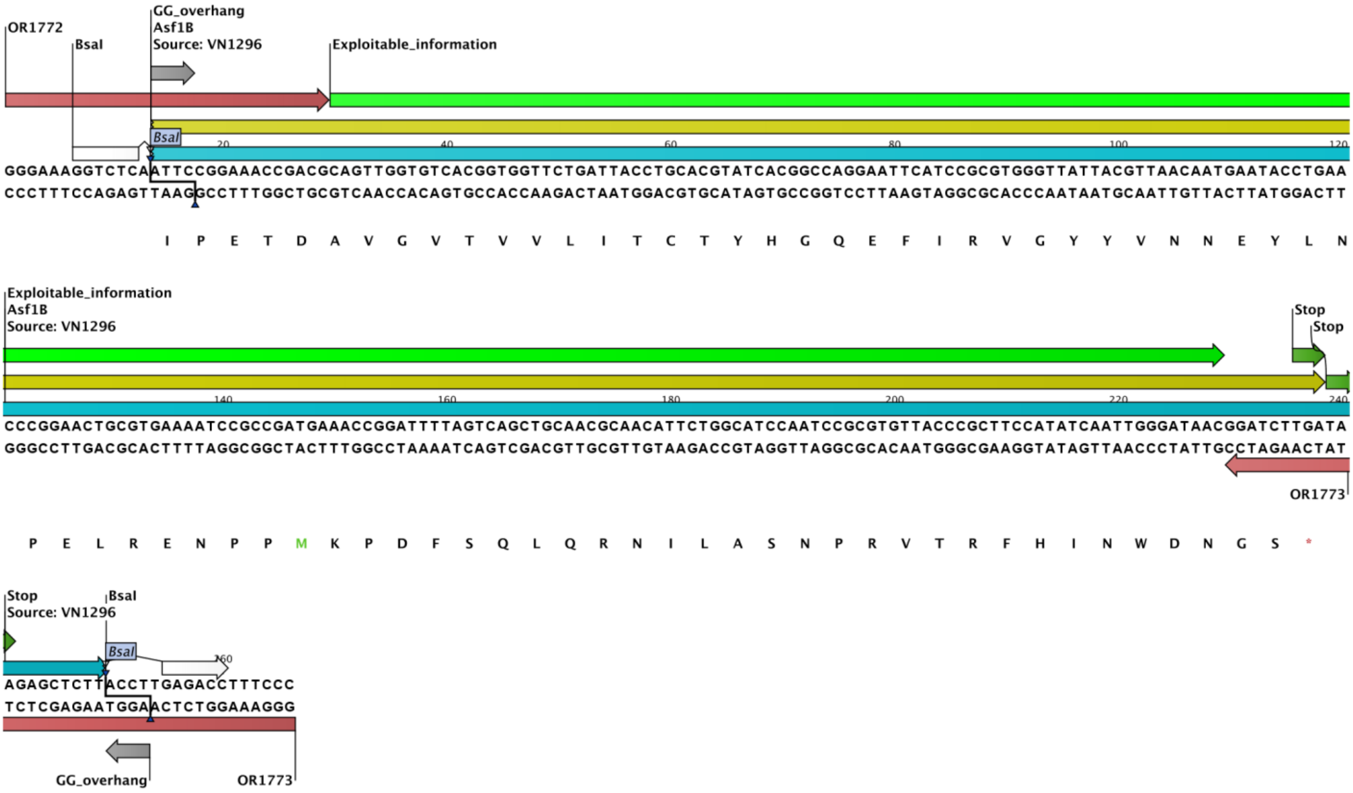

OR1772
BsaI
GG_overhang
Asf1B
Source: VN1296
Exploitable_information
GGGAAAGGTCTCAATTCCGGAAACCGACGCAGTTGGTGTCACGGTGGTTCTGATTACCTGCACGTATCACGGCCAGGAATTCATCCGCGTGGGTTATTACGTTAACAATGAATACCTGAA
CCCTTTCCAGAGTTAAGGCCTTTGGCTGCGTCAACCACAGTGCCACCAAGACTAATGGACGTGCATAGTGCCGGTCCTTAAGTAGGCGCACCCAATAATGCAATTGTTACTTATGGACTT
I P E T D A V G V T V V L I T C T Y H G Q E F I R V G Y Y V N N E Y L N
Exploitable_information
Asf1B
Source: VN1296
Stop
Stop
CCCGGAACTGCGTGAAAATCCGCCGATGAAACCGGATTTTAGTCAGCTGCAACGCAACATTCTGGCATCCAATCCGCGTGTTACCCGCTTCCATATCAATTGGGATAACGGATCTTGATA
GGGCCTTGACGCACTTTTAGGCGGCTACTTTGGCCTAAAATCAGTCGACGTTGCGTTGTAAGACCGTAGGTTAGGCGCACAATGGGCGAAGGTATAGTTAACCCTATTGCCTAGAACTAT
OR1773
P E L R E N P P M K P D F S Q L Q R N I L A S N P R V T R F H I N W D N G S *
Stop
Source: VN1296
BsaI
AGAGCTCTTACCTTGAGACCTTTCCC
TCTCGAGAATGGAACTCTGGAAAGGG
GG_overhang
OR1773

E

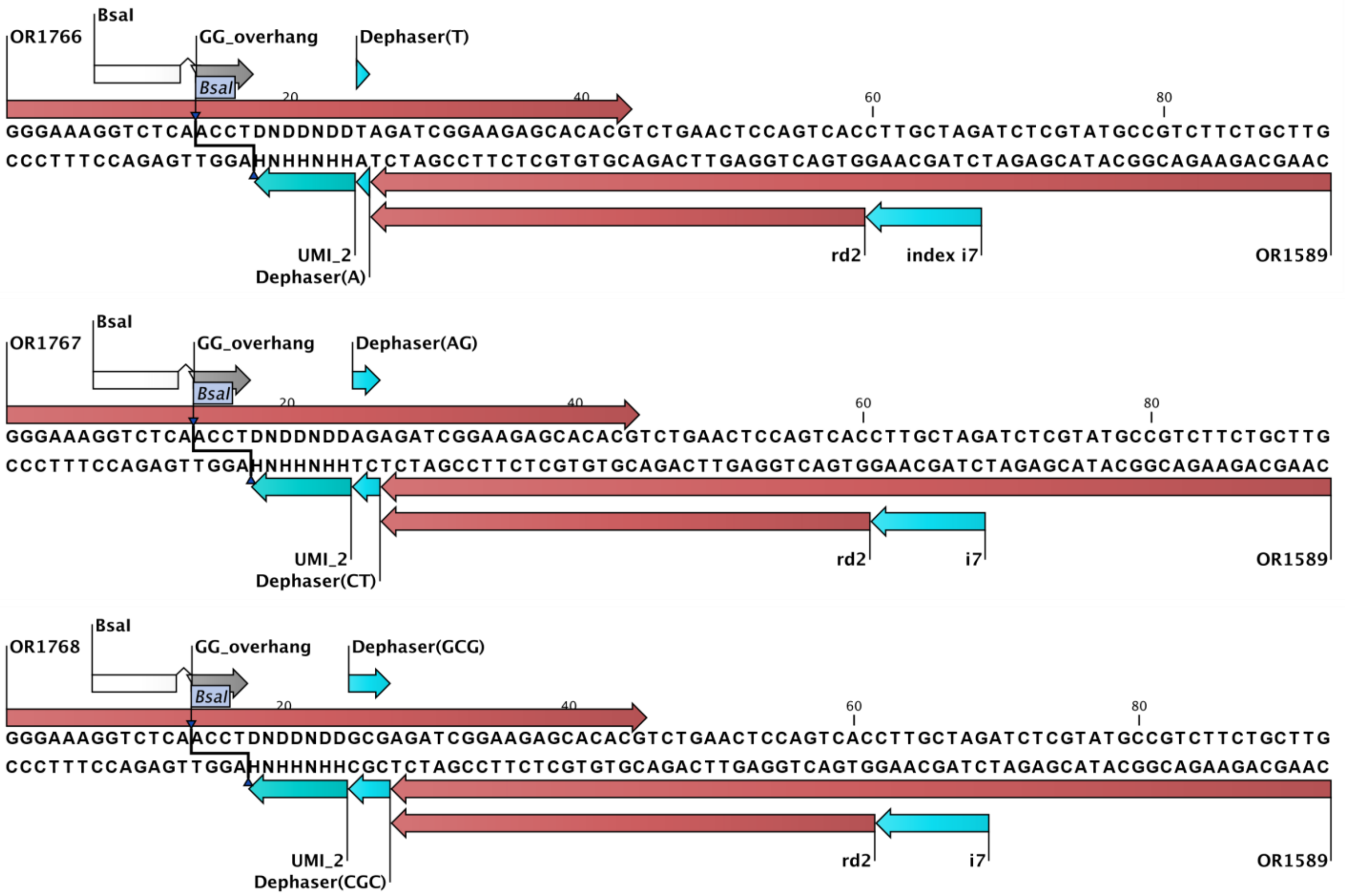
BsaI
OR1766
GG_overhang
Dephaser(T)
BsaI
20
40
60
80
GGGAAAGGTCTCAACCTDNDDNDDTAGATCGGAAGAGCACACGTCTGAACTCCAGTCACCTTGCTAGATCTCGTATGCCGTCTTCTGCTTG
CCCTTTCCAGAGTTGGAHNHHNHHATCTAGCCTTCTCGTGTGCAGACTTGAGGTCAGTGGAACGATCTAGAGCATACGGCAGAAGACGAAC
UMI_2
Dephaser(A)
rd2
index i7
OR1589
BsaI
OR1767
GG_overhang
Dephaser(AG)
BsaI
20
40
60
80
GGGAAAGGTCTCAACCTDNDDNDDAGAGATCGGAAGAGCACACGTCTGAACTCCAGTCACCTTGCTAGATCTCGTATGCCGTCTTCTGCTTG
CCCTTTCCAGAGTTGGAHNHHNHHTCTCTAGCCTTCTCGTGTGCAGACTTGAGGTCAGTGGAACGATCTAGAGCATACGGCAGAAGACGAAC
UMI_2
Dephaser(CT)
rd2
i7
OR1589
BsaI
OR1768
GG_overhang
Dephaser(GCG)
BsaI
20
40
60
80
GGGAAAGGTCTCAACCTDNDDNDDGCGAGATCGGAAGAGCACACGTCTGAACTCCAGTCACCTTGCTAGATCTCGTATGCCGTCTTCTGCTTG
CCCTTTCCAGAGTTGGAHNHHNHHCGCTCTAGCCTTCTCGTGTGCAGACTTGAGGTCAGTGGAACGATCTAGAGCATACGGCAGAAGACGAAC
UMI_2
Dephaser(CGC)
rd2
i7
OR1589

F

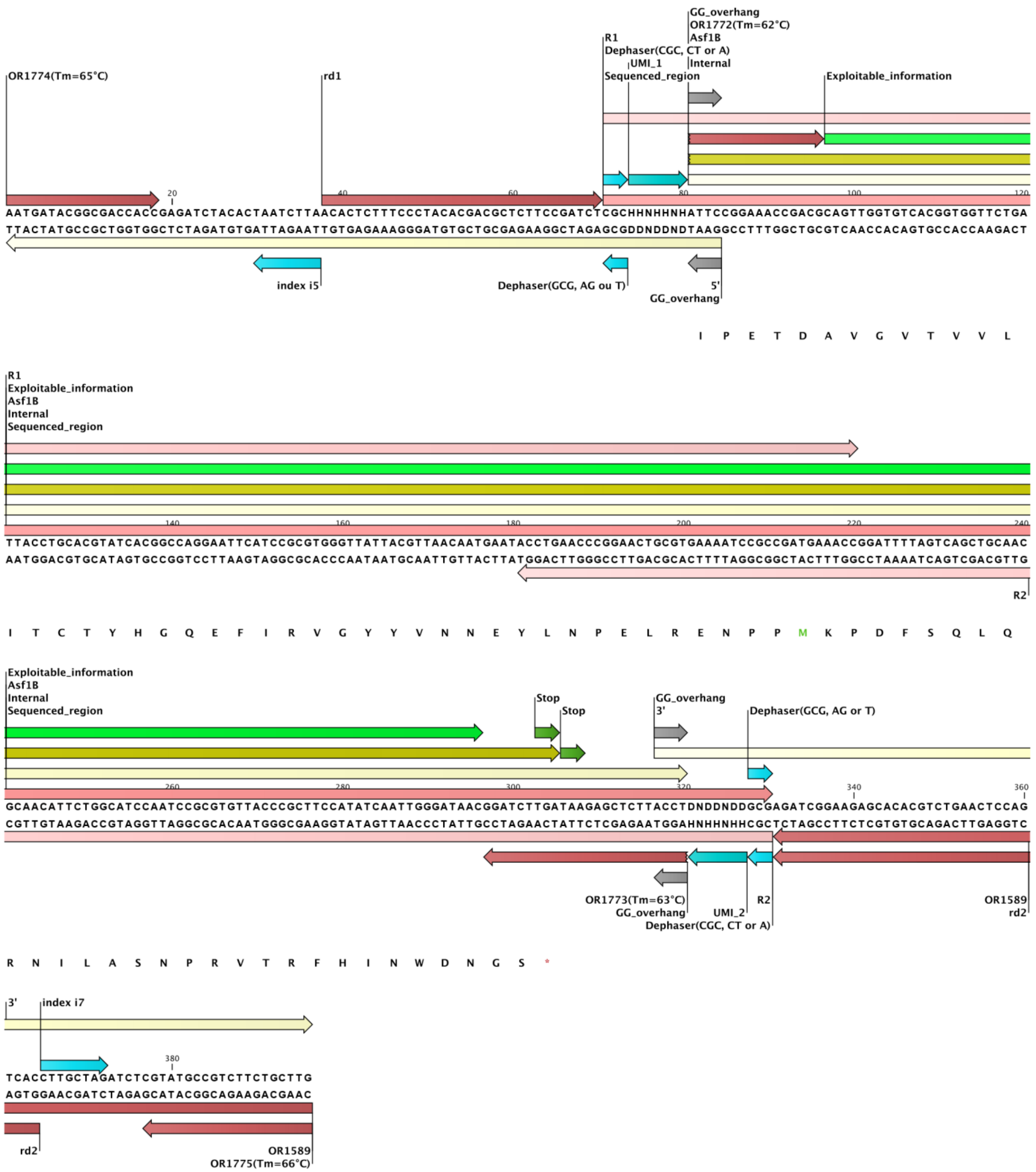

#### Figure S8: Interface mapping strategy

T0 (plasmids extracted before selection) and TF (plasmids extracted after selection) samples were submitted to the interface mapping pipeline, starting with polymerase chain reaction (PCR) amplification of two ASF1 target regions (N-terminus, Middle or C-terminus) using primers that add flanking *Bsa*I recognition sites. These fragments were fused with compatible 5’ and 3’ Illumina adapters by Golden Gate assembly. Each adapter harbors a unique molecular identifier (UMI) with the sequence HHNHHNH; therefore, two UMIs are combined into the final assembly product, providing approximately $1.51 \times 10^7$ theoretical combinations. Adapters also include dephasers (1–3 bases) to improve cluster clonality of low-throughput Illumina sequencing systems (i.g., iSeq100). Assembled products were PCR amplified (PCR 1) to increase the available amount for a reliable quantification **(A)**. Then, amplicons are diluted and approximately 50,000 individual molecules are recovered for the second PCR round (PCR 2), with products directly used as next-generation sequencing (NGS) samples. A sequencing depth of at least 100-fold coverage of the diversity was achieved using ≥1.5 Gb of a 2 × 150bp NovaSeq 6000 run. NGS data were processed using custom Python scripts and analyzed using JupyterLab notebooks, resulting in a ranking of perturbation score for each position above a threshold that was mapped to the target structure **(B)**. Annotated sequences of the 5′ adapter **(C)**, an example of the internal target region fragment **(D)**, the 3′ adapter **(E)**, and the fully assembled product **(F)** are presented to illustrate the Golden Gate assembly approach.

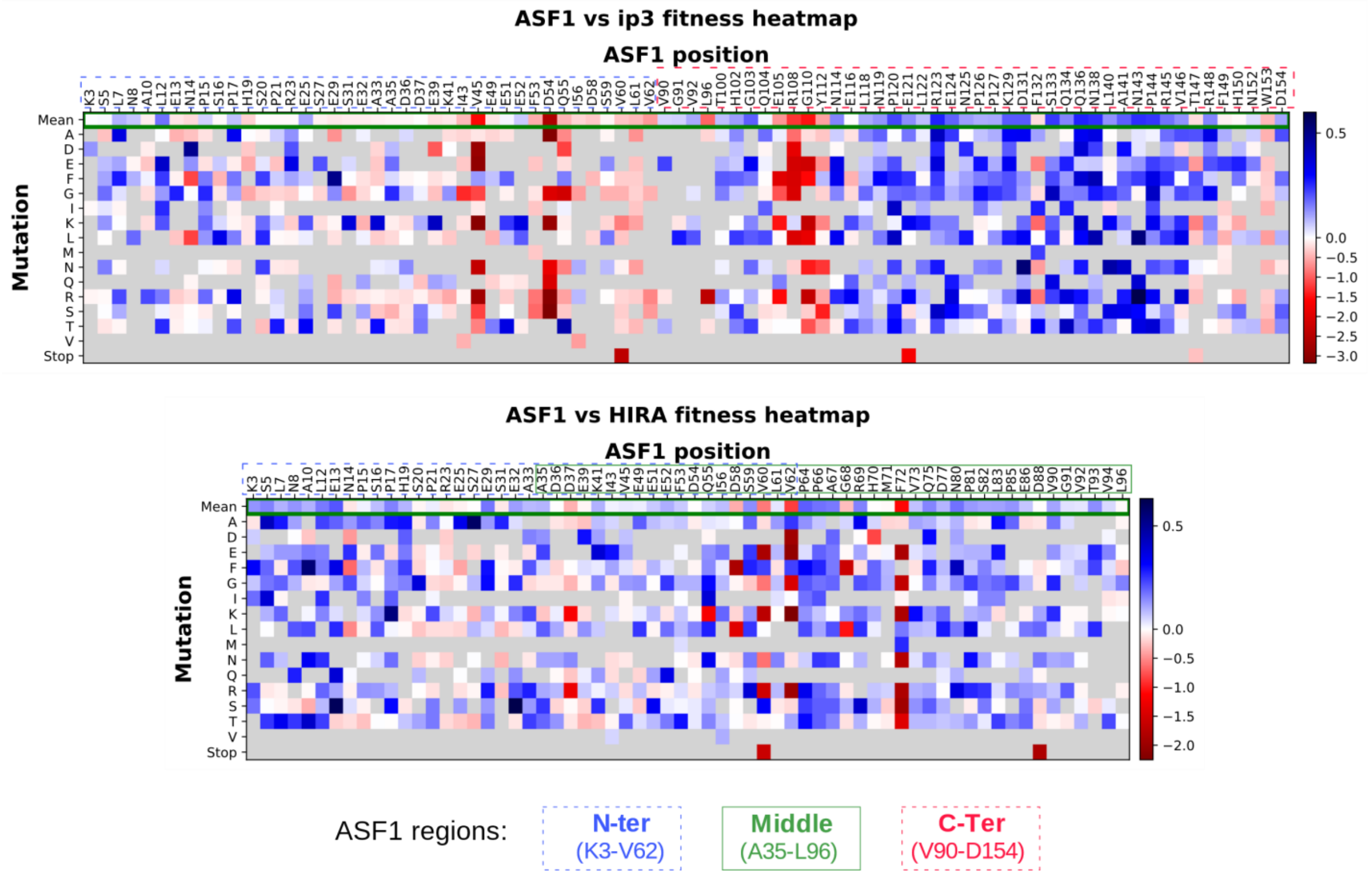

### Figure S9: ASF1B$_N$ MAVE – fitness analysis

Variant fitness is represented as heatmaps of enrichment normalized to wild-type. **(A)** Heatmap for the ASF1B$_N$–ip3 complex. **(B)** Heatmap for the ASF1B$_N$–HIRA complex. Positions corresponding to ASF1 regions (N-terminus, Middle, and C-terminus) are indicated in colored boxes.

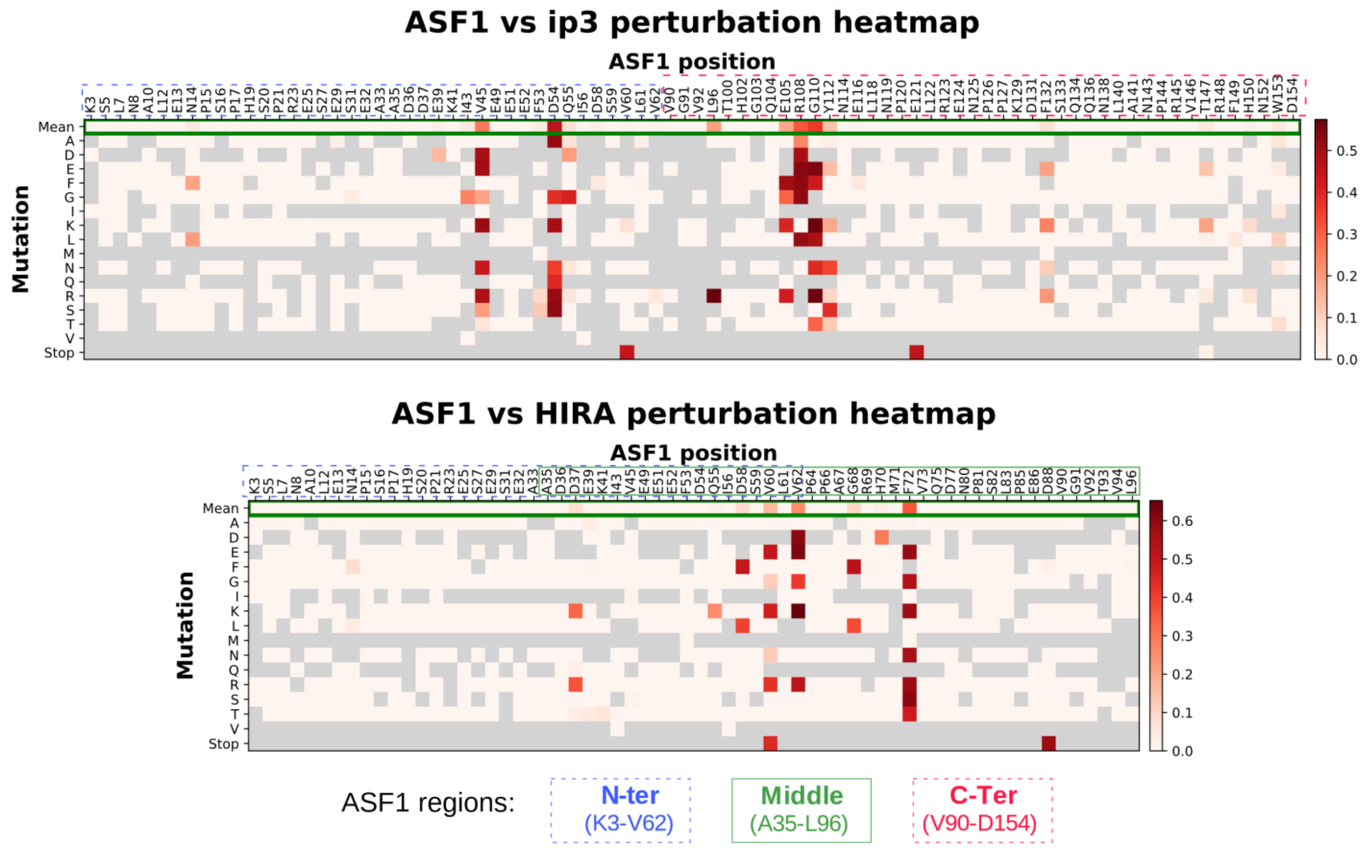

#### Figure S10: ASF1B$_N$ MAVE – perturbation analysis

Variant perturbation scores for loss-of-interaction mutations represented as heatmaps. **(A)** Heatmap for the ASF1B$_N$–ip3 complex. **(B)** Heatmap for the ASF1B$_N$–HIRA complex. Positions corresponding to ASF1 regions (N-terminus, Middle, and C-terminus) are indicated in colored boxes.

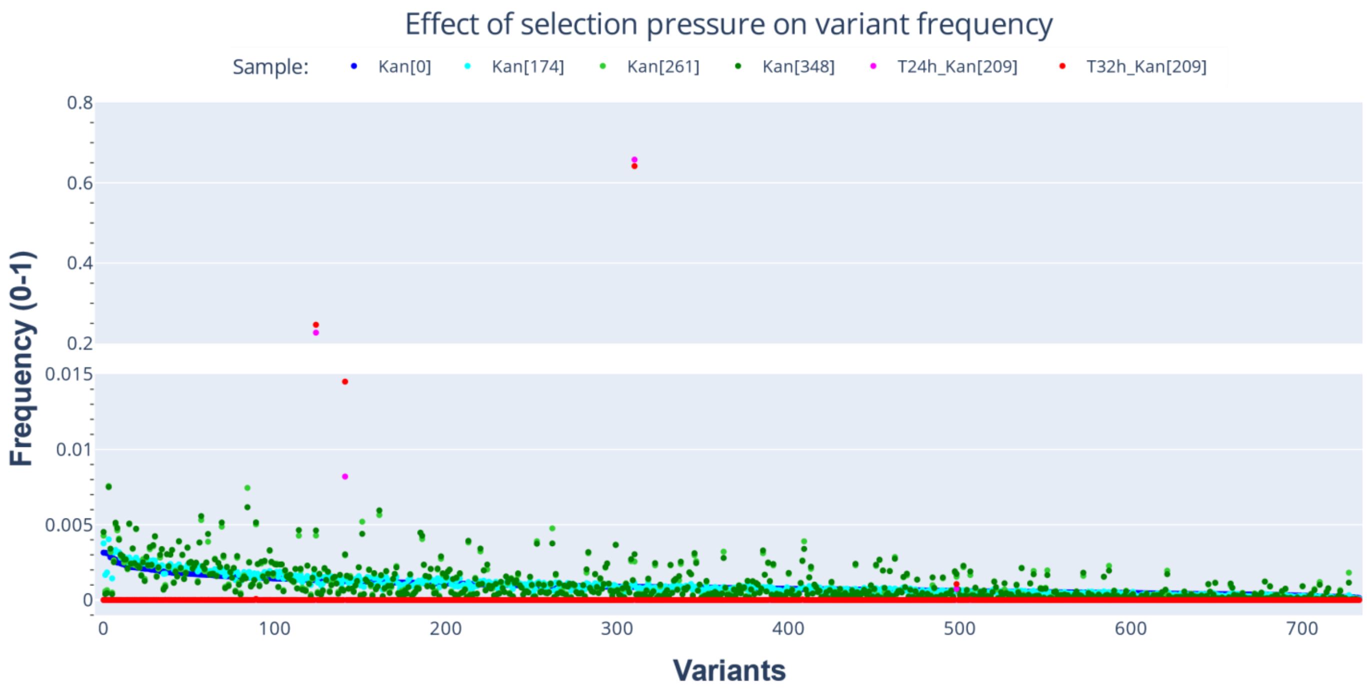

#### Figure S11: Variant frequencies in different binder selection conditions

A total of 734 variants present in all samples are shown. Conditions: Kan[0] (no selection); Kan[174], Kan[261], Kan[348], T24h_Kan[209], and T32h_Kan[209].

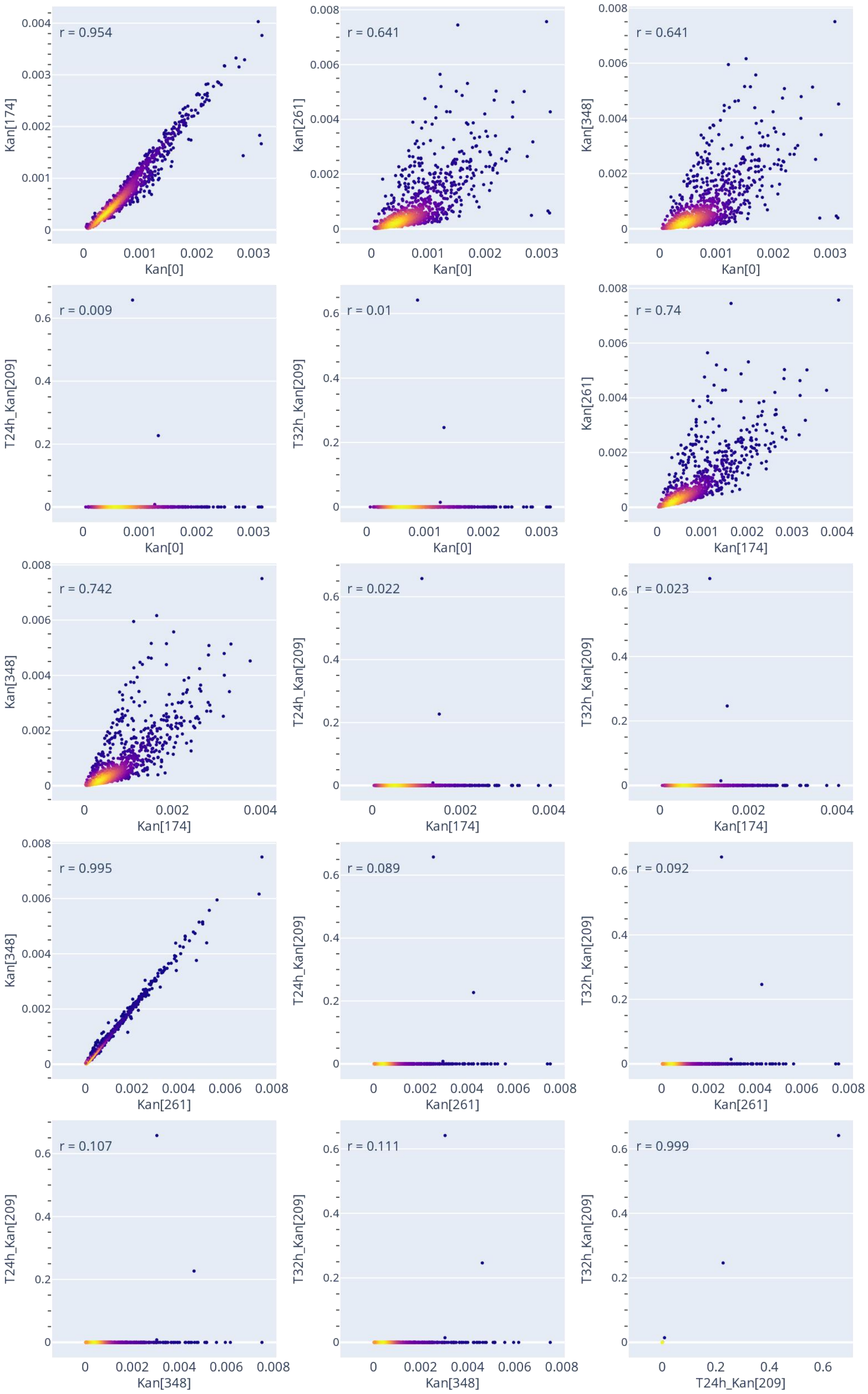
r = 0.954
r = 0.641
r = 0.641
r = 0.009
r = 0.01
r = 0.74
r = 0.742
r = 0.022
r = 0.023
r = 0.995
r = 0.089
r = 0.092
r = 0.107
r = 0.111
r = 0.999
Kan[0]
Kan[174]
Kan[261]
Kan[348]
T24h_Kan[209]
T32h_Kan[209]

#### Figure S12: Variant frequency correlation plots

Correlation plots are shown for each pair of selection conditions, with the corresponding Pearson correlation coefficient (*r*) indicated at the top of each plot. Seven hundred thirty-four (n = 734) variants present in all samples were used for correlation assessment. For improved visualization, the density of data points is indicated by a color gradient from dark blue (low density) to yellow (high density).

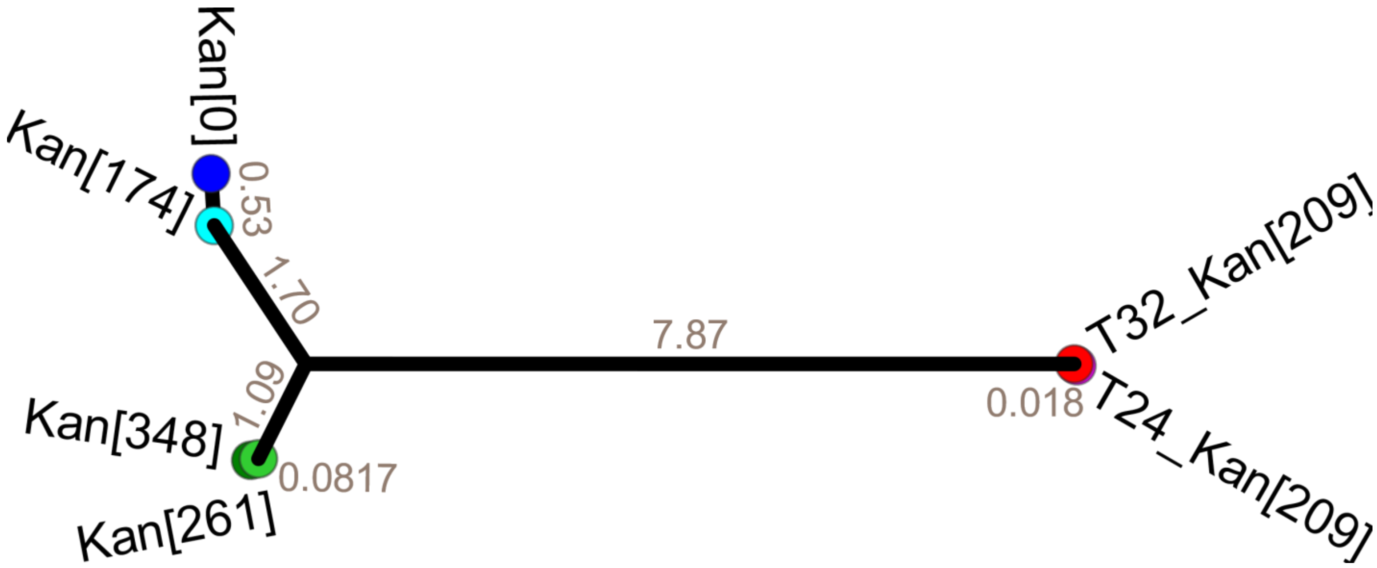

#### Figure S13: Phylogenetic tree of frequency correlation distances among selection conditions

A distance matrix was calculated by ubtracting the Pearson correlation coefficients (*r*) between each pair of conditions from 1. following the method described in the section "Comparison of samples submitted to different selection pressures".

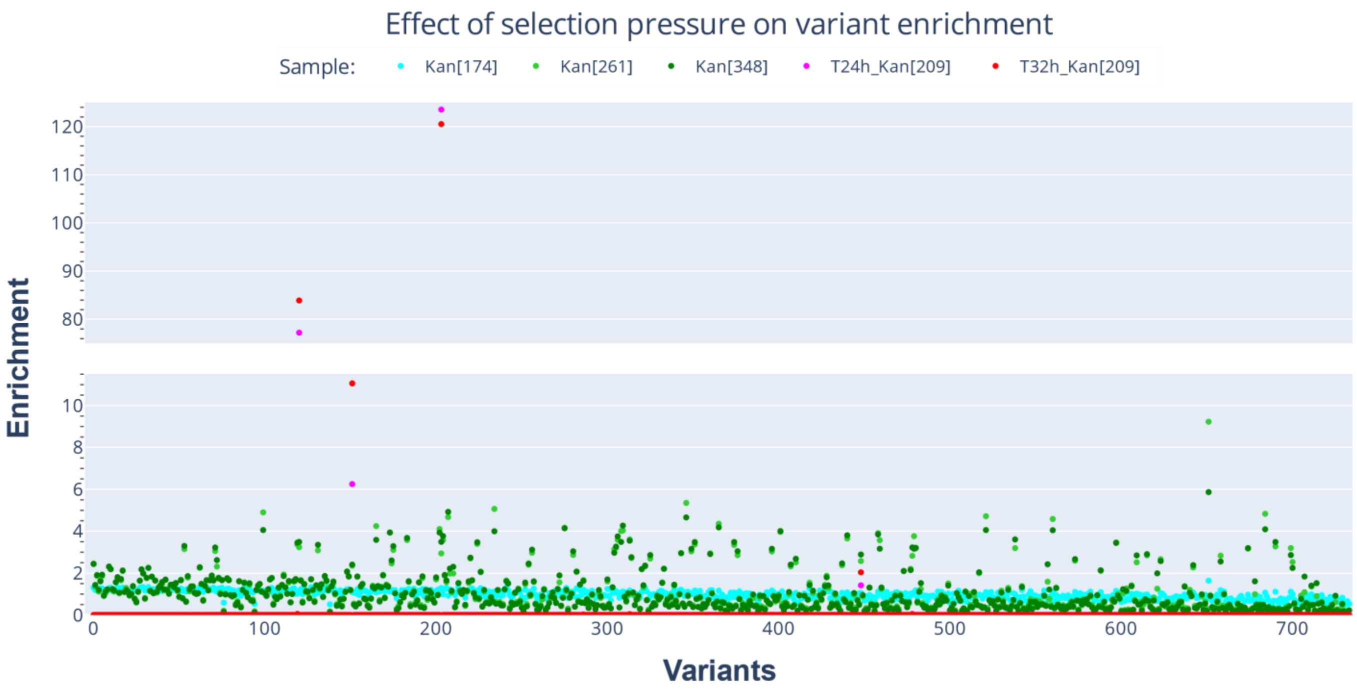

#### Figure S14: Variant enrichment in different binder selection conditions

A total of 734 variants present in all samples are displayed. Conditions: Kan[0] (no selection); Kan[174], Kan[261], Kan[348], T24h_Kan[209], and T32h_Kan[209].

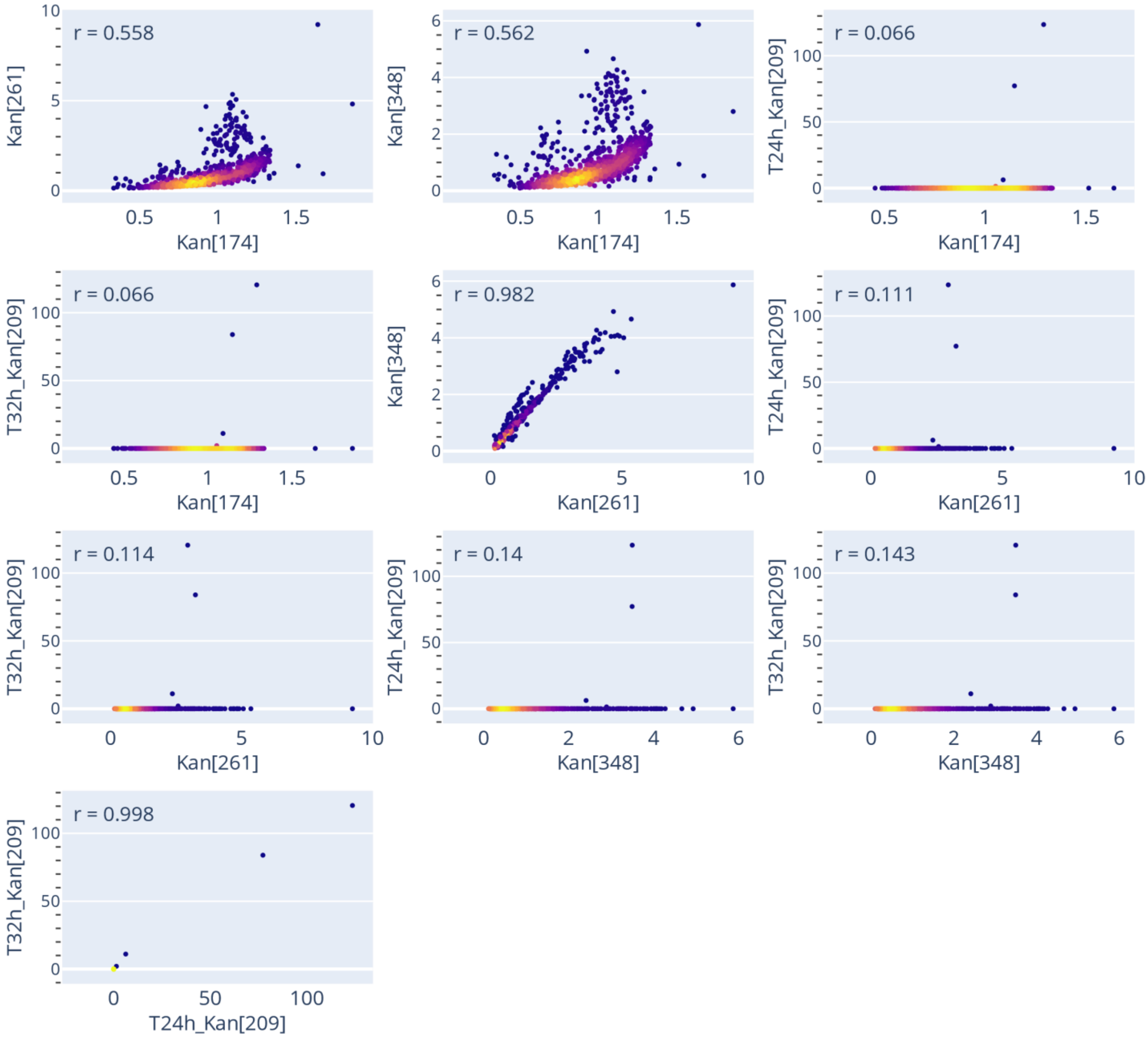

#### Figure S15: Variant enrichment correlation plots

Correlation plots are shown for each pair of selection conditions, and the corresponding Pearson correlation coefficient (*r*) is indicated on the top of each plot. For better visualization, the density of coordinate points is indicated by colors from dark blue (low density) to yellow (high density).

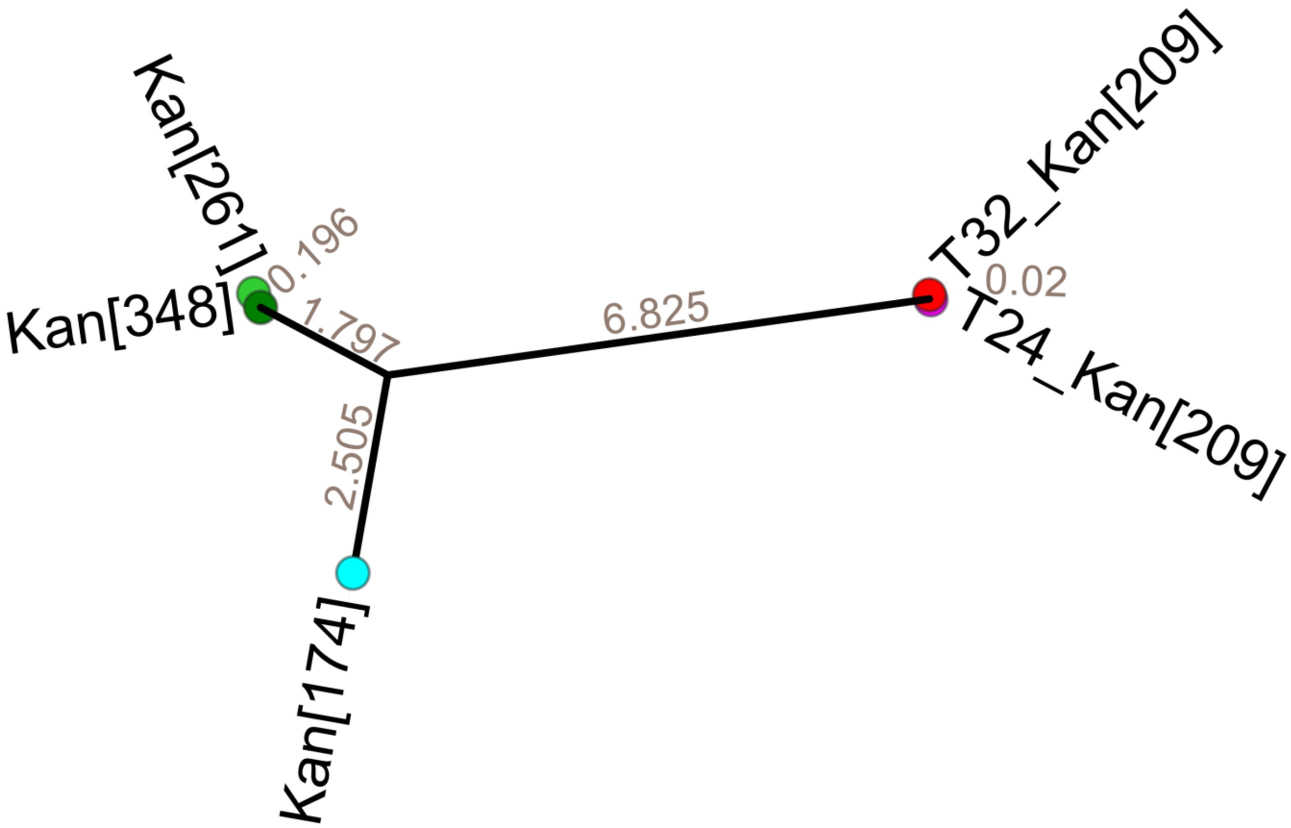

#### Figure S16: Phylogenetic tree of enrichment correlation distances among selection conditions

A distance matrix was calculated by subtracting the Pearson correlation coefficients (*r*) between each pair of conditions from 1. The distance matrix was subsequently used to construct a phylogenetic tree following the method described in the section "Comparison of samples submitted to different selection pressures".

#### Figure S17: Phylogenetic tree and frequency heatmap of variants

See supplementary file "Figure_S17_Tree+Heatmap_frequency.pdf." N-terminal extension sequences were clustered by Levenshtein distance and used to construct a dendrogram (left panel). A heatmap was generated in which cells correspond to each N-terminal sequence and linker combination, with values (reddish color intensity) representing their frequencies. Combinations of N-terminal sequence and linker that were not observed are shown in gray.

#### Figure S18: Phylogenetic tree and enrichment heatmap of variants

See supplementary file "Figure_S18_Tree+Heatmap_enrichment.pdf." N-terminal extension sequences were clustered by Levenshtein distance and used to construct a dendrogram (left panel). A heatmap was generated in which cells correspond to each N-terminal sequence and linker combination, with values (reddish color intensity) representing their enrichments. Combinations of N-terminal sequence and linker that were not observed are shown in gray.

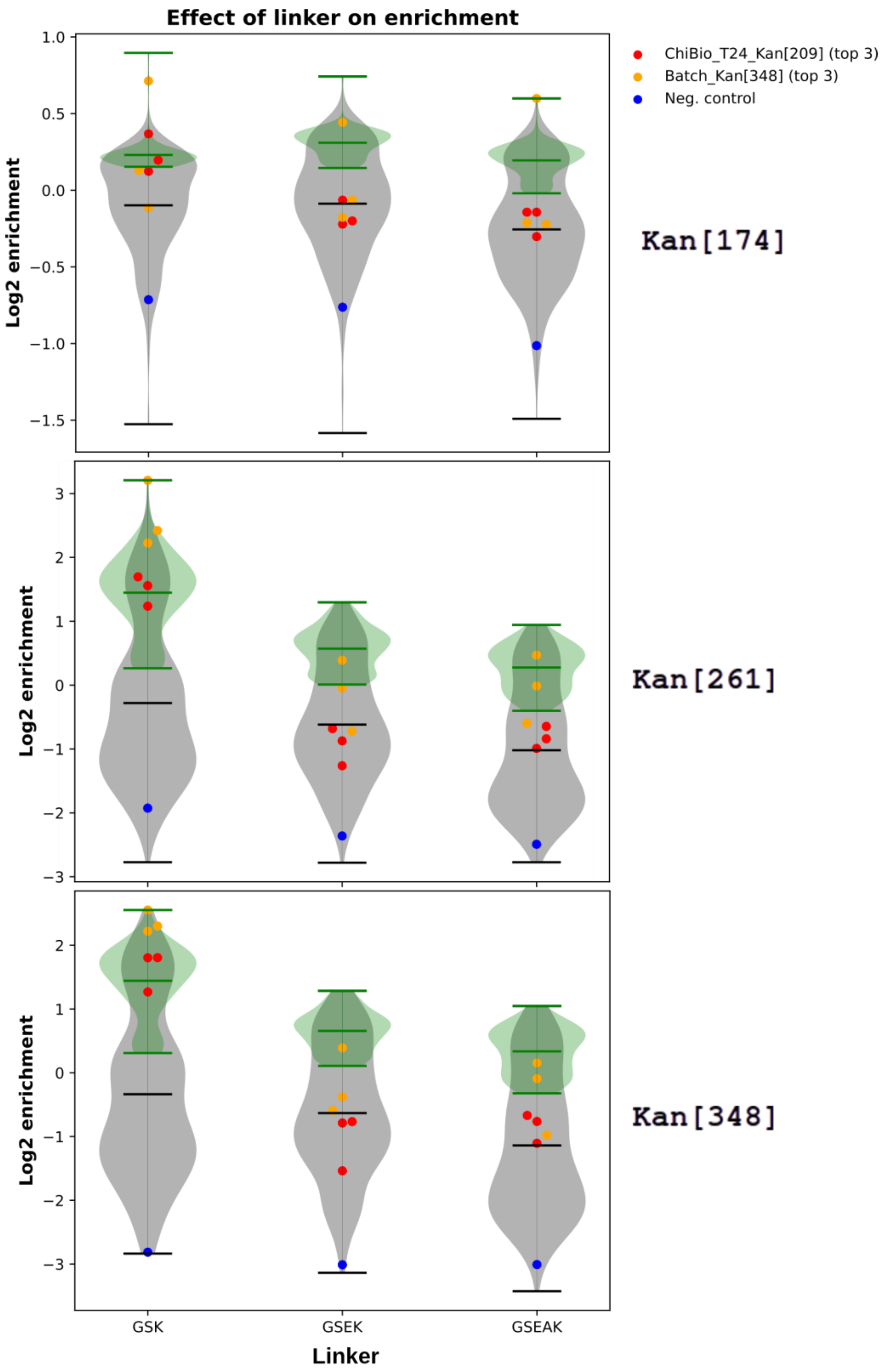

#### Figure S19: Effect of linker on enrichment during short-term (batch) selection conditions

Bacterial cultures were grown in an orbital shaker for 4–5 h in the presence of varying kanamycin concentrations: 174 µg/mL (Kan[174]), 261 µg/mL (Kan[261]), and 348 µg/mL (Kan[348]). Violin plot distributions represented in gray correspond to the entire population, while the top 100 enriched variants are represented in green.

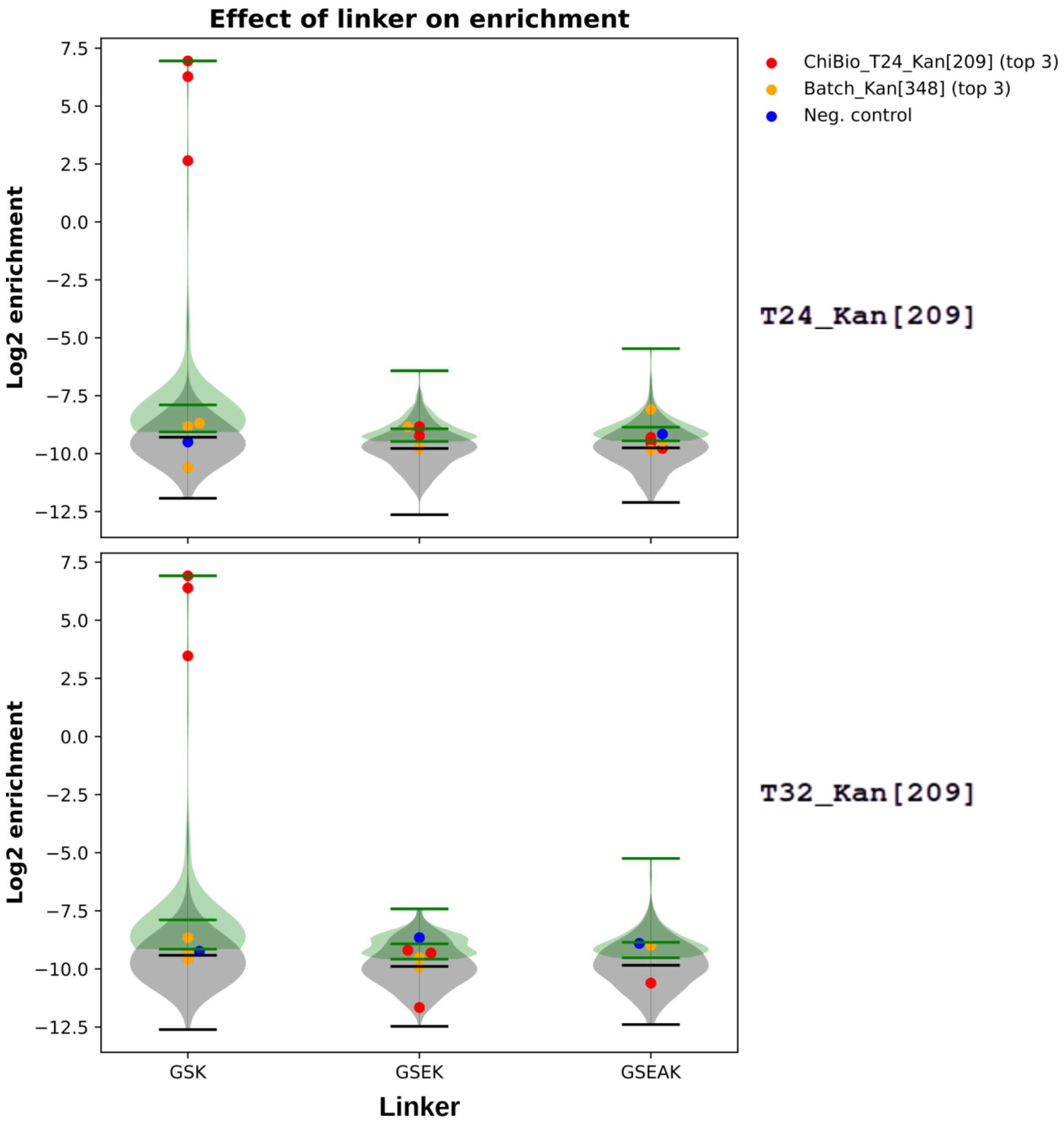

#### Figure S20: Effect of linker on variant enrichment during long-term (turbidostat) selection conditions

Bacterial cultures were grown in the presence of 209 µg/mL kanamycin for 24 h (T24_Kan[209]) or for 32 h (T32_Kan[209]). Violin plot distributions represented in gray correspond to the entire population, while the top 100 enriched variants are represented in green.

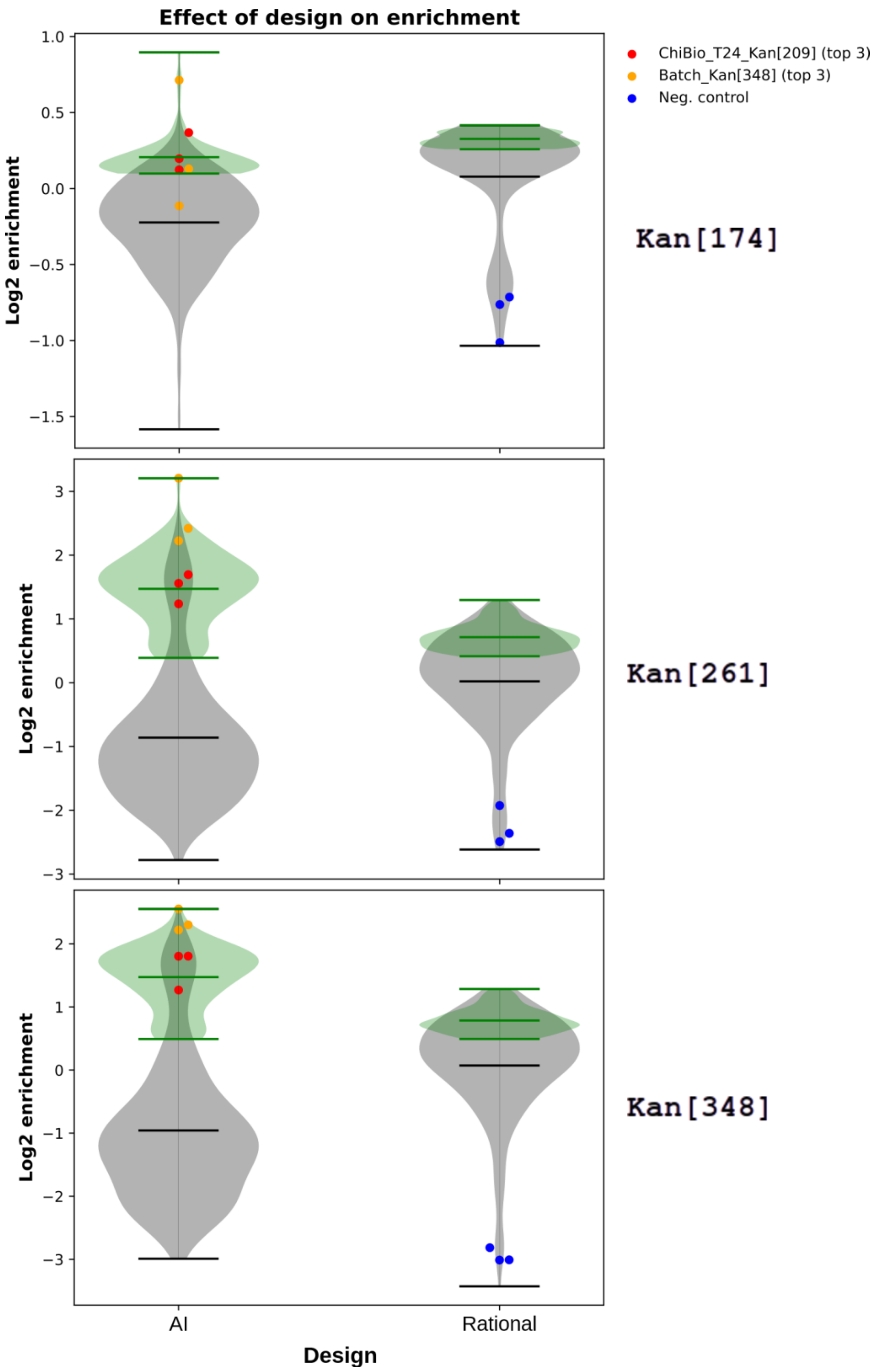

#### Figure S21: Effect of design on variant enrichment during short-term (batch) selection conditions

Bacterial cultures were grown in an orbital shaker for 4–5 h in the presence of varying kanamycin concentrations: 174 µg/mL (Kan[174]), 261 µg/mL (Kan[261]), and 348 µg/mL (Kan[348]). Violin plot distributions represented in gray correspond to the entire population, while the top 100 enriched variants are represented in green.

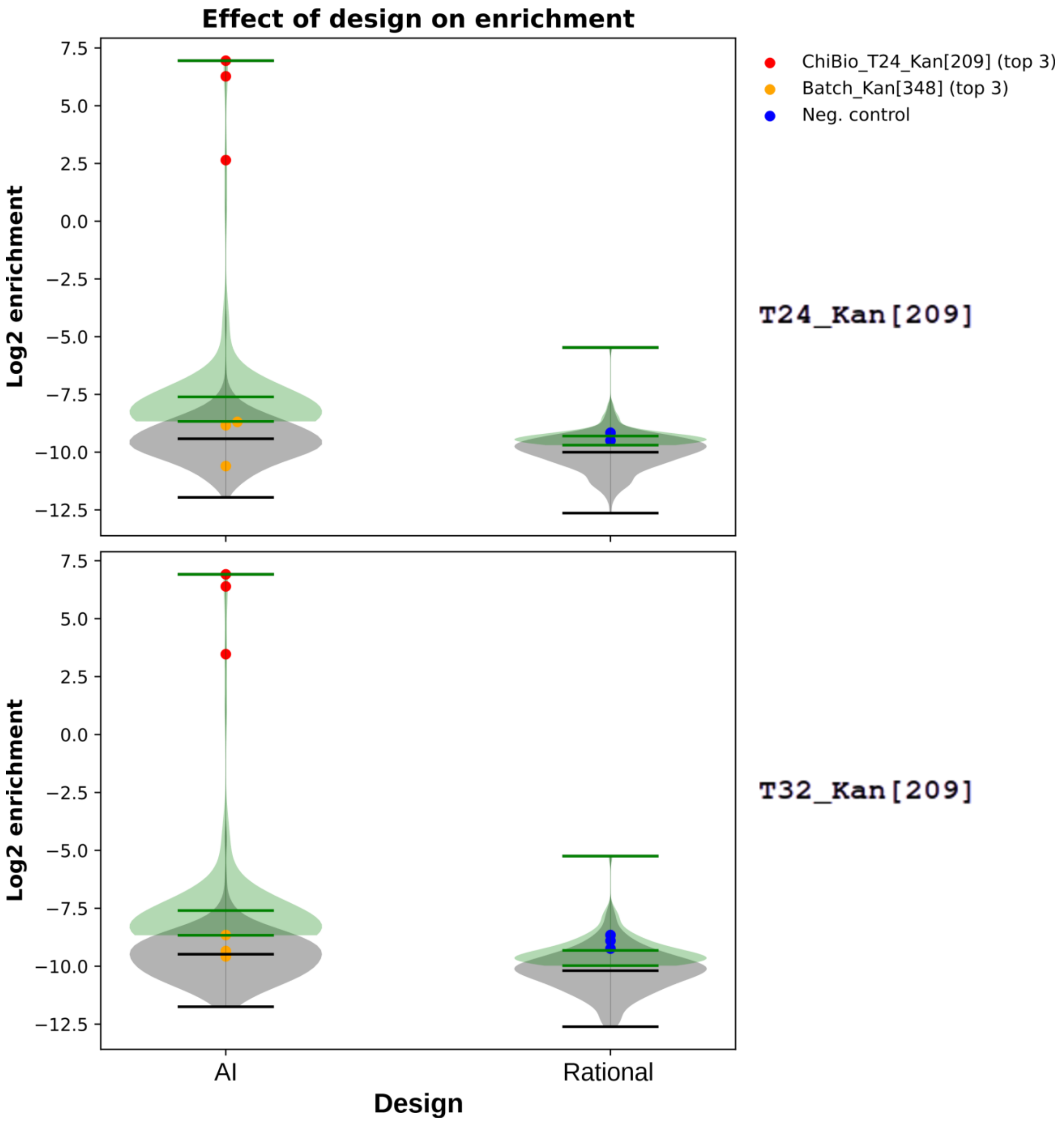

#### Figure S22: Effect of design on variant enrichment during long-term (turbidostat) selection conditions

Bacterial cultures were grown in the presence of 209 µg/mL kanamycin for 24 h (T24_Kan[209]) or for 32 h (T32_Kan[209]). Violin plot distributions represented in gray correspond to the entire population, while the top 100 enriched variants are represented in green.

A

| Name | Sequence | Affinity (µM) |
|---|---|---|
| ip4 | AGSEAKWARLARRIAGAGGVTLDGFG | **0.0024** ± 0.0004 |
| ip4mutG | AGSEAKWARLARRTAGAGGVTLNGAG | **10.6** ± 1 |
| T32_Kan[209]_top1 | AEEEAARFAALLAALPGSKWARLARRTAGAGGVTLNGAG | **0.152**± 0.025 |
| Kan[348]_top1 | SAAARWAAALAALPGSKWARLARRTAGAGGVTLNGAG | **0.235**±0.026 |
| Kan[348]_top2 | AAAERARFAALLAKLPGSKWARLARRTAGAGGVTLNGAG | **0.581**±0.098 |

B

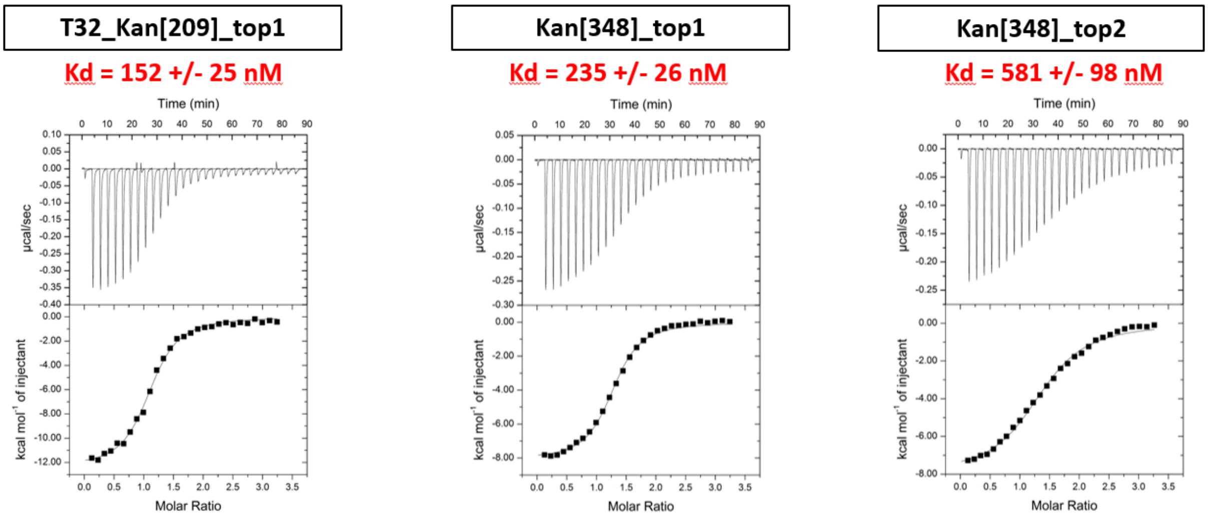

#### Figure S23: Affinity of three selected candidates with $ASF1A_N$ measured by isothermal titration calorimetry (ITC)

**(A)** Sequences of the selected binders obtained using two selection modes (batch and turbidostat) at maximum stringency. **(B)** ITC thermograms and binding curve fitting for ip4mutG and the three selected binders.

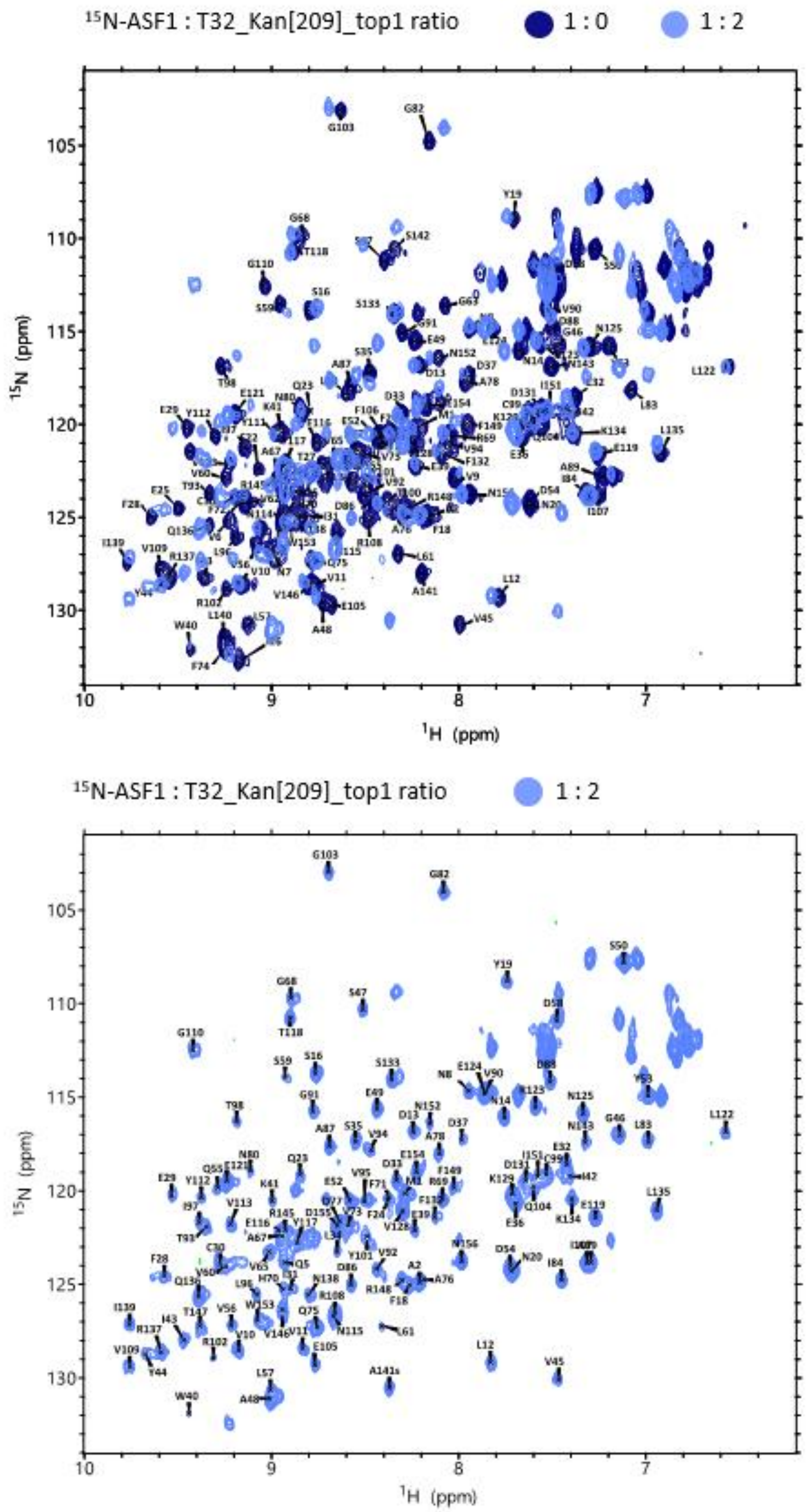

#### Figure S24: NMR spectra of $^{15}$N-labeled ASF1A$_N$ alone and in complex with T32_Kan[209]

Overlay of the $^{1}$H–$^{15}$N spectra of uniformly $^{15}$N-labeled ASF1A$_N$ alone (dark blue) and after addition of two equivalents of unlabeled T32_Kan_top1 peptide (light blue).

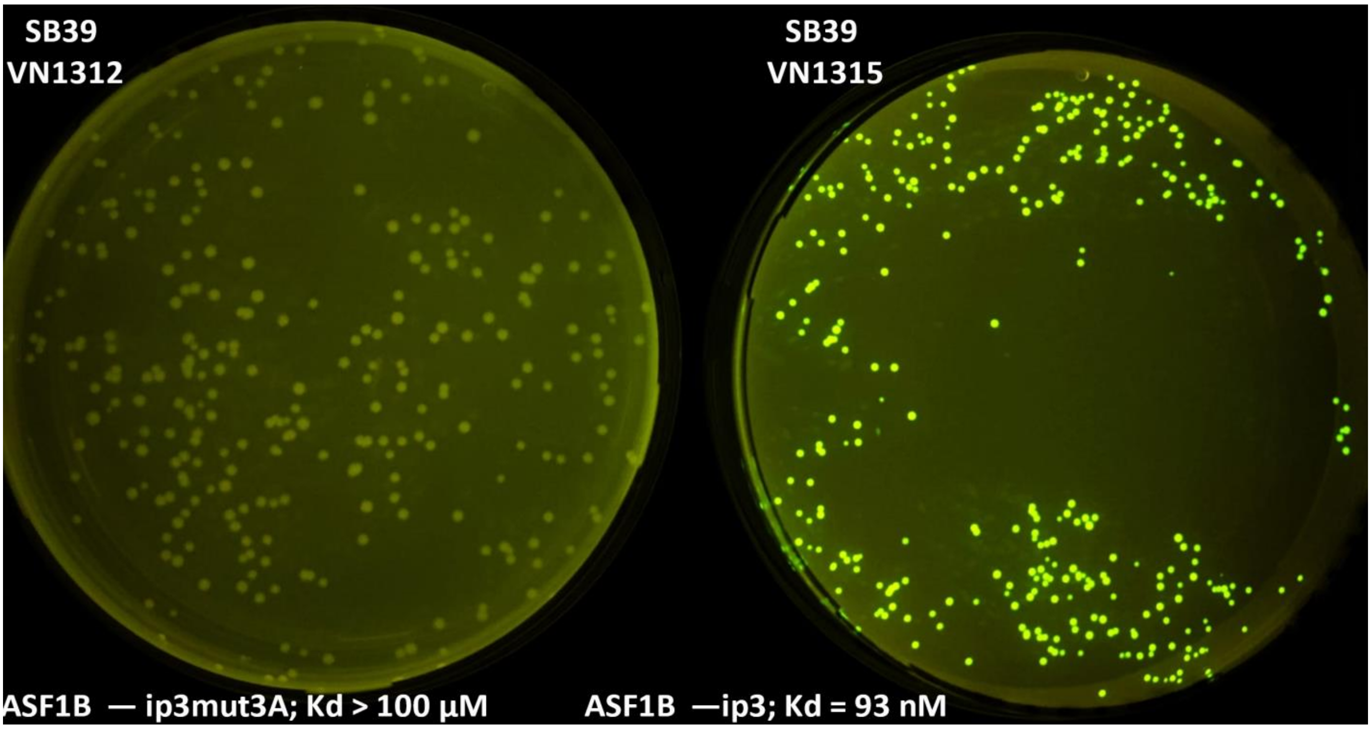

#### Figure S25: Fluorescence of bacterial colonies as a function of interaction strength

Overnight cultures of SB39 VN1312 (left, corresponding to the $ASF1B_N$–ip3mut3a interaction, $K_d > 100$ µM) and SB39 VN1315 (right, corresponding to the $ASF1B_N$–ip3, $K_d = 93$ nM) grown on LB agar plates supplemented with chloramphenicol (34 µg/mL), IPTG (200 µM), and anhydrotetracycline (200 ng/µL aTc) were visualized using a blue light transilluminator (Safe Imager$^{TM}$, Invitrogen, Cat. No. G6600EU).

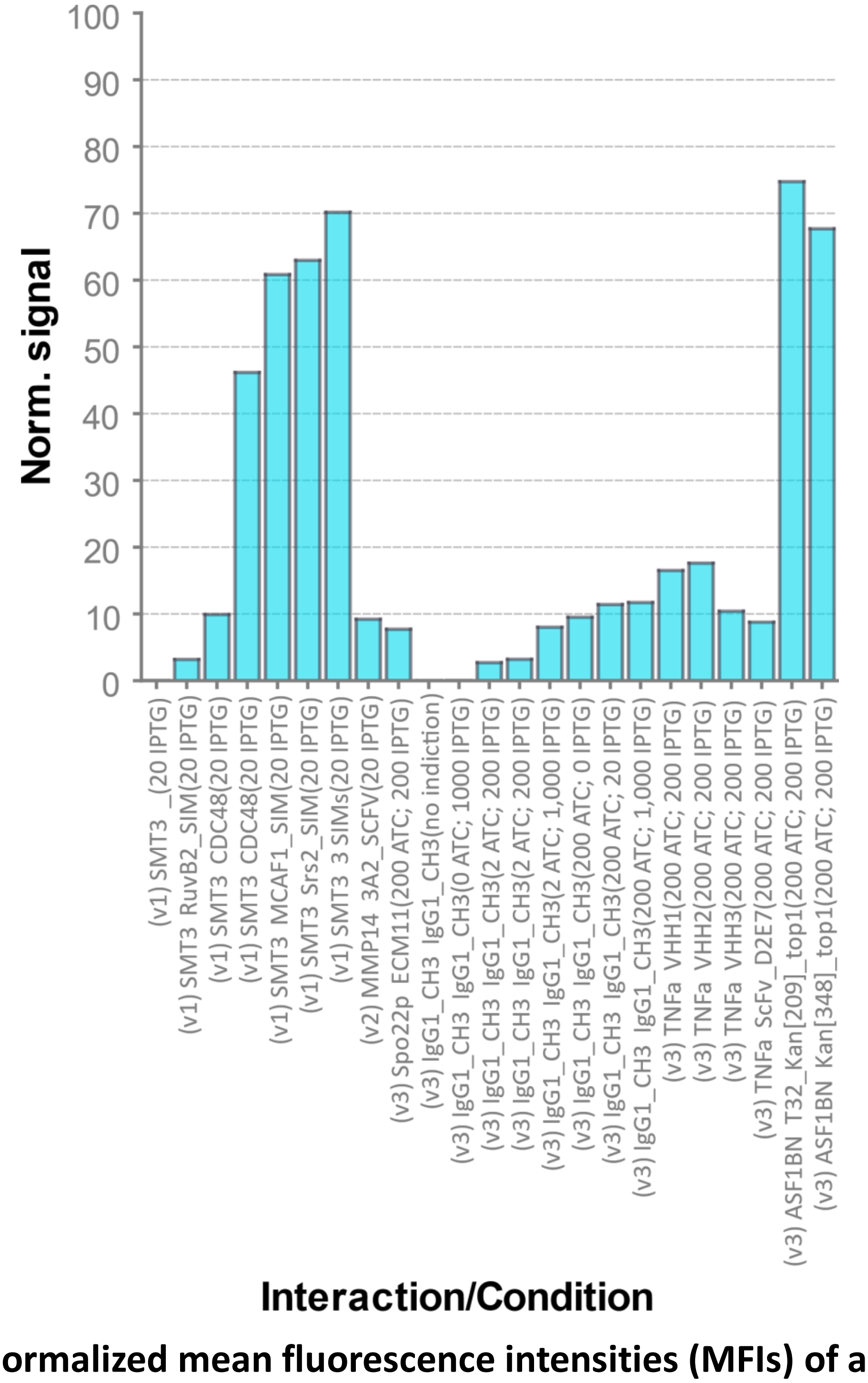

#### Figure S26: Normalized mean fluorescence intensities (MFIs) of alternative protein–protein interactions using qB2H system variants and inducing conditions

Interactions involving distinct protein domains and folding variants were assessed using qB2H systems v1, v2, and v3. The plot displays normalized interaction signals relative to the reference pairs ASF1B$_N$–ip3 and ASF1B$_N$–ip3_mut3a, as described in the main text. The x-axis labels indicate the qB2H version and the interacting partners, formatted as follows: (qB2H version) Partner_A–Partner_B (inducers). Anhydrotetracycline (aTc) concentrations are reported in ng/mL; IPTG concentrations are given in µM. All experimental conditions, acronyms, and constructs are defined in the Methods section.

### 4. References

(1) Kuznedelov, K., Minakhin, L., Niedziela-Majka, A., Dove, S. L., Rogulja, D., Nickels, B. E., Hochschild, A., Heyduk, T., and Severinov, K. (2002) A Role for Interaction of the RNA Polymerase Flap Domain with the σ Subunit in Promoter Recognition. *Science 295*, 855–857.

(2) Bakail, M., Gaubert, A., Andreani, J., Moal, G., Pinna, G., Boyarchuk, E., Gaillard, M.-C., Courbeyrette, R., Mann, C., Thuret, J.-Y., Guichard, B., Murciano, B., Richet, N., Poitou, A., Frederic, C., Le Du, M.-H., Agez, M., Roelants, C., Gurard-Levin, Z. A., Almouzni, G., Cherradi, N., Guerois, R., and Ochsenbein, F. (2019) Design on a Rational Basis of High-Affinity Peptides Inhibiting the Histone Chaperone ASF1. *Cell Chem Biol 26*, 1573-1585.e10.

(3) McLaughlin Jr, R. N., Poelwijk, F. J., Raman, A., Gosal, W. S., and Ranganathan, R. (2012) The spatial architecture of protein function and adaptation. *Nature 491*, 138–142.

(4) Rennig, M., Martinez, V., Mirzadeh, K., Dunas, F., Röjsäter, B., Daley, D. O., and Nørholm, M. H. H. (2018) TARSyn: Tunable Antibiotic Resistance Devices Enabling Bacterial Synthetic Evolution and Protein Production. *ACS Synth. Biol. 7*, 432–442.

(5) Watson, J. L., Juergens, D., Bennett, N. R., Trippe, B. L., Yim, J., Eisenach, H. E., Ahern, W., Borst, A. J., Ragotte, R. J., Milles, L. F., Wicky, B. I. M., Hanikel, N., Pellock, S. J., Courbet, A., Sheffler, W., Wang, J., Venkatesh, P., Sappington, I., Torres, S. V., Lauko, A., De Bortoli, V., Mathieu, E., Ovchinnikov, S., Barzilay, R., Jaakkola, T. S., DiMaio, F., Baek, M., and Baker, D. (2023) De novo design of protein structure and function with RFdiffusion. *Nature 620*, 1089–1100.

(6) McLachlan, A. D. (1982) Rapid comparison of protein structures. *Acta Cryst A 38*, 871–873.

(7) Dauparas, J., Anishchenko, I., Bennett, N., Bai, H., Ragotte, R. J., Milles, L. F., Wicky, B. I. M., Courbet, A., de Haas, R. J., Bethel, N., Leung, P. J. Y., Huddy, T. F., Pellock, S., Tischer, D., Chan, F., Koepnick, B., Nguyen, H., Kang, A., Sankaran, B., Bera, A. K., King, N. P., and Baker, D. (2022) Robust deep learning-based protein sequence design using ProteinMPNN. *Science 378*, 49–56.

(8) Evans, R., O'Neill, M., Pritzel, A., Antropova, N., Senior, A., Green, T., Žídek, A., Bates, R., Blackwell, S., Yim, J., Ronneberger, O., Bodenstein, S., Zielinski, M., Bridgland, A., Potapenko, A., Cowie, A., Tunyasuvunakool, K., Jain, R., Clancy, E., Kohli, P., Jumper, J., and Hassabis, D. (2022) Protein complex prediction with AlphaFold-Multimer. *bioRxiv* 2021.10.04.463034.

(9) Mirdita, M., Schütze, K., Moriwaki, Y., Heo, L., Ovchinnikov, S., and Steinegger, M. (2022) ColabFold: making protein folding accessible to all. *Nat Methods 19*, 679–682.

(10) Tu, Q., Yin, J., Fu, J., Herrmann, J., Li, Y., Yin, Y., Stewart, A. F., Müller, R., and Zhang, Y. (2016) Room temperature electrocompetent bacterial cells improve DNA transformation and recombineering efficiency. *Sci Rep 6*, 24648.

(11) Steel, H., Habgood, R., Kelly, C., and Papachristodoulou, A. Chi.Bio: An open-source automated experimental platform for biological science research. *bioRxiv*.

(12) Steel, H., Habgood, R., Kelly, C. L., and Papachristodoulou, A. (2020) In situ characterisation and manipulation of biological systems with Chi.Bio. *PLoS Biol 18*, e3000794.

(13) Zhang, J., Kobert, K., Flouri, T., and Stamatakis, A. (2014) PEAR: a fast and accurate Illumina Paired-End reAd mergeR. *Bioinformatics 30*, 614–620.
